# Cerebellar microRNA-206 tunes Purkinje neuron firing dynamics to control sensorimotor gating

**DOI:** 10.64898/2026.06.29.734826

**Authors:** Mary P. Heyer, Masago Ishikawa, Junshi Wang, J. Erol Evangelista, Amanda K. Fakira, Avi Ma’ayan, Guoping Feng, Paul J. Kenny

**Affiliations:** Nash Family Department of Neuroscience, Icahn School of Medicine at Mount Sinai, New York, NY 10029, USA; Nathan S. Kline Institute for Psychiatric Research, Orangeburg, NY 10962, USA; Department of Pharmacological Sciences, Mount Sinai Center for Bioinformatics, Icahn School of Medicine at Mount Sinai, New York, NY 10029, USA; Department of Biomedical Sciences, Cooper Medical School of Rowan University, Camden, NJ 08103, USA; McGovern Institute for Brain Research, Department of Brain and Cognitive Sciences, Massachusetts Institute of Technology, Cambridge, MA 02139, USA; Stanley Center for Psychiatric Research, Broad Institute of MIT and Harvard, Cambridge, MA 02142, USA

## Abstract

MicroRNAs are potent regulators of gene expression in the brain, yet the cellular mechanisms through which they shape neuronal activity and behavior remain poorly understood. MicroRNA-206 (miR-206) has been genetically and transcriptionally linked to schizophrenia and other neuropsychiatric disorders, but its functions in the nervous system are largely unknown. Here we show that miR-206 expression in the brain is restricted to postnatal cerebellar Purkinje cells (PCs). miR-206 was dispensable for PC cell fate specification, dendritic morphogenesis, and cerebellar-regulated motor coordination. Transcriptional profiling with single-nucleus and spatial resolution, integrated with Ago2-associated miRNA-target repression mapping (HITS-CLIP) and ribosome-associated RNA profiling (TRAP-seq), showed that miR-206 regulates translational programs in PCs controlling neuronal excitability. Accordingly, miR-206 deficiency increased the tonic firing of PCs, whereas elevating miR-206 expression shifted PCs from tonic to high-frequency burst firing. Constitutive or PC-specific deletion of miR-206 impaired prepulse inhibition (PPI) of the acoustic startle response, a conserved form of sensorimotor gating disrupted in schizophrenia and related disorders. Restoration of miR-206 expression in PCs rescued PPI deficits in miR-206-deficient mice, while elevating miR-206 expression in PCs impaired PPI in wild-type animals. Together, these findings reveal that a schizophrenia-linked microRNA tunes Purkinje neuron firing dynamics to control sensorimotor gating.

## INTRODUCTION

Neuropsychiatric disorders are highly heritable yet clinically heterogeneous, due to complex interactions of genetic variants and epigenetic factors^1,2^. MicroRNAs (miRNAs) are small, noncoding RNAs that act as potent post-transcriptional epigenetic regulators^3^. miRNAs bind to complementary sequences in the 3’ untranslated region (3’UTR) of gene transcripts to facilitate their recruitment to the RNA-induced silencing complex (RISC), where they undergo translational repression, transcript destabilization, or degradation^4,5^. miRNAs are evolutionarily conserved, enriched in the brain, and regulate neuronal development, activity, plasticity, and survival^6–8^. miRNA and target gene variants, as well as dysregulated miRNA expression, are increasingly implicated in psychiatric and neurodevelopmental disorders including schizophrenia, bipolar disorder, major depressive disorder (MDD), and autism spectrum disorder (ASD)^9–17^. However, the molecular, cellular, and circuit-level mechanisms through which disease-associated miRNAs influence brain function remain largely unknown.

The conserved microRNA miR-206 has been strongly linked with schizophrenia and other neuropsychiatric disorders. Predicted miR-206 binding sites are highly enriched among genetic loci associated with schizophrenia risk^18^, while a schizophrenia protective allele identified by genome-wide association studies disrupts miR-206-mediated repression of *NT5C2*^19^. Common variants in the *MIR206/MIR133b* locus are associated with schizophrenia and ASD^20,21^, and *MIR206* polymorphisms are also associated with susceptibility to bipolar disorder and treatment response in patients^22^. miR-206 is among the most dysregulated miRNAs in blood exosomes of schizophrenia patients^23^, and is similarly elevated in serum of patients with cognitive impairment^24^, a core feature of schizophrenia and related disorders. These findings suggest that miR-206 may influence conserved mechanisms across neuropsychiatric domains, yet the brain regions, neuronal populations, and physiological functions through which miR-206 acts are currently undefined.

miR-206 expression has been detected in mammalian brain regions including the prefrontal cortex (PFC) and hippocampus^25–27^, and has also been found to be particularly enriched in cerebellar Purkinje cells (PCs)^28,29^. Although traditionally associated with motor coordination, the cerebellum is increasingly recognized as an important regulator of cognitive, affective, and sensorimotor processes implicated in neuropsychiatric disorders^30–37^. Structural and functional cerebellar abnormalities, including altered Purkinje cell (PC) organization and connectivity, have been reported in schizophrenia, bipolar disorder, and ASD^38–44^. Cerebellar dysfunction has also been implicated in abnormalities in prepulse inhibition (PPI) and other forms of sensorimotor gating^45–50^, which are considered translational endophenotypes of schizophrenia-spectrum disorders^51–54^.

Here, using high-resolution transcriptional and translational mapping and precise genetic targeting in mice, we show that brain-expressed miR-206 is restricted to postnatal cerebellar PCs, where it is dispensable for PC specification, dendritic morphogenesis, and survival. Instead, miR-206 acts in cerebellar PCs to regulate gene programs controlling neuronal excitability. Accordingly, miR-206 bidirectionally regulates PC firing dynamics, with corresponding bidirectional effects on PPI of the auditory startle response. These findings establish that a schizophrenia-linked microRNA regulates the intrinsic firing state of cerebellar Purkinje neurons to control sensorimotor gating.

## RESULTS

### miR-206 in mammalian brain is expressed exclusively in cerebellar Purkinje cells

miR-206 is reported to be densely expressed in cerebellar PCs^29,28^, and across other brain regions in postnatal rodent brain, including prefrontal cortex (PFC), hippocampus, amygdala, hypothalamus, and ventral midbrain^25–27^. However, the short antisense probes used to detect mature miR-206 potentially yield non-specific signals by binding to its paralog miR-1 or homologous regions within other RNAs. Therefore, to determine the precise expression patterns of miR-206 in brain we designed a custom probe for a region of the miR-206 precursor previously validated by northern blot^55^. RNAscope *in situ* hybridization in adult mouse brain detected *pre-miR-206* exclusively in cerebellar PCs (Figures 1A–C), identified by co-expression of the vesicular GABA transporter gene *Slc32a1* (Vgat), a marker of GABAergic inhibitory neurons including PCs, location in the PC layer, and large cell size. This was confirmed by colocalization of *pre-miR-206* with cells expressing the PC marker *Calb1* (Figure 1D). *Pre-miR-206* signal was absent in all other brain regions examined, including mPFC (Figure S1A), hippocampus (Figure S1B), and ventral midbrain (Figure S1C). Quantitative RT-PCR (qPCR) of adult brain regions confirmed that expression of mature miR-206 was restricted to the cerebellum (Figure 1E).

**Figure 1.**
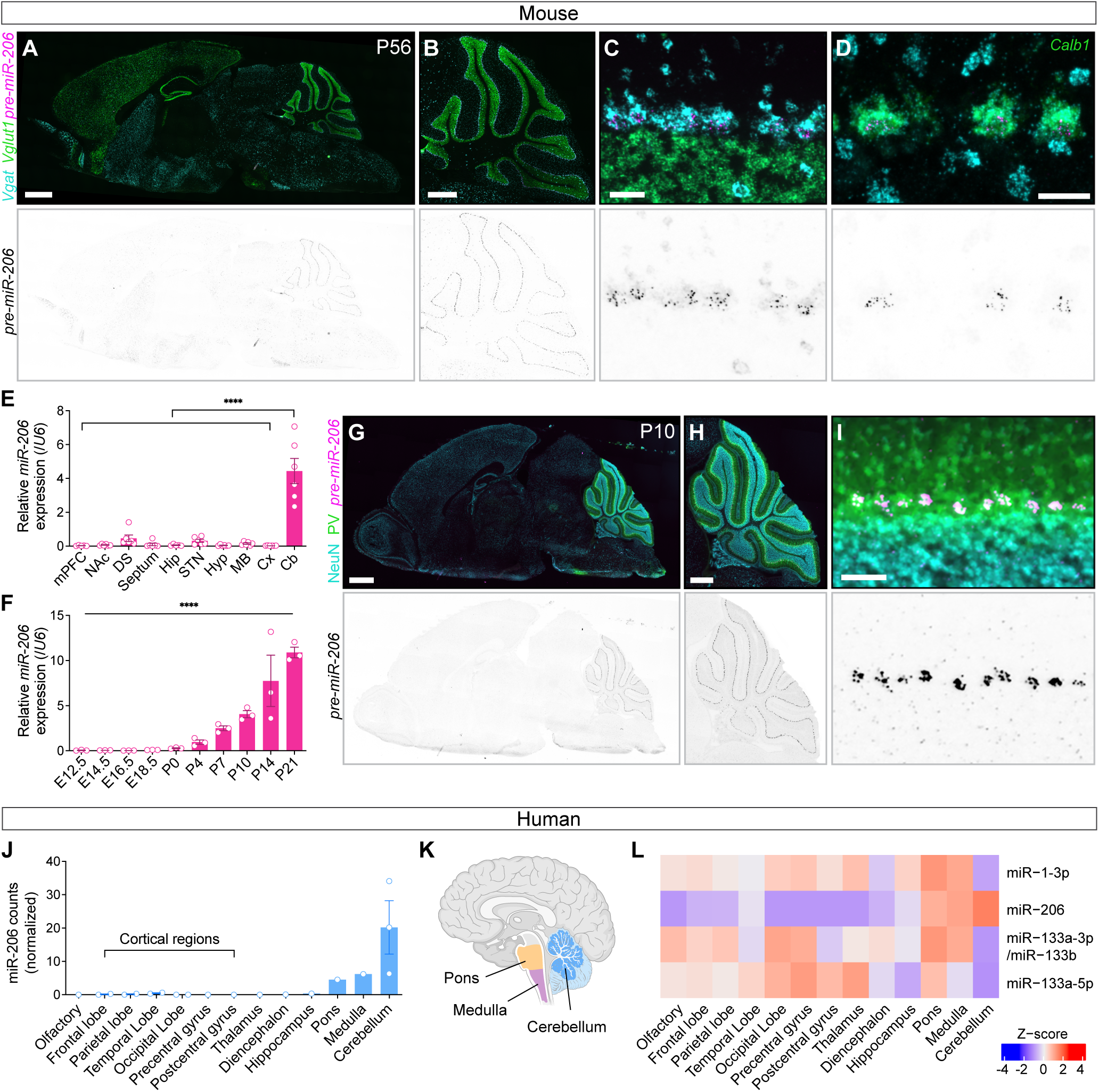
miR-206 in the brain is expressed exclusively in postnatal cerebellar Purkinje cells. **(A–D)** RNAscope fluorescence signal for miR-206 precursor (magenta, upper panels; black, lower panels), *Vgat* (cyan, upper panels), and *Vglut1* (A–C) or *Calb1* (D) (green, upper panels) transcript markers of GABAergic and glutamatergic neurons, or Purkinje cells (PCs), respectively, in postnatal day 56 male mouse brain. *pre-miR-*206 puncta are localized to PCs in cerebellum, while more diffuse, non-punctate signal is due to background and *Vgat* signal bleedthrough. Brain- and cerebellum-wide images were stitched from multiple tiled epifluorescence images. **(A)** Medial sagittal section of brain. Scale bar: 1 mm. **(B)** Cerebellar region. Scale bar: 500 µm. **(C)** *pre-miR-206* detection in PC layer. Scale bar: 50 µm. (B) and (C), inset from (A). **(D)** *pre-miR-206* colocalizes with *Calb1* in PCs. Orthogonal projection of confocal z-stack. Scale bar: 20 µm. **(E)** Quantitative RT-PCR shows enrichment of mature miR-206 in cerebellum (Cb) but not other brain regions (mPFC, medial prefrontal cortex; NAc, nucleus accumbens; DS, dorsal striatum; Septum; Hip, hippocampus; STN, subthalamic nucleus; Hyp, hypothalamus; MB, ventral midbrain; Cx, cerebral cortex). n = 6 two-month-old male mice. ***p < 0.0001, one-way ANOVA. **(F)** Quantitative RT-PCR detects increasing mature miR-206 expression in whole brain during postnatal but not embryonic development. n = 3 mice per age. ***p < 0.0001, one-way ANOVA. **(G–I)** RNAscope detection of *pre-miR-206* (magenta, upper panels; black, lower panels) and immunofluorescence detection of NeuN (cyan, upper panels) and parvalbumin (PV, green, upper panels) in postnatal day 10 brain. **(G)** Medial sagittal section of brain. Scale bar: 1 mm. **(H)** Cerebellar region. Scale bar: 500 µm. **(I)** *pre-miR-206* detection in PC layer. Scale bar: 50 µm. (H) and (I), inset from (G). **(J)** Normalized miR-206 counts across human brain regions. **(K)** Schematic of anatomical proximity of human pons and medulla to cerebellum. **(L)** Heat map of relative sequencing counts of miR-206 paralog miR-1 and downstream miR-133 across human brain regions. Results in (J–L) were generated by analysis of previously published raw bulk small RNA sequencing data^60^. Error bars indicate SEM.

miRNAs play key roles in developmental timing and cell fate determination^56,57^. To test whether miR-206 is expressed in the cerebellum and other brain regions during development, we performed miR-206 qPCR on E12.5 to P21 brain samples. miR-206 was not detected in brain before birth, but its expression rapidly increased during the first three postnatal weeks (Figure 1F). To determine whether miR-206 is similarly restricted to PCs during early postnatal development, *pre-miR-206* expression was analyzed by RNAscope in P10 brain. *pre-miR-206* signal was detected exclusively in cerebellar PCs, identified by colocalization with parvalbumin (PV), which is expressed by PCs, location in the PC layer, and large cell size (Figures 1G–I).

Altered miR-206 expression has been observed in extra-cerebellar brain regions in adult animals, including PFC and hippocampus, after social defeat stress and other stressful stimuli^27,58,59^. NMDA receptor channel blockade was also reported to modify miR-206 expression in hippocampus^25^. This could reflect *de novo* induction of miR-206 expression in brain regions that do not otherwise express this miRNA in response to stress and other perturbations. However, we found that miR-206 expression remained undetectable across all profiled non-cerebellar brain regions in adult mice after acute and chronic restraint stress, chronic forced-swim stress, and chronic social defeat stress (CSDS) (Figures S1E–H). Likewise, NMDA receptor channel blockade with the psychotomimetic dizocilpine (MK-801) had no effects on the brain-wide expression patterns of miR-206 (Figure S1I), although miR-206 expression was reduced in the cerebellum.

Finally, to confirm that miR-206 expression in the human brain is similarly restricted to the cerebellum, we analyzed publicly available small RNA sequencing data from adult human brain regions^60^. We found that miR-206 is highly expressed in human cerebellum, with lower sequencing counts in hindbrain regions, possibly reflecting contamination with adjacent cerebellar tissue (Figures 1J–L). However, miR-206 expression was absent in all sequenced forebrain and midbrain regions (Figures 1J–L). In contrast, miR-1, a paralog of miR-206, and downstream miR-133 were only detected in extra-cerebellar regions but absent in the cerebellum (Figure 1L). Together, these data establish a conserved pattern of miR-206 enrichment in postnatal cerebellum.

### miR-206 is dispensable for Purkinje cell fate specification, morphogenesis, and survival

miRNA signaling is essential for PC development, maturation, and survival^61–64,28^. Knockdown of miR-206 in cerebellar PCs, achieved using virus-mediated expression of an antisense ‘sponge’ containing high-affinity response elements to sequester miR-206, was reported to severely disrupt PC proximal dendritic branching and their synaptic innervation by cerebellar climbing fibers^28^. However, miRNA sponge approaches rely on seed-sequence recognition domains and can potentially sequester related miRNAs with similar seed sequences. Therefore, to precisely define the role of miR-206 in PC development and function, we generated constitutive and conditional *miR-206* knockout (KO) mice by homologous recombination in embryonic stem cells (Figures 2A–D). We confirmed that miR-206 expression was ablated in KO mice (Figure 2C). No changes in the expression of mature or precursor levels of miR-1 were detected in the cerebellum or across any other profiled brain region of miR-206 KO mice (Figures S2A–C). This suggests that compensatory alterations in miR-1 expression are unlikely to occur in miR-206-deficient mice.

**Figure 2.**
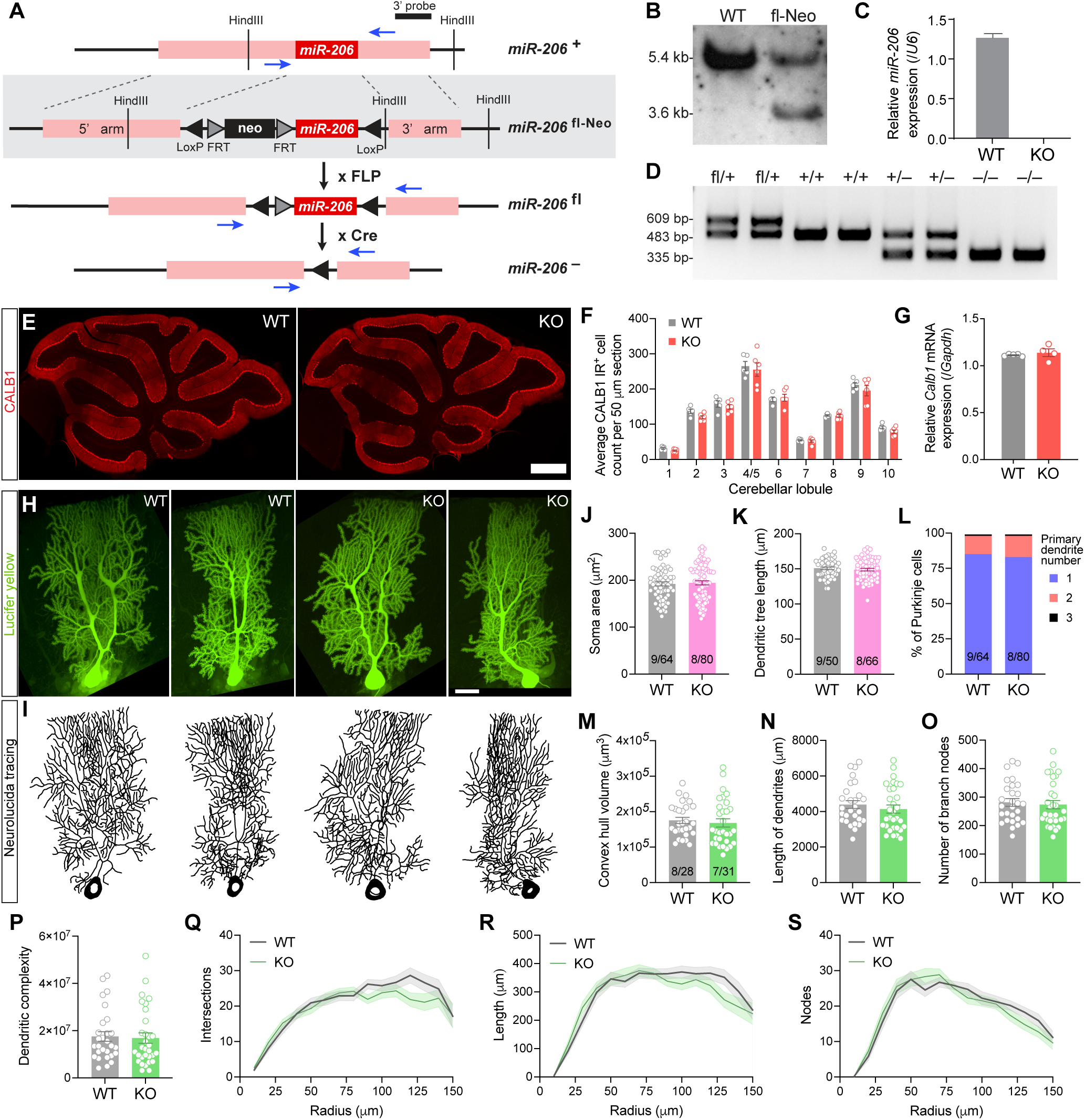
miR-206 is dispensable for Purkinje cell maturation and morphology. **(A)** Schematic of miR-206 gene targeting strategy. Mice produced from correctly targeted embryonic stem cells were bred to germline *Flp* and *Cre* driver lines to generate miR-206 floxed and knockout alleles, respectively. tk, thymidine kinase; neo, neomycin resistance cassette. Blue arrows indicate locations of PCR primers for genotyping. **(B)** Southern blot following HindIII digestion of WT and correctly targeted embryonic stem cell genomic DNA. The 3’ probe binding site is indicated in (A). **(C)** Quantitative RT-PCR demonstrates loss of mature miR-206 expression in KO mouse brain. n = 2 WT, 3 KO mice. *p = 0.0273, Welch’s t-test. **(D)** PCR genotyping of miR-206 floxed, wildtype, and null alleles from mouse genomic DNA. **(E)** Immunofluorescence staining for calbindin 1 (CALB1), a PC marker, in representative WT and KO sagittal sections of adult cerebellar vermis. Scale bar: 500 µm. **(F)** miR-206 deletion does not alter the number of CALB1 immunoreactive PCs per lobule in cerebellar vermis sagittal sections. n = 5 WT, 6 KO male mice. Counts were summed from six 50-µm sections per animal. **(G)** Quantitative RT-PCR shows no change in mRNA expression of *Calb1* in KO cerebellum. n = 5 WT, 4 KO males. **(H)** Representative orthogonal projection confocal images of WT and KO PCs in lobule VI vermis iontophoretically filled with Lucifer Yellow. Scale bar: 20 µm. **(I)** Neurolucida360 skeleton tracings of neurons in (H). **(J)** PC soma cross-section area, **(K)** PC length, and **(L)** percentage of PCs with one, two, or three primary dendrites were not altered in KO mice. (J) and (K): n = 64 WT, 80 KO neurons from 9 WT, 8 KO male mice; (L): n = 50 WT, 66 KO neurons from 8 WT, 7 KO male mice. **(M)** Envelope (convex hull) volume, **(N)** total length of dendrites, **(O)** total number of branch nodes, and **(P)** dendritic complexity are similar between WT and KO PC tracings. **(Q–S)** Sholl analyses of tracings show no significant differences in branch intersections (Q), length (R), or nodes (S) of PC dendrites of WT and KO mice. (M–S): n = 28 WT, 31 KO neurons from 8 WT, 7 KO male mice. Error bars and bands indicate SEM.

Using calbindin 1 (CALB1) immunoreactivity to detect PCs, we found that numbers of cerebellar PCs were unchanged between miR-206 KO and WT mice across all lobules within cerebellar vermis (Figures 2E and 2F). Global *Calb1* gene expression was also unaffected after miR-206 deletion when assessed using qPCR of bulk cerebellar tissue (Figure 2G). RNAscope profiling similarly confirmed that *Calb1* transcript levels in PCs across cerebellar vermis were equivalent in WT and KO (Figures S2D–F).

To determine whether miR-206 deficiency altered PC morphology, PCs from WT and KO mice were iontophoretically filled with Lucifer Yellow dye and PC dendritic trees reconstructed (Figures 2H and 2I). No differences in soma area (Figure 2J), tree length (Figure 2K), or number of primary dendrites (Figure 2L) were detected between groups. Measures of overall dendritic arbor, including convex hull volume, length of dendrites, number of branch nodes, and dendritic complexity were also unaffected (Figures 2M–P). Sholl analysis^65^ revealed no change in PC dendritic branch intersections, lengths, or nodes in KO mice (Figures 2Q–S). These findings demonstrate that miR-206 is dispensable for PC cell fate specification, differentiation, and morphogenesis.

### Transcriptional and spatial profiling of miR-206-deficient cerebellum

To define the cellular and transcriptional consequences of miR-206 deficiency across the cerebellum at higher resolution, we performed single-nucleus RNA sequencing (snRNA-seq) of cerebella from WT and miR-206 KO mice. Unsupervised graph-based clustering identified transcriptionally distinct cell populations (Figures 3A–C). This revealed no obvious genotype-dependent alterations in cluster structure or cellular composition in the KO animals. The expression of canonical cell type marker genes was similar in KO mice relative to controls (Figure S3A), enabling high-confidence cell type annotation using a reference dataset^66^. Total numbers and relative proportions of each cerebellar cell type including PCs were unchanged between WT and KO mice (Figure 3B). PCs were subclustered into Aldoc and anti-Aldoc subtypes using the same reference dataset (Figures 3E and S3B), which showed that total numbers and marker gene expression levels for these PC subpopulations were also unchanged (Figures 3F and S3C). This was further confirmed by RNAscope detection of *Aldoc* expression in the cerebellum of WT and KO mice (Figures S3D–F).

**Figure 3.**
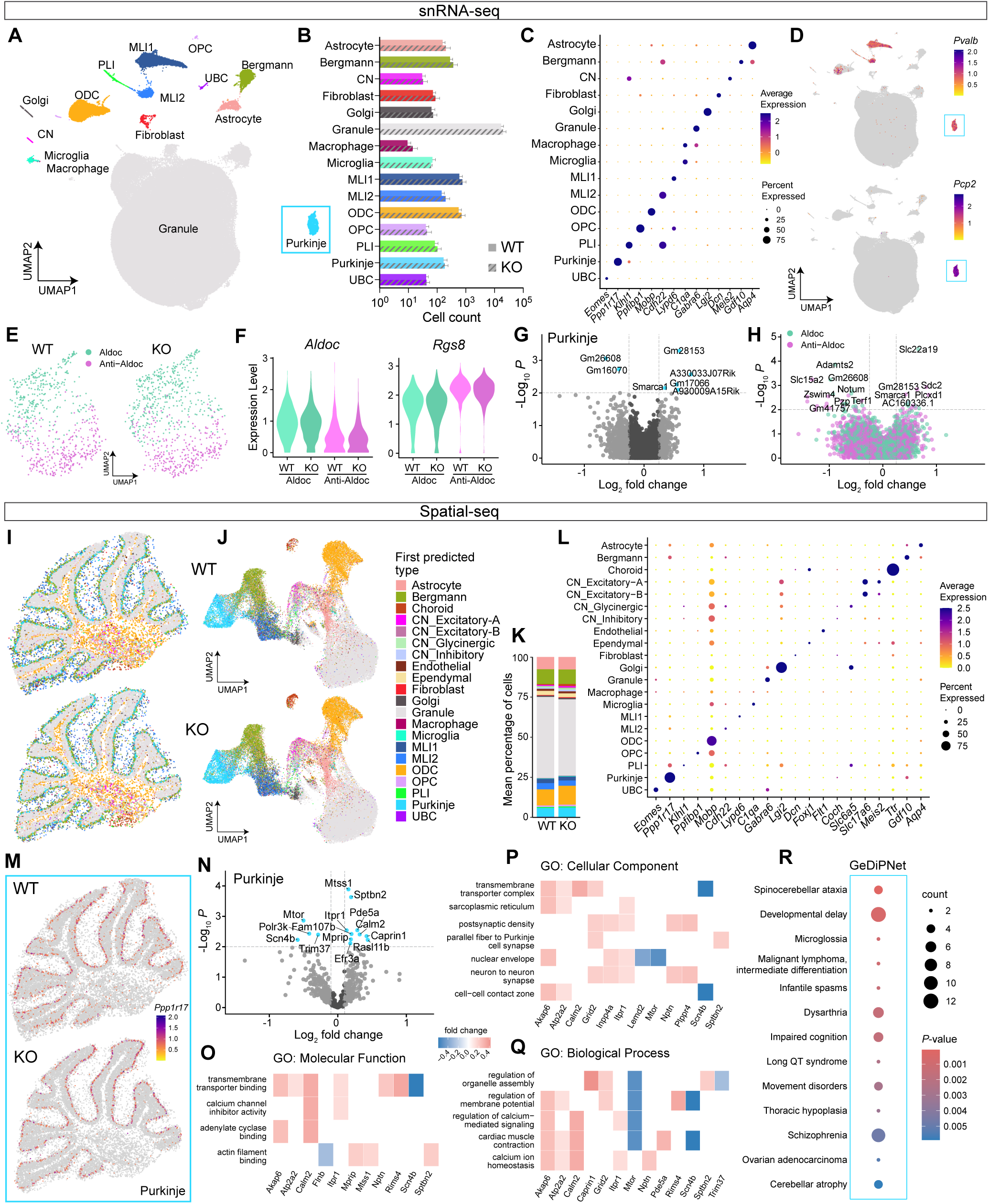
miR-206 regulates transcriptional programs related to synaptic function, neuronal excitability, and neurodevelopmental disorders. **(A)** UMAP plot of cerebellar cell types from WT and miR-206 KO mice. After performing unsupervised clustering, cells were annotated using a reference cerebellar snRNA-seq dataset^66^. CN, cerebellar nuclei neuron; MLI, molecular layer interneuron; ODC, oligodendrocyte; OPC, oligodendrocyte precursor; PLI, Purkinje layer interneuron; UBC, unipolar brush cell. **(B)** Total counts of each cell type are unaffected by miR-206 deletion. n = 4 WT, 4 KO male mice. **(C)** Dot plot of scaled expression of known marker genes for cerebellar cell types. **(D)** Feature plots detect expression of *Pvalb* mRNA in interneuron and Purkinje clusters and *Pcp2* mRNA expression only in the Purkinje cluster. **(E)** UMAP plot of Aldoc and Anti-Aldoc PC subtypes in WT and KO cerebellum, annotated using the same reference as in (A). **(F)** Violin plots show no difference in scaled expression of markers of Aldoc (*Aldoc*) and Anti-Aldoc (*Rgs8*) PC subtypes in WT and KO. **(G)** Volcano plot of differential gene expression in KO PCs. **(H)** Overlaid volcano plot of differential gene expression in KO Aldoc and Anti-Aldoc cells as compared to WT. **(I)** Representative spatial plots of first predicted cerebellar cell types (highest confidence RCTD assignments) from slide-seq (Curio Seeker) of WT (top) and miR-206 KO (bottom) sagittal cerebellar sections. Cell types were predicted by RCTD using the same reference as in (A) and a CN snRNA-seq reference^69^. **(J)** UMAP plots of cells in (I). **(K)** Average percentage of each cell type for WT and KO samples. n = 4 WT, 4 KO male mice. **(L)** Dot plot of scaled expression of known marker genes for cerebellar cell types. **(M)** Spatial feature plot of scaled expression of *Pp1r17*, a PC marker, in WT and KO samples. **(N)** Volcano plot of differential gene expression in KO PCs using C-SIDE analysis. **(O–Q)** Heat plots showing fold change of differentially expressed genes in PCs mapped to enriched molecular function (O), cellular component (P), and biological process (Q) gene ontology pathways. **(R)** Dot plot of Gene-Disease Pathway Networks (GeDiPNet) terms enriched in PC differentially expressed genes. Error bars indicate SEM.

Differentially expressed genes (DEGs) between KO and WT mice included noncoding RNAs and chromatin remodeling factor genes (Figures 3G, 3H, and S3G), suggesting that miR-206 influences post-transcriptional/translational regulatory pathways. However, the impact of miR-206 deletion on gene expression was subtle across all cerebellar cell types (Figure S3H, Tables S1 and S2), consistent with minimal perturbations in PC development and maturation in miR-206-deficient animals.

MicroRNAs canonically regulate cytoplasmic mRNA translation and stability, while snRNA-seq profiles nuclear gene transcripts^67,68^. Therefore, effects of miR-206 deletion on gene transcript levels may not be fully resolved by nuclear transcript profiling alone. To more comprehensively characterize the impact of miR-206 deletion on cerebellar gene networks, we performed Slide-seq whole-transcriptome profiling on cerebellar sections from WT and miR-206 KO mice. Slide-seq preserves native tissue architecture while enabling region-specific spatial transcriptomic profiling. Cell type annotation was performed using robust cell type decomposition (RCTD) with cerebellar cortex and cerebellar nuclei snRNA-seq reference datasets^66,69,70^. First-predicted cell types aligned with well-established cerebellar topography (Figure 3I). UMAP plots of unsupervised clustering (Figure 3J) as well as the proportions of each cell type (Figure 3K) were similar between WT and KO mice. Total marker gene expression for each predicted cell type was consistent with these annotations (Figure 3L). Spatial expression patterns of marker genes for PCs (Figure 3M) and other cell types (Figures S4A and S4F) were also similar in WT and KO mice.

DEGs in KO versus WT mice across cell types were predicted using cell type-specific inference of differential expression (C-SIDE)^71^ (Table S3 and Figures 3N and S4B–E). Top DEGs in PCs included *Mtss1*, *Sptbn2*, *Mtor*, and *Scn4b* (Figure 3N), which all have established roles in PC development and physiology^72–78^. PC DEGs were enriched in gene ontology pathways linked with ion channel and synapse function (Figures 3O–Q), and in gene networks associated with schizophrenia, ion channel-related disorders, ataxia, and developmental delay (Figure 3R).

As PCs provide the principal output of the cerebellar cortex to cerebellar nuclei neurons (CNs)^79,80^ and exert tonic inhibitory control over CN activity^81^, we focused our attention on these cells. The majority of DEGs in CNs were downregulated in KO animals (Figure S4G and Table S4)^82–88^. DEGs in CNs were enriched for genes involved in synaptic transmission, dendritic transport, mRNA processing, and phospholipid signaling (Figures S4H and S4I). DEGs were also associated with lactic acidosis and mitochondrial diseases (Figure S4J), consistent with metabolic stress. To confirm this observation, we performed HPLC-based metabolite profiling in whole cerebella from WT and KO mice. Increased lactate and succinate levels were detected in the KO mice (Figure S4K), while neurotransmitter and proteinogenic amino acid levels were unchanged (Figures S4L and S4M). Finally, when comparing DEGs across cerebellar cell types, we observed an overlapping pattern of enrichment in intellectual disability and neurodevelopmental delay or disorder pathways (Figure S4N).

These findings confirm that miR-206 is dispensable for the development, maturation, and topographical organization of PCs and other neuronal and non-neuronal cell types in the cerebellum. However, miR-206 deficiency influenced gene networks associated with mitochondrial stress in CNs and elevated extracellular lactate levels, consistent with altered neural activity or metabolic stressors in the cerebellum of miR-206 KO mice.

### miR-206 regulates sensorimotor gating in a sex-dependent manner

Individuals with schizophrenia and related neuropsychiatric disorders exhibit impaired sensorimotor gating, including PPI deficits^51,52,89^. PPI occurs when the magnitude of a startle reflex to a loud sound (pulse) is attenuated by a weaker, non-startling sound (prepulse) presented shortly before it^90,91^ (Figure 4A). Given the genetic and transcriptional links between miR-206 and schizophrenia-related disorders, together with growing evidence implicating the cerebellum in sensorimotor gating, we investigated whether miR-206 regulates PPI of the acoustic startle response.

**Figure 4.**
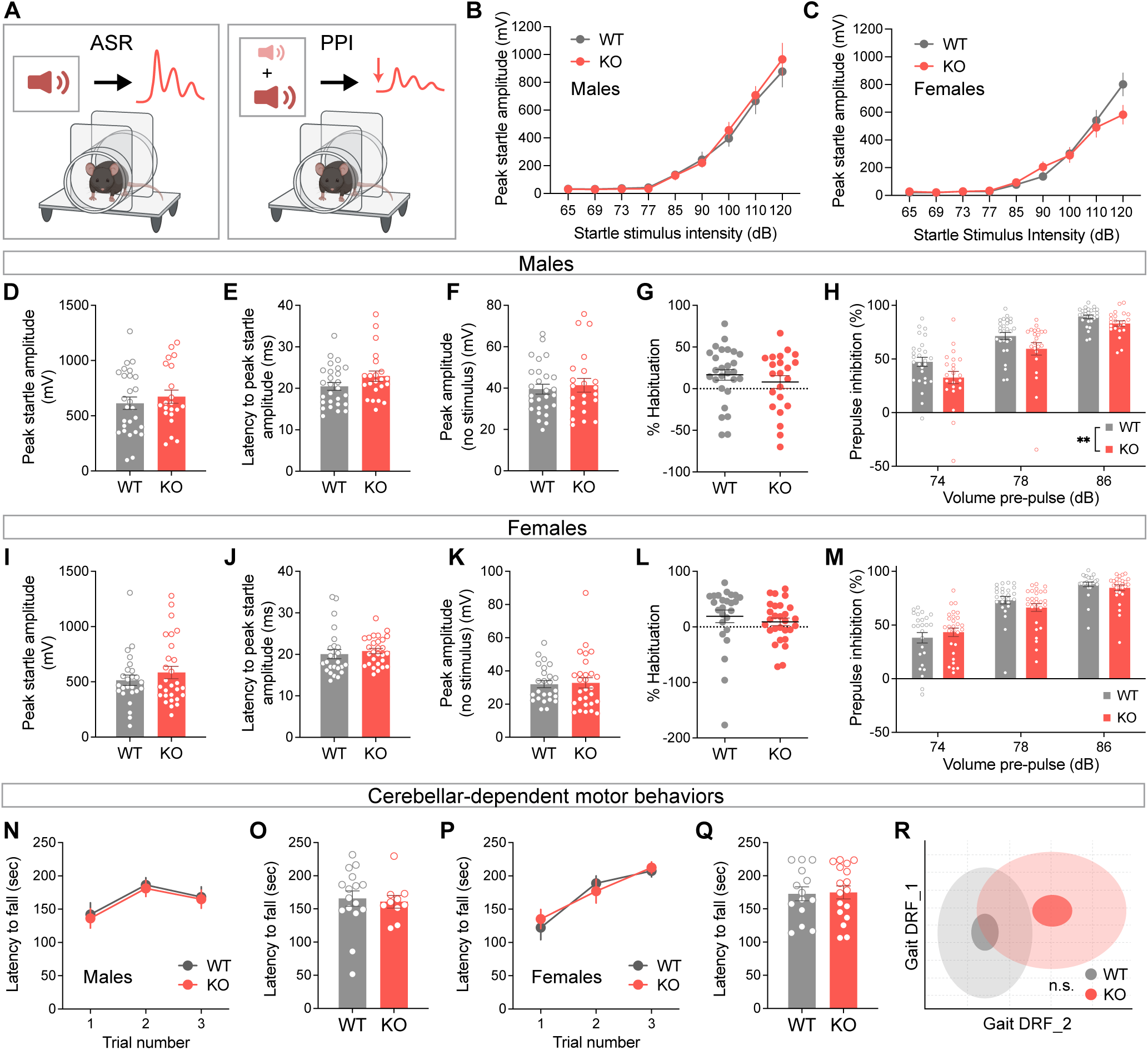
miR-206 deficiency induces sex-dependent impairment in sensorimotor gating. **(A)** Schematic of acoustic startle response of a mouse to a loud “pulse” of white noise (left) and prepulse inhibition (PPI), where a quieter acoustic prepulse preceding the pulse results in reduced amplitude of startle (right). **(B–C)** Acoustic startle threshold is unchanged in KO mice. n = 12 WT, 13 KO males (B), n = 12 WT, 11 KO females (C). **(D–G)** miR-206 deletion in male mice does not alter peak amplitude of the startle response to a 120 dB acoustic white noise stimulus (D), latency to peak startle (E), amplitude of baseline movement without stimulus (F), or habituation to successive startle stimuli during the PPI protocol (G). **(H)** miR-206 KO male mice display reduced PPI in response to 74, 78, and 86 dB prepulse stimuli. n = 27 WT, 22 KO mice. **p = 0.0012, main effect of genotype, two-way RM ANOVA. **(I–M)** miR-206 deletion in female mice does not alter peak amplitude of startle response (I), latency to peak startle (J), amplitude of baseline movement without stimulus (K), habituation to startle stimuli (L), or PPI (M). n = 12 WT, 11 KO female mice. **(N–Q)** Latency to fall from an accelerating rota-rod across three trials is not altered after miR-206 deletion. n = 16 WT, 10 KO males (N–O), 14 WT, 17 KO females (P–Q). **(R)** 2D representation of composite gait features that best discriminate between groups (DRF1 and DRF2, Brunner *et al.*, 2015). Center, small and large ellipses are the mean, standard error and standard deviation of the composite gait features for each group. WT and KO ellipses overlap, indicating similar gait characteristics (57% discrimination, p = 0.453). n = 14 WT, 15 KO males. Error bars indicate SEM.

The acoustic startle threshold was unaltered in male and female miR-206 KO mice relative to WT controls (Figures 4B and 4C), indicating intact auditory processing and brainstem startle reflex response. Male and female KO mice also exhibited no change in magnitude or latency of the acoustic startle response, baseline movement, or startle habituation (Figures 4D–G, 4I–L). However, PPI in male but not female KO mice was impaired relative to WT mice (Figures 4H and 4M).

As the cerebellum plays essential roles in motor learning and coordination^92,93^, we investigated whether impaired PPI in miR-206-deficient mice may reflect broader motor abnormalities. Male and female miR-206 KO mice exhibited normal motor learning on the accelerating rotarod and intact ambulatory gait relative to WT controls (Figures 4N–R), indicating preserved cerebellar-dependent motor coordination. However, male but not female KO mice displayed subtle hypolocomotion in stressful contexts, including exposure to a novel open field, isolation in a clean home cage, and acute restraint stress^94–96^, without alterations in vertical activity or body mass (Figure S5). Stress-induced hyperthermia and anxiety-like behaviors in the dark-light emergence, tail suspension, and hyponeophagia tests were unchanged in KO mice of both sexes (Figure S6), indicating that loss of miR-206 does not produce heightened anxiety-like behavior.

Fear states induced by acute stressors can reduce locomotor output concurrent with the onset of freezing behavior^97^, and the cerebellum regulates fear responses in mice^98,99^. During the training phase of a classical fear conditioning procedure, no differences in footshock-induced freezing or locomotion were observed in male or female KO mice, suggesting no change in innate fear responses (Figure S7). However, male KO mice displayed modest increases in freezing during contextual fear memory retrieval and cued fear extinction (Figure S7), consistent with enhanced fear memory encoding in the male KO mice. The cerebellum is implicated in social behaviors^32,37^, and its dysfunction is linked to social deficits in neurodevelopmental disorders including ASD^39,40,100,101^. Disruption of miRNA function in PCs also leads to social deficits in mice^28^. Social approach behavior of male and female miR-206 KO mice was similar to WT animals, with all groups showing a strong preference for a novel mouse over a novel object (Figure S8), suggesting that miR-206 does not play a major role in cerebellar-mediated social behavior.

These findings suggest that sensorimotor gating is selectively disrupted in miR-206-deficient male mice without accompanying alterations in other behaviors known to be regulated by the cerebellum, including motor coordination, stress- and anxiety-related behaviors, and social preference.

### miR-206 acts in cerebellar PCs to regulate sensorimotor gating

We sought to confirm that miR-206 acts in PCs to regulate sensorimotor gating. As PCs express parvalbumin (PV), we bred *miR-206^fl/fl^* mice to mice expressing Cre in PV^+^ neurons (*Pvalb^Cre^* mice) including PCs (Figure 5A). Loss of miR-206 in the cerebellum of *miR-206^fl/fl^*::*Pvalb^Cre^* conditional knockout mice was confirmed by qPCR (Figure 5B). Similar to the whole-body miR-206 KO mice, male but not female mice with PV^+^ neuron-specific miR-206 deletion displayed deficits in PPI but not startle amplitude or latency (Figures 5C–E, males; Figures S9A–C, females).

**Figure 5.**
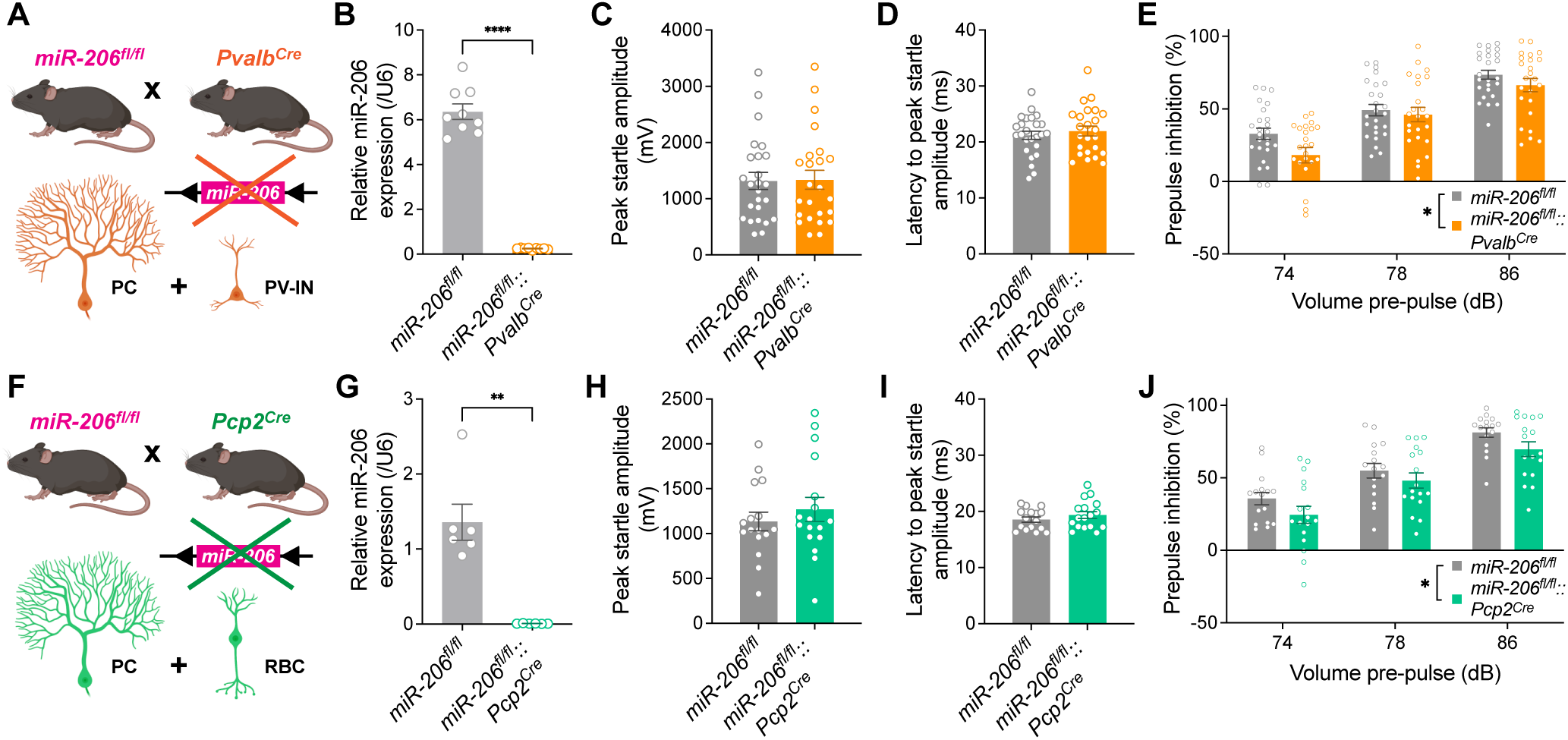
miR-206 acts exclusively in Purkinje cells to regulate sensorimotor gating. **(A)** Schematic of miR-206 conditional deletion in parvalbumin (PV)-expressing neurons including PCs and interneurons (PV-IN). **(B)** qRT-PCR shows that miR-206 is effectively deleted in cerebellum of *miR-206^fl/fl^*::*Pvalb^Cre^* mice. n = 9 *miR-206^fl/fl^*, 9 *miR-206^fl/fl^*::*Pvalb^Cre^* male mice. ***p<0.0001, Welch’s t-test. **(C–E)** Peak startle amplitude (C) and latency to peak startle (D) are unaltered but PPI (E) is impaired in mice with conditional deletion of miR-206 in parvalbumin-expressing neurons. n = 25 *miR-206^fl/fl^*, 24 *miR-206^fl/fl^*::*Pvalb^Cre^* male mice. *p = 0.0202, main effect of genotype, two-way RM ANOVA. **(F)** Schematic of miR-206 conditional deletion in PCP2-expressing neurons comprising PCs and retinal bipolar neurons (RBC). **(G)** qRT-PCR shows that miR-206 is effectively deleted in cerebellum of *miR-206^fl/fl^*::*Pcp2^Cre^*mice. n = 6 *miR-206^fl/fl^*, 6 *miR-206^fl/fl^*::*Pcp2^Cre^*male mice. **p = 0.0024, Welch’s t-test. **(H–J)** Peak startle amplitude (H) and latency to peak startle (I) are unchanged but PPI (J) is impaired in mice with conditional deletion of miR-206 in PCs and retinal bipolar neurons. n = 16 *miR-206^fl/fl^*, 17 *miR-206^fl/fl^*::*Pcp2^Cre^*male mice. *p = 0.0153, main effect of genotype, two-way RM ANOVA. Error bars indicate SEM.

Schizophrenia patients exhibit lower numbers of cortical PV^+^ interneurons and/or reduced cortical PV expression levels relative to unaffected controls^102,103^. It was previously reported that miR-206 is expressed by inhibitory PV^+^ interneurons in the neocortex^29^. While we found that miR-206 levels were undetectable in neocortex during development and in adulthood (Figures 1 and S1), it is possible that genetic ablation of even low levels of miR-206 reduces numbers of PV^+^ interneurons or *Pvalb* expression in the cortex, which contributes to PPI deficits in the KO mice. However, we found that numbers of PV immunoreactive neurons were similar in the cortex and across all other sampled brain regions of WT and miR-206 KO mice (Figures S9D and S9E). *Pvalb* transcript levels were also comparable throughout the brains of WT and KO mice (Figure S9F).

To generate converging genetic evidence that miR-206 acts in cerebellar PCs to regulate sensorimotor gating, we bred *miR-206^fl/fl^*mice to *Pcp2^Cre^* mice, in which Cre expression in the brain is restricted to cerebellar PCs (Figure 5F). qPCR confirmed that cerebellar miR-206 expression was ablated in *miR-206^fl/fl^*::*Pcp2^Cre^* mice (Figure 5G). Male but not female mice with PC-specific miR-206 deletion displayed deficits in PPI relative to controls, with no change in startle amplitude or latency (Figures 5H–J, males; Figures S9G–I, females). Collectively, these findings demonstrate that miR-206 signaling in cerebellar PCs regulates sensorimotor gating in male animals.

### HITS-CLIP identifies miR-206-regulated transcripts in Purkinje cells

Canonically, miRNAs bind complementary sequences within the 3′UTRs of target transcripts to facilitate their recruitment to the RISC complex, resulting in translational repression^4,5^. Although some RISC-associated transcripts subsequently undergo destabilization and degradation, they generally exhibit minimal changes in steady-state abundance following miRNA binding^4,5^, making conventional RNA sequencing insufficient for comprehensively identifying miRNA-regulated transcripts. Therefore, to identify transcripts undergoing miR-206-mediated translational repression, we performed high-throughput sequencing after crosslinking immunoprecipitation (HITS-CLIP) of the RISC-associated protein Argonaute-2 (AGO2) in cerebellar PCs from miR-206-deficient mice^104,105^. *Pvalb^Cre^* mice were bred to mice expressing Cre-dependent FLAG-tagged AGO2, thereby producing tagged AGO2 in PCs (Figure 6A). These mice were then bred to either miR-206 floxed mice to produce conditional deletion of miR-206 in PV^+^ neurons (“*Pvalb^Cre^*-cKO”) or to WT mice as controls (“*Pvalb^Cre^*-WT”). To generate a second, independent dataset of high-confidence miR-206 target transcripts in PCs, we crossed *Pcp2^Cre^* mice with the Cre-dependent FLAG-tagged AGO2 mice and then bred the resulting mice to miR-206 floxed or WT mice. The HITS-CLIP procedure was performed using whole cerebellar tissue from cKO and WT mice and AGO2-bound RNA was sequenced (Figure 6A). Loss of AGO2-associated miR-206 sequencing reads in both *Pvalb^Cre^* and *Pcp2^Cre^*cKO but not WT mice confirmed effective deletion of miR-206 (Figure 6B).

**Figure 6.**
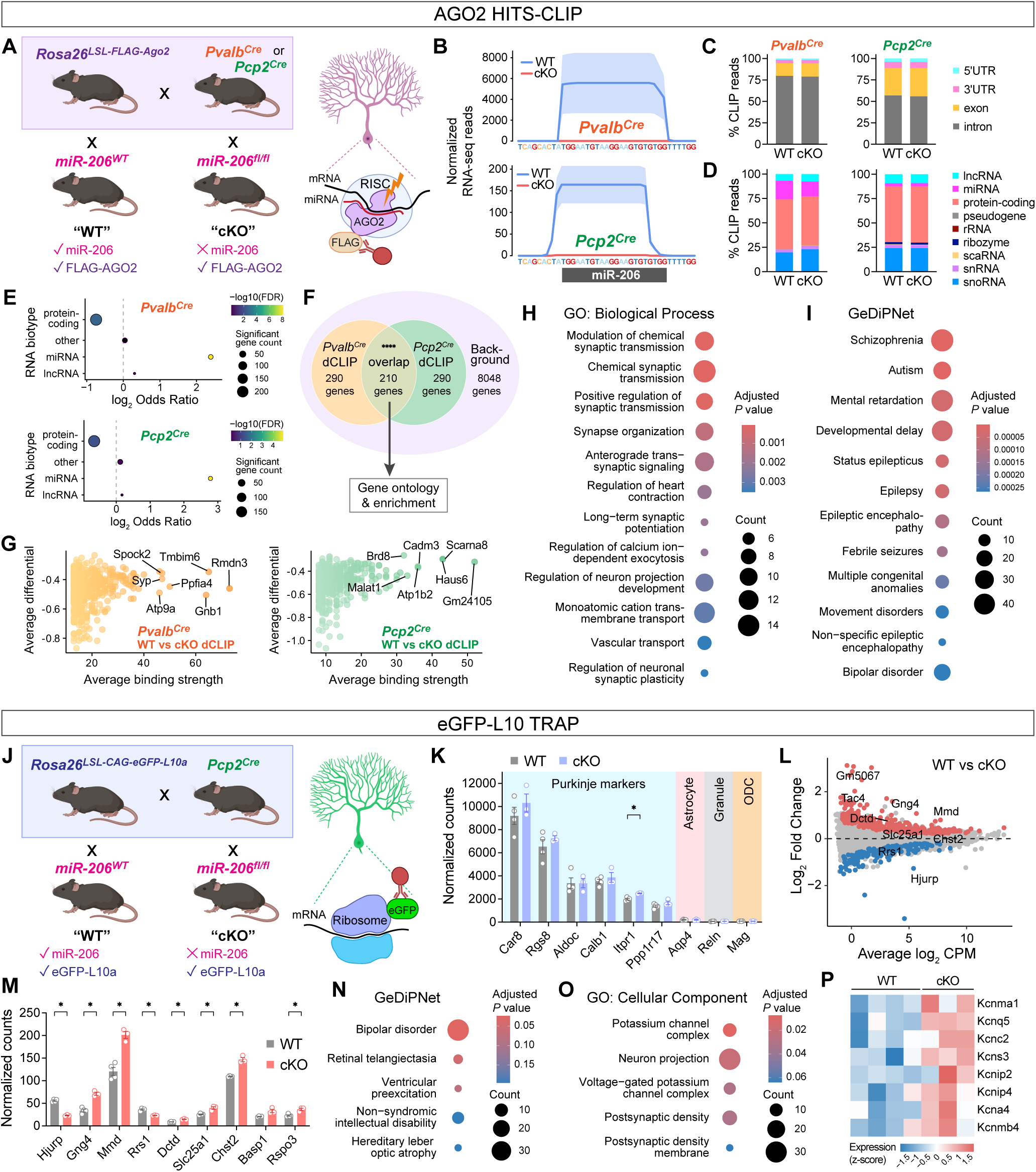
Profiling of RISC- and ribosome-associated transcripts identifies miR-206 targets relevant to neuronal excitability. **(A)** Schematic of breeding strategy for comparative high throughput sequencing after crosslinking immunoprecipitation (HITS-CLIP) of RISC-associated RNAs. Cre expression downstream of the *Pvalb* or *Pcp2* promoter in PCs drives production of FLAG-tagged Ago2 from the *Rosa26* locus, as well as deletion of miR-206 in the presence of floxed but not WT miR-206 alleles. microRNAs and mRNA targets are bound by FLAG-tagged Ago2 within the RISC of PCs and isolated using HITS-CLIP, and differential mRNA binding in the presence (“WT”) or absence (“cKO”) of miR-206 can be determined. **(B)** Mean normalized HITS-CLIP RNA-seq reads mapped to the miR-206 locus are abundant in WT but not detected in cKO mice. n = 5 WT, 5 cKO male mice for each Cre line. **(C)** Percentage of CLIP reads mapped to gene transcript regions for *Pcp2^Cre^* and *Pvalb^Cre^* HITS-CLIP experiments. **(D)** Percentage of CLIP reads mapped to different RNA biotypes. **(E)** Bubble plot of differentially expressed gene number, significance, and odds ratio (WT vs cKO) for different RNA biotypes shows highest significant gene count for protein coding genes but highest odds ratio and significance for miRNAs. **(F)** Transcripts with differential Ago2 binding following miR-206 deletion were detected using comparative dCLIP analysis^111^. 210 differentially bound transcripts were common to both *Pvalb^Cre^* and *Pcp2^Cre^* data sets. Input was 8838 genes with non-zero counts. ****p = 3.81x10^-142^, Odds Ratio = 20.1, Fisher’s exact test. **(G)** Scatter plots of average binding strength and differential for top dCLIP-predicted transcripts from *Pvalb^Cre^* and *Pcp2^Cre^* datasets. **(H–I)** The 210 overlapping transcripts were used for gene ontology enrichment analyses. Biological Process (H) Gene Ontology (GO) terms related to synaptic transmission and ion channel activity were particularly enriched, while disease-associated enrichment using GeDiPNet (Gene Disease Pathway Networks) (I) revealed significant overlap with genes linked to schizophrenia, autism, and developmental delay. **(J)** Schematic of breeding strategy for comparative translating ribosome affinity purification (TRAP). Cre expression downstream of the *Pcp2* promoter in PCs drives production of eGFP-tagged L10 ribosomal subunit from the *Rosa26* locus, as well as deletion of miR-206 in the presence of floxed but not WT miR-206 alleles. mRNAs within translating ribosomes are pulled down and sequenced, and differential expression assessed between miR-206 WT and cKO mice. **(K)** Markers for PCs but not astrocytes, granule cells, or oligodendrocytes (ODC) are enriched after TRAP-seq. Expression of *Itpr1* is increased but other markers are unchanged in miR-206 cKO mice. n = 4 WT, 3 cKO male mice. *p = 0.0277, multiple t tests + Bonferroni correction. **(L)** MA plot of differential expression of TRAP-seq mRNAs. Red and blue colors denote significantly increased or decreased expression in cKO, respectively. **(M)** Normalized counts of top differentially expressed TRAP-seq mRNAs by *P*-value. *p < 0.05, multiple t tests + Bonferroni correction. **(N)** GeDiPNet terms including bipolar disorder are enriched in TRAP-seq DE genes. **(O)** Cellular component gene ontology analysis reveals enrichment of potassium channel complex-related pathways among TRAP-seq DE genes. **(P)** Heat map of scaled TRAP-seq expression of top differentially expressed potassium channel transcripts in cKO. Error bars indicate SEM.

For each comparative HITS-CLIP experiment (*Pvalb^Cre^*-cKO and *Pcp2^Cre^*-cKO *vs.* respective WT mice), sequencing reads mapped predominantly to intronic regions, with reads concentrated in 3’UTRs and 5’UTRs, and exon reads detected to a lesser extent (Figure 6C). As might be expected, protein-coding mRNAs accounted for the largest relative proportion of RISC-associated RNA transcripts that differed between both cKO lines and WT mice (Figures 6D and 6E, Tables S5 and S6). Notably, neither RISC association nor transcript abundance of previously reported miR-206 target genes from other cell types and tissues differed between both lines of cKO mice and WT controls (Figures S10A and S10B)^25,26,58,106–110^.

In addition to protein-coding mRNAs, a variety of other RNA biotypes were also pulled down with AGO2, including miRNAs, lncRNAs, and snoRNAs (Figure 6D). Analyses of all RNA biotypes revealed striking alterations in AGO2-associated miRNAs between the cKO mice and controls (Figure 6E, Tables S5 and S6), with miRNAs representing the most significantly altered RNA biotype in miR-206-deficient PCs. This suggests that miR-206 deletion remodels the composition of AGO2-associated miRNA networks and that miR-206 broadly shapes miRNA-mediated post-transcriptional regulatory programs in cerebellar PCs.

To identify high-confidence gene transcripts directly targeted by miR-206, we next performed differential CLIP (dCLIP) analysis^111^. Whereas HITS-CLIP defines the overall landscape of AGO2-associated transcripts, dCLIP identifies transcripts that lose AGO2 binding following miR-206 deletion, enriching for transcripts directly targeted by miR-206 and the broader remodeling of AGO2-associated miRNA networks in miR-206-deficient animals. We performed dCLIP analyses independently in the *Pvalb^Cre^*-cKO and *Pcp2^Cre^*-cKO datasets by comparing AGO2 occupancy in each cKO line relative to its corresponding WT control. Transcripts exhibiting reduced AGO2 association after miR-206 deletion significantly overlapped between the two independent conditional knockout models (Figures 6F and 6G, Tables S7–S9), increasing the likelihood that they represent miR-206-regulated targets in PCs. Many overlapping transcripts were also independently predicted to contain miR-206 binding sites by the miRNA target prediction algorithm TargetScan^112^ (bold font in tables; full list in Table S10), supporting their identification as high-confidence direct miR-206 targets.

Gene ontology analyses of dCLIP-predicted miR-206 targets that overlapped between both cKO lines revealed enrichment for pathways related to ion channel function and synaptic transmission (Figure 6H), together with disease-associated gene networks linked to schizophrenia, developmental delay, ASD, and epilepsy (Figure 6I).

### TRAP-seq profiling of the miR-206 translatome in Purkinje cells

miRNA-mediated recruitment of transcripts to the RISC complex suppresses protein synthesis through translational repression and transcript destabilization^4,5^. However, some miRNA target transcripts, particularly highly abundant mRNAs, can potentially remain associated with translating ribosomes despite miRNA-mediated repression, permitting residual protein synthesis^4^. To identify mRNAs with altered ribosome association following miR-206 deletion and define the miR-206-regulated translatome, we performed translating ribosome affinity purification and sequencing (TRAP-seq)^113,114^ in PCs from WT and miR-206-deficient mice. To isolate ribosome-associated RNAs selectively from PCs, *Pcp2-Cre* mice were bred to mice expressing Cre-dependent eGFP-tagged ribosomal protein L10a (*Rosa26^LSL-CAG-eGFP-L10a^*mice), thereby producing tagged ribosomes in PCs (Figure 6J). These mice were then crossed with miR-206 WT mice as controls (“WT”) or with miR-206 floxed mice to generate PC-specific miR-206 conditional knockout mice (“cKO”). TRAP-seq was performed using whole cerebellar tissue from WT and cKO mice, and eGFP-L10a-associated ribosome-bound RNAs were affinity purified and sequenced.

Sequencing reads were highly enriched for PC marker genes in WT and cKO mice, with minimal representation of markers from other cerebellar cell types (Figure 6K). Expression analyses identified multiple ribosome-associated transcripts that differed between WT and cKOs (Figures 6L and 6M, Table S11), although the functions of many of these transcripts in PCs are currently unknown. Similar to the HITS-CLIP findings, we observed no changes in previously reported miR-206 targets identified in other tissues (Figure S10C)^109,110^, or in transcripts identified following antisense sponge-mediated miR-206 knockdown in PCs (Figure S10D)^28^. qPCR also found no effect of miR-206 deletion on expression of previously reported miR-206 target genes, including *Esr1*^115^ and *Bdnf*^25,26,58,106–108^, across multiple brain regions (Figures S10E and S10F).

However, altered ribosome-associated transcripts in cKO mice were enriched for gene networks linked to neuropsychiatric disorders (Figure 6N), as well as pathways related to neuronal excitability (Figure 6O). In particular, multiple voltage-gated potassium channel transcripts exhibited increased ribosome association in cKO mice (Figure 6P), suggesting that miR-206 regulates translational programs controlling the intrinsic excitability of PCs.

### miR-206 deficiency increases the spontaneous firing of Purkinje cells

PCs exhibit spontaneous action potential firing driven by intrinsic voltage- and calcium-dependent conductances^116^, and abnormalities in PC firing dynamics have been linked to behavioral and movement disorders^117^. Both HITS-CLIP and TRAP-seq analyses showed that miR-206 regulates gene programs that gate neuronal excitability. We therefore investigated whether miR-206 regulates the intrinsic firing properties of PCs in lobule VI of the posterior cerebellar vermis, a cerebellar region implicated in schizophrenia and other neuropsychiatric disorders^43,118–120^ (Figure 7A).

**Figure 7.**
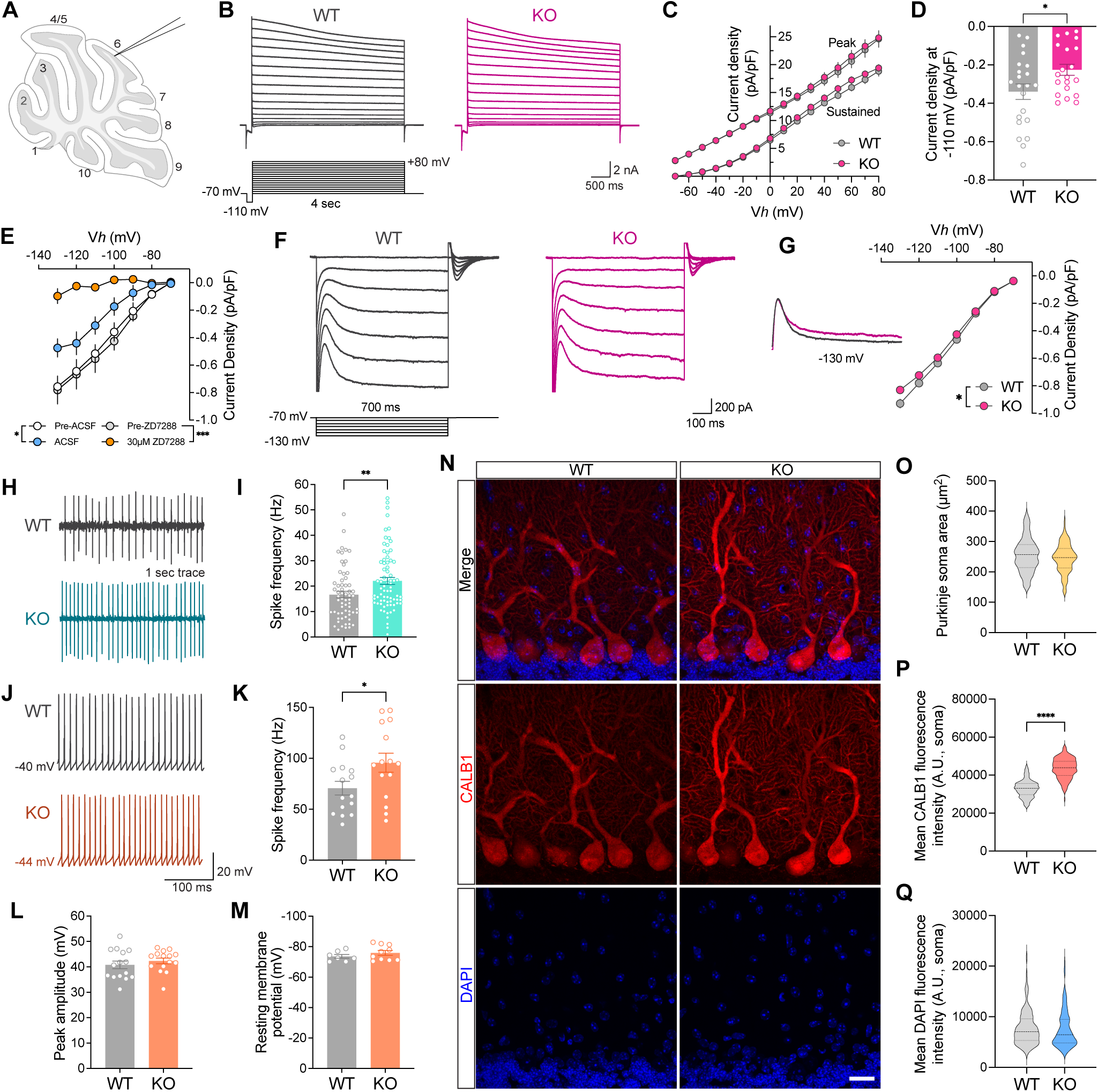
miR-206 deficiency induces Purkinje cell hyperactivity. **(A)** *Ex vivo* electrophysiology recording pipette positioned in lobule VI of cerebellar vermis. **(B)** Example whole-cell current traces recorded from WT and KO PCs during the voltage step function used to elicit potassium current in PCs. Cells were held at -70 mV, hyperpolarized at -110 mV, and depolarized from -70 to +80 mV in 10 mV increments. **(C)** Peak and sustained current-voltage curve for depolarizing potentials in (B) does not reveal differences in KO. Sustained current was measured four seconds after each depolarizing voltage step. **(D)** Reduced hyperpolarization-activated current at -110 mV in KO PCs. n = 22 WT, 20 KO cells from 8 WT, 8 KO male mice (C–D). *p = 0.0287, Welch’s t-test. **(E)** Hyperpolarization-activated current in WT PCs is blocked by the HCN channel blocker ZD7288; note that current density at hyperpolarizing holding potentials before and after application of ACSF or ZD7288 shows significant run-down with repeated hyperpolarization. n = 4 WT neurons per group. ***p = 0.0007, main effect pre vs post ZD7288; *p = 0.0390, main effect pre vs post ACSF; two-way RM ANOVA. **(F)** Example whole-cell hyperpolarization-activated current traces in WT and KO PCs. Cells were held at -70 mV and hyperpolarized in 10 mV increments down to -130 mV. **(G)** Hyperpolarization-activated current in miR-206 KO PCs is reduced with greater hyperpolarization. n = 74 WT, 72 KO cells from 10 WT, 10 KO male mice. *p = 0.0387 (voltage x genotype interaction), two-way RM ANOVA. **(H)** Representative cell-attached patch clamp recording traces of spontaneous firing in WT and KO PCs. **(I)** Action potential frequency is increased in KO PCs (cell-attached). n = 66 WT, 79 KO neurons from 8 WT, 10 KO male mice. **p = 0.0051, Welch’s t-test. **(J)** Representative whole-cell recording traces of spontaneous firing in WT and KO PCs. **(K)** Action potential frequency is increased in KO PCs (whole-cell). n = 15 WT, 15 KO neurons from 4 WT, 3 KO male mice. *p = 0.0395, Welch’s t-test. **(L)** Peak action potential amplitude and **(M)** resting membrane potential are not altered in KO PCs (whole-cell). **(N)** Representative orthogonal projections of confocal z-stack images of WT and KO PCs in lobule VI of cerebellar vermis stained for calbindin 1 (CALB1, red) and DAPI (blue). Scale bar: 20 µm. **(O)** Purkinje soma cross section area is similar in WT and KO. **(P)** CALB1 immunofluorescence intensity is elevated in KO PC soma. n = 162 WT, 148 KO neurons from 4 WT, 4 KO mice. ****p < 0.0001 and **p = 0.0013, Welch’s t-test (neuron-level and mouse-level analysis, respectively). **(Q)** DAPI fluorescence intensity is equivalent in WT and KO PC soma. Error bars indicate SEM.

We first assessed effects of miR-206 deletion on voltage-dependent membrane currents using whole-cell voltage clamp recordings while pharmacologically inhibiting sodium and calcium currents (Figure 7B). Peak and sustained outward currents evoked by depolarizing voltage steps were unchanged in KO PCs (Figure 7C). In contrast, hyperpolarization-evoked inward currents were reduced following miR-206 deletion (Figure 7D). These currents were effectively blocked by the hyperpolarization-activated cyclic nucleotide-gated (HCN) channel inhibitor ZD7288 (Figure 7E), indicating that they were mediated predominantly by HCN channels. HCN channels are activated by hyperpolarization and cyclic AMP signaling, are permeable to sodium and potassium, and are key regulators of intrinsic excitability^121,122^. Consistent with reduced HCN channel activity, currents evoked by hyperpolarizing voltage steps were attenuated in KO PCs (Figures 7F and 7G), although the magnitude of this effect may have been underestimated due to their rapid rundown during whole-cell recordings^123,124^.

We next examined whether altered membrane conductances in miR-206 deficient animals influenced the firing dynamics of PCs. Cell-attached recordings revealed increased spontaneous firing frequency in PCs from KO mice relative to WT controls (Figures 7H and 7I). Increased spontaneous firing was similarly observed during whole-cell current clamp recordings, with no changes in resting membrane potential or action potential amplitude (Figures 7J–7M). Notably, CALB1 immunofluorescence intensity was higher in PCs of KO mice (Figures 7N–Q), consistent with increased calcium handling associated with elevated spontaneous activity.

### miR-206 promotes Purkinje cell burst firing

We next investigated whether increasing miR-206 availability in mature PCs alters their intrinsic firing dynamics. The *pre-miR-206* stem-loop sequence was cloned and functional miR-206 expression was validated *in vitro* by northern blot analysis (Figure 8A) and 3′UTR luciferase reporter assay (Figure 8B). GFP alone, or *pre-miR-206* with GFP, were packaged into the PHP.eB AAV serotype under the control of the *Pcp2* promoter (AAV-control and AAV-miR-206, respectively). After intravenous (IV) administration (Figure 8C), GFP expression was detected exclusively in PCs throughout the cerebellum of both AAV-control and AAV-miR-206 treated mice (Figure 8D). qPCR confirmed elevated mature miR-206 expression in the cerebellum of AAV-miR-206 mice relative to AAV-control mice (Figure 8E).

**Figure 8.**
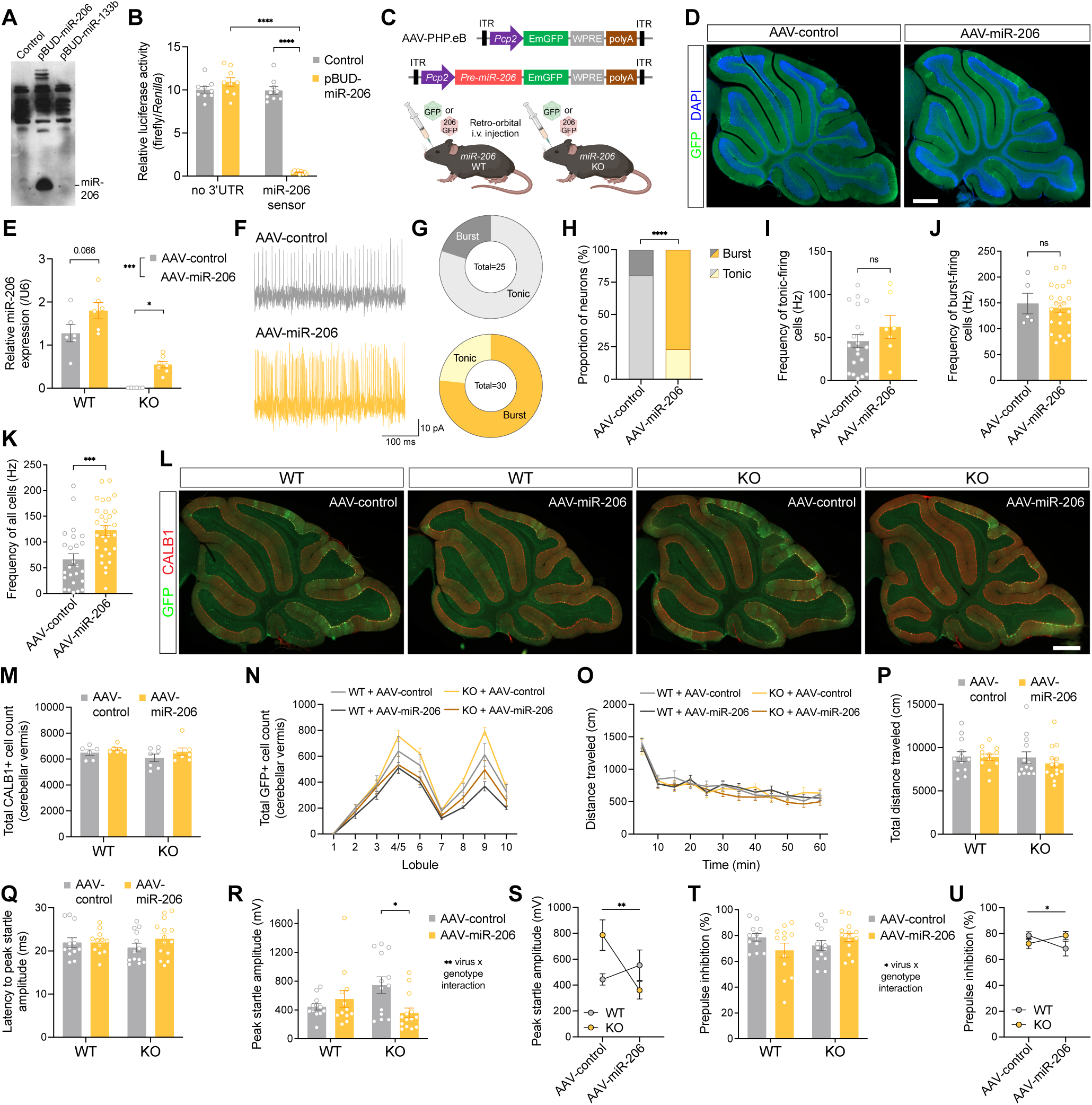
Viral-mediated miR-206 expression promotes Purkinje cell burst firing and bidirectionally regulates sensorimotor gating. **(A)** Northern blot detects mature miR-206 in COS cells transfected with miR-206-expressing plasmid (pBUD-miR-206-EGFP) but not with control (pBUD-EGFP) or miR-133b-expressing (pBUD-pre-miR133b-EGFP) plasmids. **(B)** miR-206 expression in COS cells knocks down a miR-206 luciferase sensor containing a 3’UTR comprised of miR-206 antisense sequences. Relative firefly to *Renilla* luciferase activity was measured 24 h after transfection. Data shown are the mean of three experiments. n = 9 per group. ****p < 0.0001, main effects of miRNA plasmid and luciferase plasmid, two-way ANOVA with Tukey’s multiple comparisons test. **(C)** PHP.eB AAV vectors for expression of miR-206 and GFP in PCs. Control virus (top) expresses GFP while miR-206 virus (bottom) expresses both GFP and *pre-miR-206* downstream of the PC-specific *Pcp2* promoter. Viruses were injected intravenously in WT and miR-206 KO mice by retro-orbital injection at one month of age. **(D)** GFP-expressing PCs (green) are visible in sagittal sections of cerebellar vermis 3 weeks after retro-orbital injection of KO mice with control or miR-206 viruses. DAPI, blue. Scale bar: 500 µm. **(E)** qRT-PCR detection of mature miR-206 in whole cerebellum shows a trend to increased miR-206 expression in WT and rescue of miR-206 to ∼30% of WT levels in KO six months after viral delivery. n = 6 male mice per group. ***p = 0.0009, main effect of virus, *p = 0.0469 (KO/control virus vs KO/miR-206^OE^ virus), two-way ANOVA and Tukey’s multiple comparisons test. **(F)** Representative extracellular recording traces of spontaneous action potentials of WT PCs one month after delivery of control or miR-206 virus. **(G)** Proportion of PCs exhibiting tonic or bursting modes of spontaneous action potential firing in control (top, grey) and miR-206 (bottom, gold) mice. **(H)** Stacked bar plot of percentage of PC firing types in (G). Most PCs infected with control virus fire tonically, while most PCs infected with miR-206 virus display burst firing. n = 25 AAV-control, 30 AAV-miR-206 infected neurons from 5 AAV-control, 5 AAV-miR-206 infected mice. ****p < 0.0001, Fisher’s exact test. **(I)** Mean frequency of tonically firing PCs is unaffected by viral miR-206 expression. n = 20 AAV-control, 7 AAV-miR-206 infected neurons. **(J)** Mean frequency of bursting PCs is unchanged with viral miR-206 expression. n = 5 AAV-control, 23 AAV-miR-206 infected neurons. **(K)** Frequency of all PCs is increased with viral miR-206 expression. ***p = 0.0004 (n = neurons), **p = 0.0064 (n = mice). **(L)** Representative images of calbindin 1(CALB1, red) immunofluorescence and GFP (green) signal in sagittal sections of cerebellar vermis of WT and KO mice infected with control or miR-206 virus for six months. Scale bar: 500 µm. **(M)** PC numbers (CALB1 immunoreactive cells) are similar in WT and KO six months after infection with control or miR-206-expressing viruses. Counts were summed across six 50-µm sections of cerebellar vermis. n = 6–7 mice per group. **(N)** Number of GFP positive PCs for each lobule of cerebellar vermis in WT or KO mice infected with control or miR-206-expressing virus. n = 6–7 mice per group. **(O)** Distance traveled in the open field by WT or KO mice infected with control or miR-206 expressing virus. n = 12–14 mice per group (Same n for O–U). **(P)** Summary open field data from (O). **(Q)** Latency to peak acoustic startle amplitude is similar for WT and KO mice infected with control or miR-206 expressing viruses. **(R–S)** Peak startle amplitude is differentially affected by viral miR-206 expression in WT and KO mice. **p = 0.0099, virus x genotype interaction; *p = 0.0191, KO/AAV-control vs KO/AAV-miR-206; two-way ANOVA and Tukey’s multiple comparisons test. **(T–U)** PPI is differentially affected by viral miR-206 expression in WT and KO mice exposed to 78 dB prepulse. *p = 0.0478, two-way ANOVA and Tukey’s multiple comparisons test. Error bars indicate SEM.

Extracellular recordings from PCs in lobule VI of the cerebellar vermis were performed in AAV-control and AAV-miR-206 mice. The majority of PCs (∼80%) exhibited tonic spontaneous firing in AAV-control mice, whereas only a minority (∼20%) displayed spontaneous high-frequency burst firing (Figures 8F–H). In contrast, the enhanced miR-206 expression in AAV-miR-206 mice induced a marked shift in PC firing state, with burst-firing PCs accounting for ∼80% of recorded neurons (Figure 8F–H). However, the firing frequencies of both tonic and burst-firing modes themselves remained unchanged in AAV-miR-206 mice compared to controls (∼50 Hz *vs.* ∼150 Hz, respectively) (Figures 8I and 8J), such that altered firing mode alone contributed to the overall increased firing frequency with AAV-miR-206 expression (Figure 8K). This suggests that miR-206 dosage bidirectionally tunes PC firing dynamics by biasing neurons between tonic and burst firing states.

### miR-206 acts in Purkinje cells to bidirectionally regulate sensorimotor gating

Finally, we investigated whether miR-206 dosage in mature cerebellar PCs bidirectionally regulates sensorimotor gating. AAV-control and AAV-miR-206 were administered by IV injection to both WT and miR-206 KO mice (Figure 8L). We confirmed that enhancing miR-206 expression caused no loss of CALB1 immunoreactive PCs in WT or miR-206 KO mice (Figure 8M), indicating that prolonged miR-206 augmentation did not impair PC survival. Furthermore, GFP expression was visible in PCs in both AAV-control and AAV-miR-206 treated mice at the time of behavioral testing, which occured 5 months after viral administration (Figure 8L). GFP signal intensity was lower in AAV-miR-206 mice (Figure 8N), likely reflecting the larger and more complex transcriptional payload of the AAV-miR-206 construct, which co-expressed GFP together with the *pre-miR-206* stem-loop sequence. However, the overall distribution of GFP-expressing PCs was similar across groups (Figure 8N), suggesting that miR-206 augmentation was broadly achieved throughout the cerebellum without evidence of loss of PC or altered viral tropism.

Locomotor activity in an open field apparatus was similar across WT and KO mice treated with AAV-miR-206 and AAV-control, suggesting that enhancing miR-206 expression in PCs of WT and KO mice did not disrupt cerebellar-mediated motor behaviors (Figures 8O and 8P). The latency to peak acoustic startle was similarly unchanged (Figure 8Q). However, PC-specific miR-206 expression exerted opposing effects on startle amplitude in WT and KO mice treated with AAV-miR-206, increasing peak startle amplitudes in WT animals while reducing amplitudes in miR-206 KO mice (Figures 8R and 8S). Strikingly, miR-206 dosage also bidirectionally regulated PPI, impairing PPI in WT mice while rescuing the constitutive PPI deficit in miR-206 KO animals (Figures 8T and 8U). This shows that miR-206 dosage in cerebellar PCs bidirectionally regulates sensorimotor gating behavior.

## DISCUSSION

Genetic and transcriptional evidence links miR-206 to schizophrenia and other neuropsychiatric disorders^18–24^. Deficits in sensorimotor gating, including impaired prepulse inhibition (PPI), are a well-established feature of schizophrenia and related disorders^52,54,125,126^. Using complementary RNAscope, qPCR, and transcriptomic approaches, we found that miR-206 expression in postnatal brain is restricted to cerebellar PCs, where it regulates gene networks that control neuronal excitability. This is consistent with prior studies showing dense miR-206 expression in the cerebellum^28,29^, but contrasts with reports of miR-206 expression in extra-cerebellar regions including PFC, hippocampus, and other forebrain and midbrain structures^25,26^. Due to their short sequence length, miRNA probes can potentially bind related miRNAs or homologous transcript regions, emphasizing the importance of stringent detection methods for defining the precise expression patterns of miRNAs. Using constitutive, conditional, and viral genetic approaches to manipulate miR-206 expression in cerebellar PCs, we further demonstrate that miR-206 bidirectionally regulates PC firing dynamics and sensorimotor gating. Loss of miR-206 increased spontaneous tonic firing of PCs and impaired PPI, whereas elevated miR-206 expression shifted PCs toward high-frequency burst firing and exerted opposing effects on PPI depending on endogenous miR-206 expression levels. Together, these findings identify miR-206 as a key regulator of PC firing state and cerebellar-mediated sensorimotor gating behavior.

miRNA signaling plays essential roles in the development, maturation, and survival of cerebellar PCs^61–64^. A recent study using antisense sponge-mediated knockdown of miR-206 in PCs reported profound abnormalities in PC dendritic morphogenesis and climbing fiber innervation^28^. In contrast, we found that miR-206 is dispensable for PC specification, dendritic morphogenesis, survival, and cerebellar-dependent motor coordination. This discrepancy may reflect fundamental differences between sponge-mediated miRNA sequestration and precise genetic deletion approaches. miRNA sponge constructs can bind related miRNAs sharing homologous seed sequences, increasing the likelihood that they disrupt broader miRNA regulatory networks beyond miR-206 itself. Consistent with this possibility, we found that miR-206 deletion altered the composition of AGO2-associated miRNA populations in PCs, suggesting that perturbation of a single highly enriched miRNA can influence broader RISC-associated regulatory interactions. Nevertheless, precise genetic deletion of miR-206 produced only minimal structural and transcriptional alterations in PCs and other cerebellar cell populations, while enhancing miR-206 expression similarly caused no detectable loss or overt structural abnormalities of PCs. This indicates that miR-206 is not required for the development or maintenance of PCs and other cerebellar neurons under physiological conditions. Similarly, despite the highly restricted enrichment of miR-206 in both PCs and skeletal muscle^109,127–129^, constitutive miR-206 deletion caused no major impairments in baseline motor learning or coordination. These findings suggest that miR-206 does not function as a developmental determinant of cerebellar organization or neuromuscular-related motor programs but instead regulates specialized aspects of mature PC physiology.

To define the molecular mechanisms through which miR-206 regulates PC activity, we employed complementary transcriptomic and translational profiling approaches. snRNA-seq and Slide-seq transcriptional proflling revealed relatively subtle alterations in PCs and other cerebellar cell populations in miR-206-deficient animals, consistent with the largely preserved structural organization of the cerebellum. In contrast, HITS-CLIP and TRAP-seq translational proflling converged on regulatory networks gating ion channel function, synaptic transmission, and neuronal excitability. These findings suggest that miR-206 regulates post-transcriptional gene programs controlling the intrinsic excitability of PCs rather than broad developmental or cell fate pathways. miR-206 deletion also markedly altered AGO2-association of many other miRNAs in PCs, raising the possibility that miR-206 organizes broader miRNA regulatory architectures within the RISC complex. Thus, the functional consequences of miR-206 deletion may extend beyond direct repression of individual target transcripts to include remodeling of miRNA-mediated post-transcriptional regulatory networks in mature PCs.

Consistent with HITS-CLIP and TRAP-seq analyses identifying miR-206-regulated gene programs linked to neuronal excitability, miR-206 deletion increased spontaneous tonic firing of PCs. miR-206-deficient PCs also exhibited reduced HCN channel currents, which may contribute to increased PC intrinsic excitability^130,131^. However, spontaneous PC firing is regulated by diverse mechanisms, including ion channel activity, synaptic input, and neuromodulation^116,132–139^. Because our profiling detected numerous miR-206 targets with roles in each of these domains, miR-206 likely modulates PC excitability via distributed effects across multiple signaling pathways rather than a single target. Supporting a hyperexcitable state in PCs, immunofluorescence detection of the calcium-buffering protein CALB1 was enhanced after miR-206 deletion, perhaps in response to elevated intracellular calcium. *Calb1* transcript levels were not altered with the loss of miR-206, indicating post-transcriptional regulation or increased antibody epitope accessibility arising from calcium-dependent conformational changes in CALB1^34,136,138,140^.

Strikingly, augmenting miR-206 expression in mature PCs shifted neuronal activity from tonic toward high-frequency burst firing without altering the characteristic firing frequencies of either mode. These findings suggest that miR-206 regulates the probability that PCs occupy distinct intrinsic firing states rather than simply scaling overall firing frequency. As such, endogenous miR-206 levels are likely subject to tight regulatory control in PCs to maintain appropriate firing dynamics, with even modest perturbations in expression capable of altering PC firing state. More broadly, our findings raise the possibility that miRNAs regulate neuronal circuit function by biasing neurons between discrete functional states through coordinated post-transcriptional control of gene networks governing intrinsic excitability. Such mechanisms may represent an underappreciated cellular phenomenon in which disease-associated miRNAs shape circuit dynamics and behavior.

PCs exert powerful inhibitory control over cerebellar nuclei neurons and regulate cerebellar output through highly precise temporal firing patterns^80,144,145^. Transitions between tonic and burst firing modes are thought to profoundly influence cerebellar information processing, synchrony, and downstream motor and cognitive computations^144,145,133,146,147^. Altered tonic versus burst firing induced by miR-206 manipulations may therefore disrupt the timing and coordination of cerebellar output signals required for appropriate sensorimotor filtering. Consistent with this possibility, we observed evidence of altered transcriptional and metabolic states in cerebellar nuclei neurons following miR-206 deletion, including enrichment of pathways associated with mitochondrial stress and elevated cerebellar lactate levels. Determining how altered PC firing states reshape cerebellar nuclei activity and downstream brain circuitries will be an important direction for future studies.

Mice with whole-body miR-206 deficiency or selective ablation of miR-206 from PCs displayed impaired PPI, a conserved form of sensorimotor gating disrupted in schizophrenia and related neuropsychiatric disorders^52,54,125,126^. Increasing miR-206 expression in mature PCs impaired PPI in WT mice while rescuing PPI deficits in miR-206-deficient animals, indicating that both insufficient and excessive miR-206 signaling disrupt sensorimotor gating. These bidirectional behavioral effects parallel the opposing effects of miR-206 deficiency and augmentation on PC firing dynamics, suggesting that the distributions of tonic and burst firing states are critical for cerebellar regulation of sensorimotor filtering. Notably, the psychotomimetic NMDA receptor antagonist dizocilpine, which reliably disrupts PPI in rodents^148,149^, reduced cerebellar miR-206 expression. Whether this effect contributes to PPI deficits induced by dizocilpine is unclear, but it raises the possibility that perturbations that disrupt sensorimotor gating may act in part by altering cerebellar miR-206 signaling. Finally, miR-206 deficiency induced PPI deficits in only male mice, which is notable given the pronounced sex differences in risk of schizophrenia-spectrum disorders and in cerebellar contributions to neuropsychiatric phenotypes^150–153^.

Together, these findings identify miR-206 as a schizophrenia-linked regulator of PC firing state and sensorimotor gating but not cerebellum-mediated motor programs. More broadly, our results suggest that disease-associated and cell type-restricted miRNAs can shape neuronal activity patterns and behavior through coordinated post-transcriptional regulation of intrinsic firing dynamics in mature neural circuits.

## ACKNOWLEDGMENTS

We thank members of the Kenny laboratory for constructive criticism; Alicia Faruzzi Brantley for training in animal behavioral testing; Bridget Wicinski, Merina Varghese, and Patrick Hof for sharing iontophoretic injection expertise and equipment; Taleen Hania and Psychogenics for gait analysis, Anne Schaefer and Jessica Ables for technical advice on HITS-CLIP and TRAP-seq; Caroline Ménard and Scott Russo for providing tissue from mice that had undergone chronic social defeat stress; Kristin Beaumont and Robert Sebra for Genomics Core sequencing services; and Isaac Marin-Valencia and Manuel Gonzalez-Rodriguez for Neurometabolomics Core services. We are grateful for funding from the National Institutes of Health (R01MH112168, DA053629, and DA047233), the Distinguished Investigator Grant Program from the Brain & Behavior Research Foundation (BBRF), and the Seaver Center for Autism Research.

This work was supported in part through the computational and data resources and staff expertise provided by Scientific Computing and Data at the Icahn School of Medicine at Mount Sinai and supported by the Clinical and Translational Science Awards (CTSA) grant UL1TR004419 from the National Center for Advancing Translational Sciences. Research reported in this publication was also supported by the Office of Research Infrastructure of the National Institutes of Health under award number S10OD026880 and S10OD030463. The content is solely the responsibility of the authors and does not necessarily represent the official views of the National Institutes of Health.

Figure schematics were created in https://BioRender.com and Adobe Illustrator.

## AUTHOR CONTRIBUTIONS

Conceptualization, M.P.H. and P.J.K.; methodology, M.P.H., M.I., J.W., J.E.E., A.K.F., A.M., G.F., and P.J.K.; software, M.P.H. and J.E.E.; investigation, M.P.H., M.I., J.W., and A.K.F.; writing, M.P.H. and P.J.K.; funding acquisition, M.P.H. and P.J.K.; resources, G.F. and P.J.K.; and supervision, G.F. and P.J.K.

## RESOURCE AVAILABILITY

### Lead contact

Further information and requests for resources, reagents, or code should be directed to and will be fulfilled by the lead contact, Dr. Paul Kenny.

### Materials availability

Novel genetically targeted mouse lines (*miR-206^flox/flox^*and *miR-206*^-/-^) and plasmids (pBUD-EGFP, pBUD-miR-206-EGFP, pBUD-miR-133b-EGFP, pLuc-control, pLuc-miR-206-sensor, pAAV-MCS-EGFP, and pAAV-miR-206-EGFP) generated in this study will be deposited to The Jackson Laboratory and Addgene. Other materials generated in this study are available from the lead contact upon reasonable request, pending institutional approval and material transfer agreements.

### Data and code availability

Single nuclei RNA-seq, Slide-seq, CLIP-seq, and TRAP-seq data generated in this study will be deposited in the Gene Expression Omnibus (GEO). Raw reads will be made available in the Sequence Read Archive (SRA).

This study used existing command-line, R, and Python packages, and does not report original code. All scripts used for analysis and figure generation will be made available at Github/mph270 and archived at Zenodo.

Any additional information required to reanalyze the data reported in this paper is available from the lead contact upon request

## STAR METHODS

### KEY RESOURCES TABLE

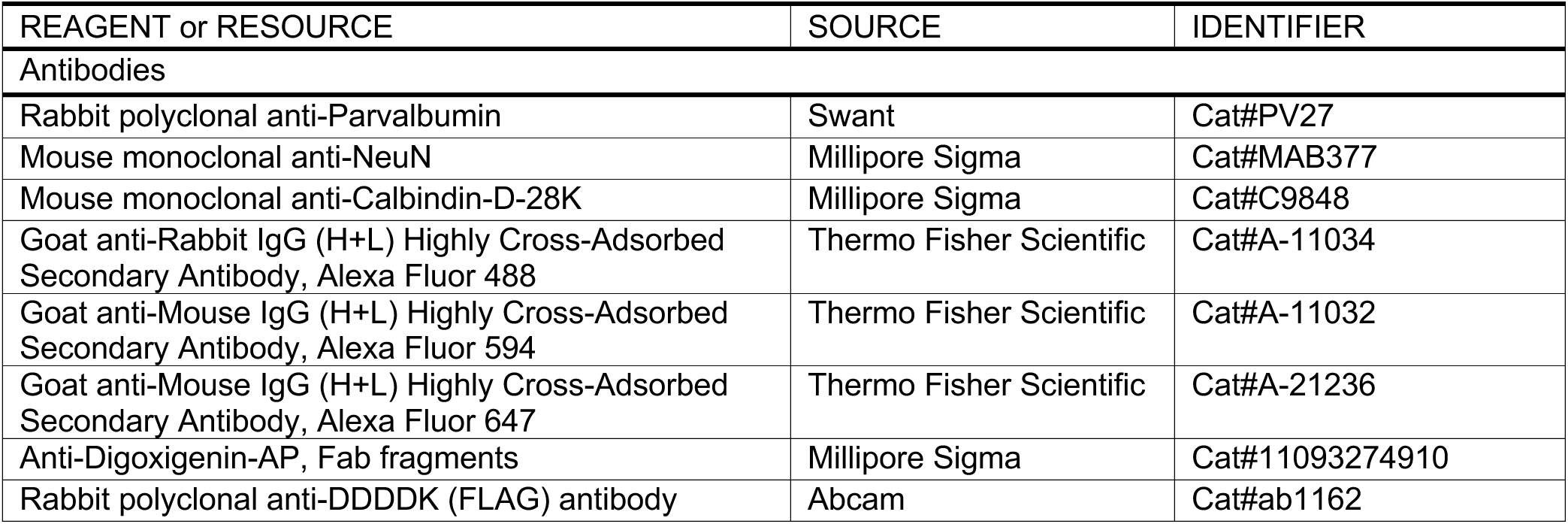
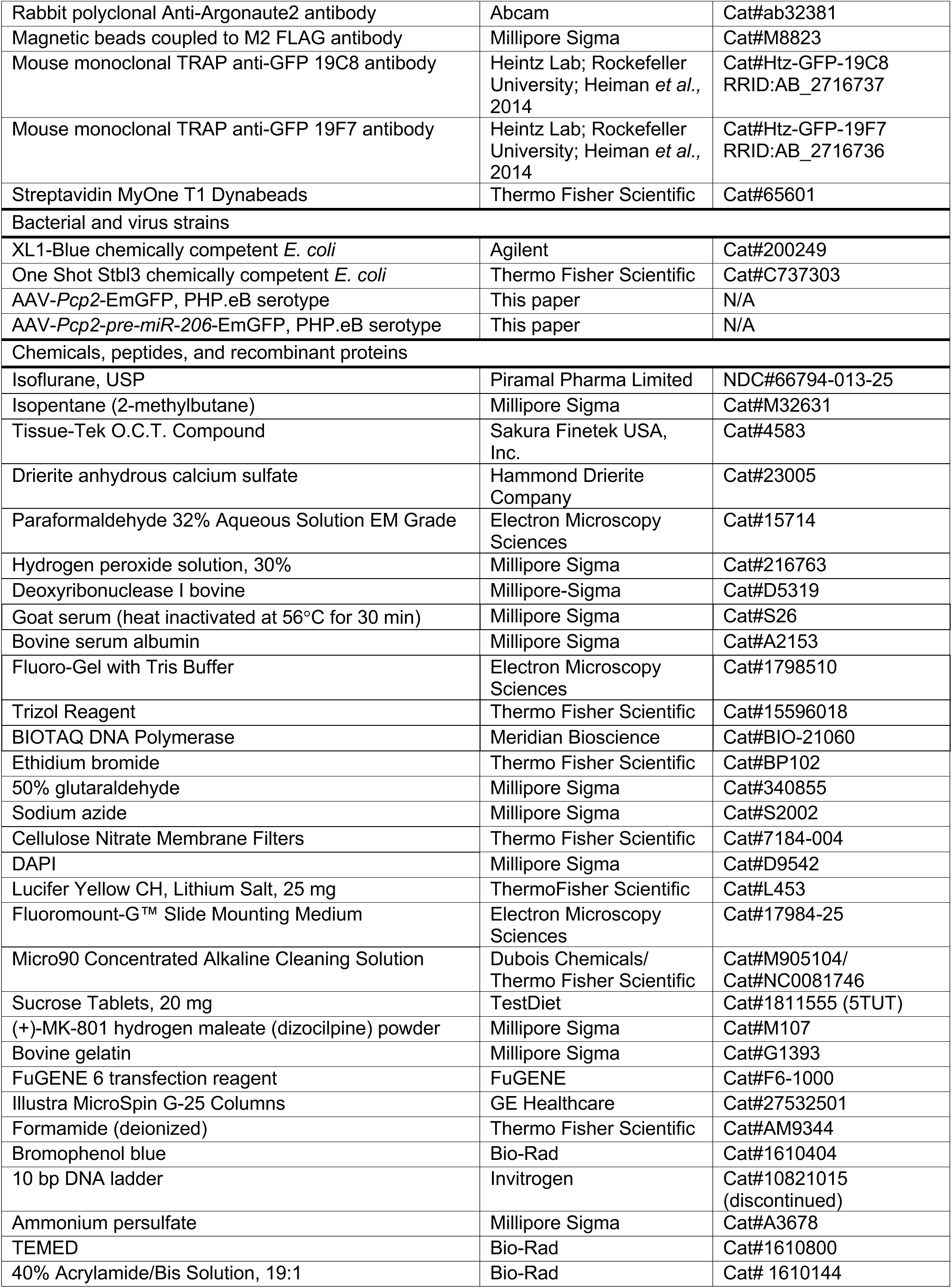
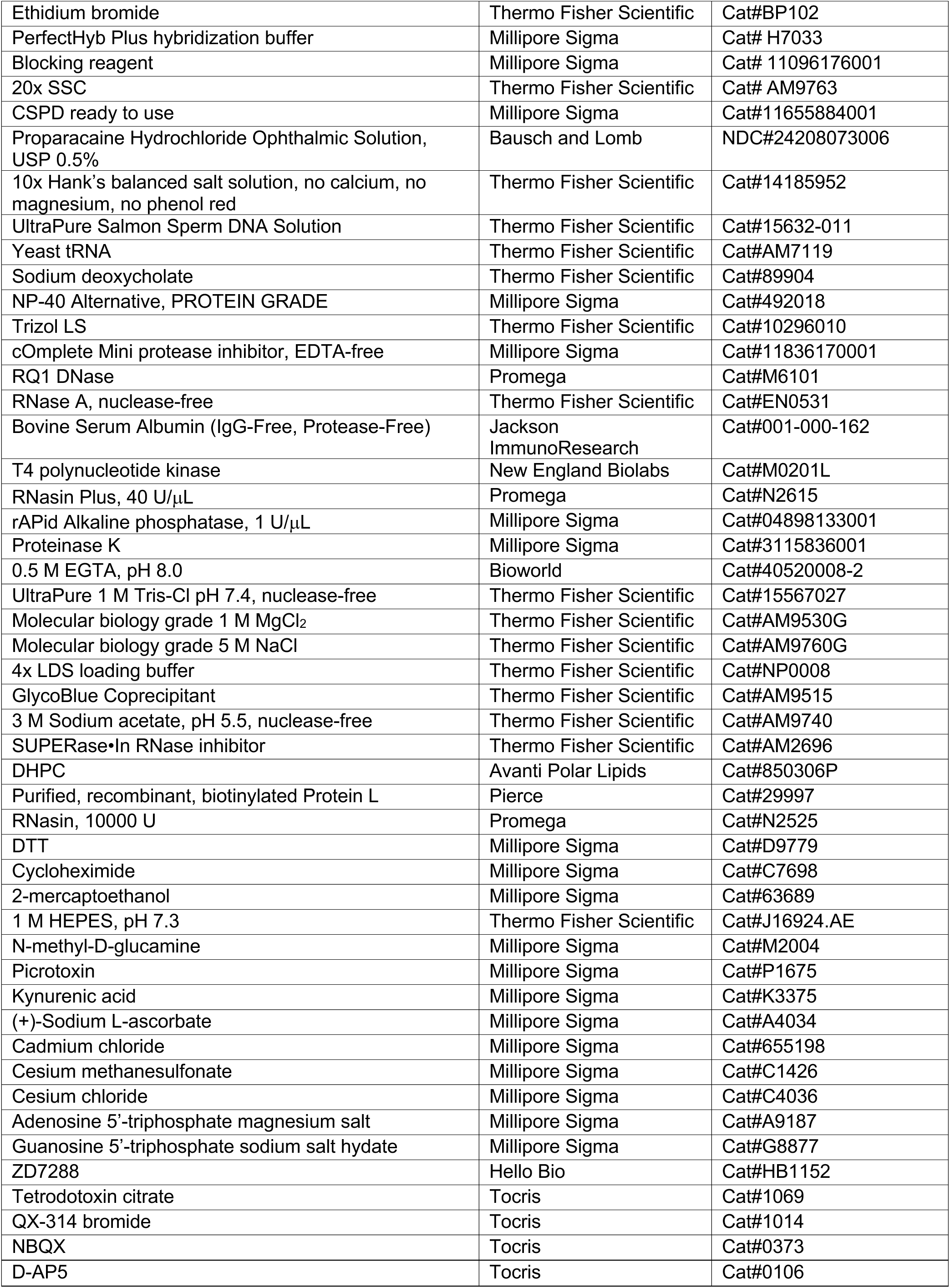
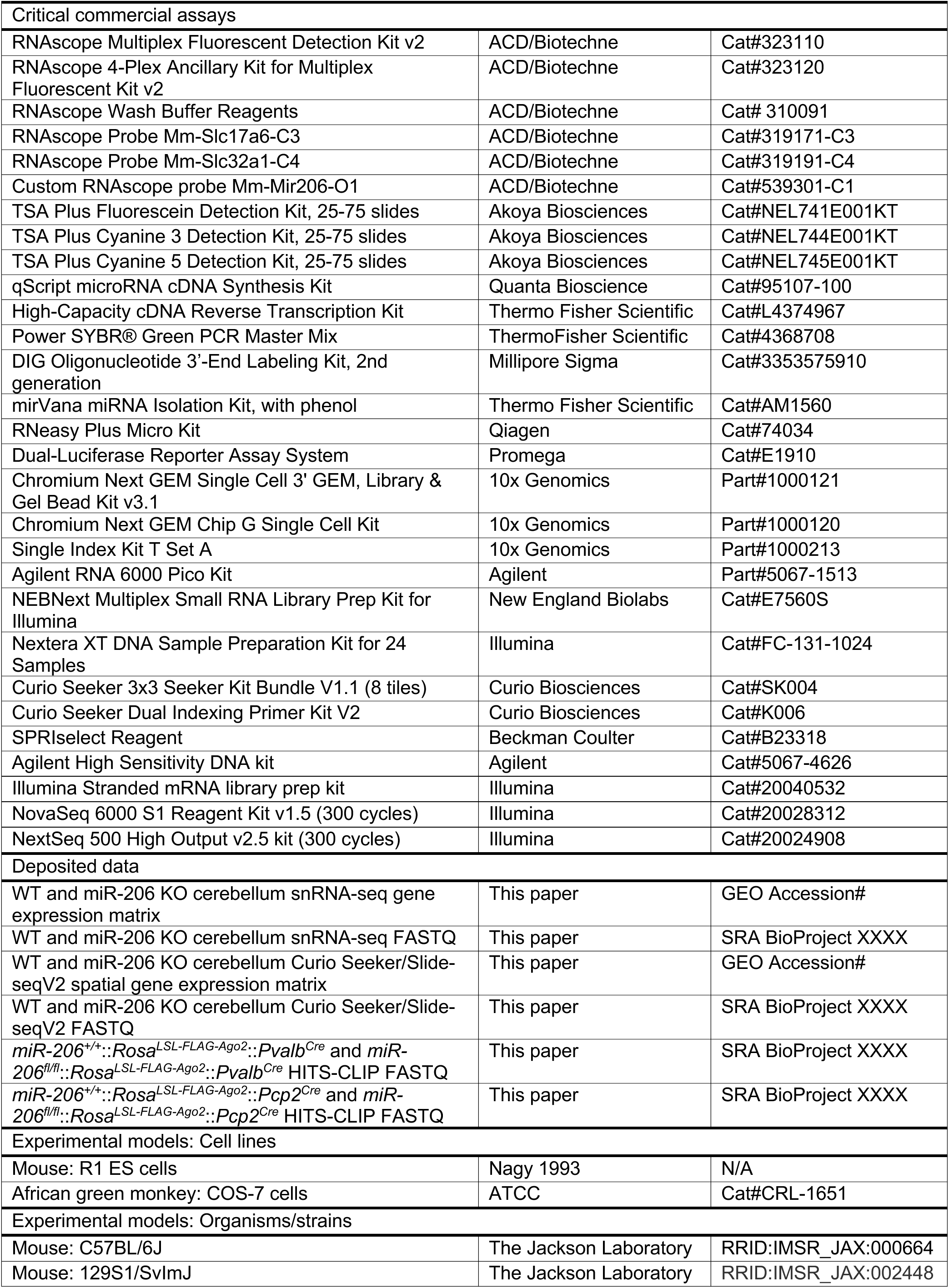
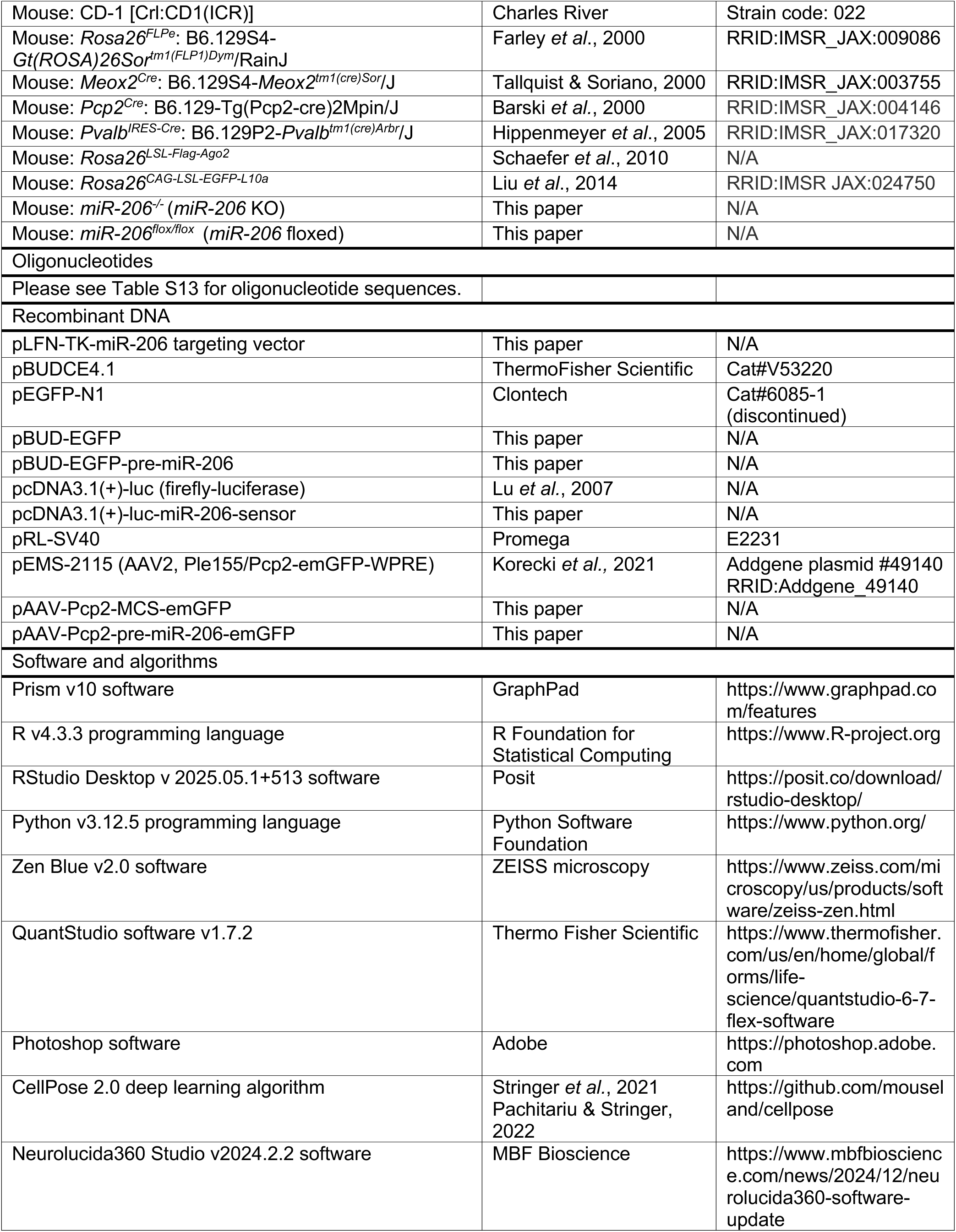
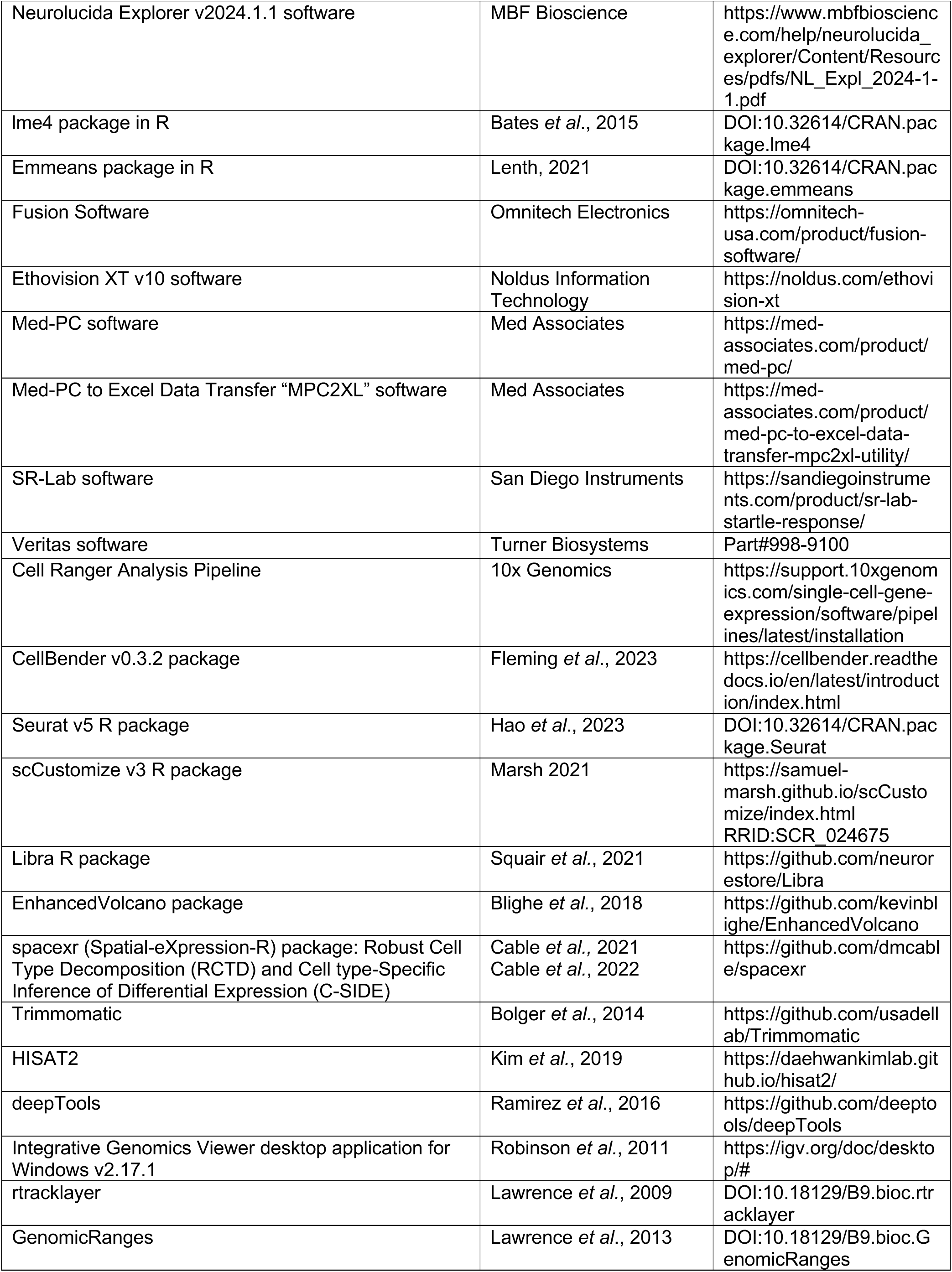
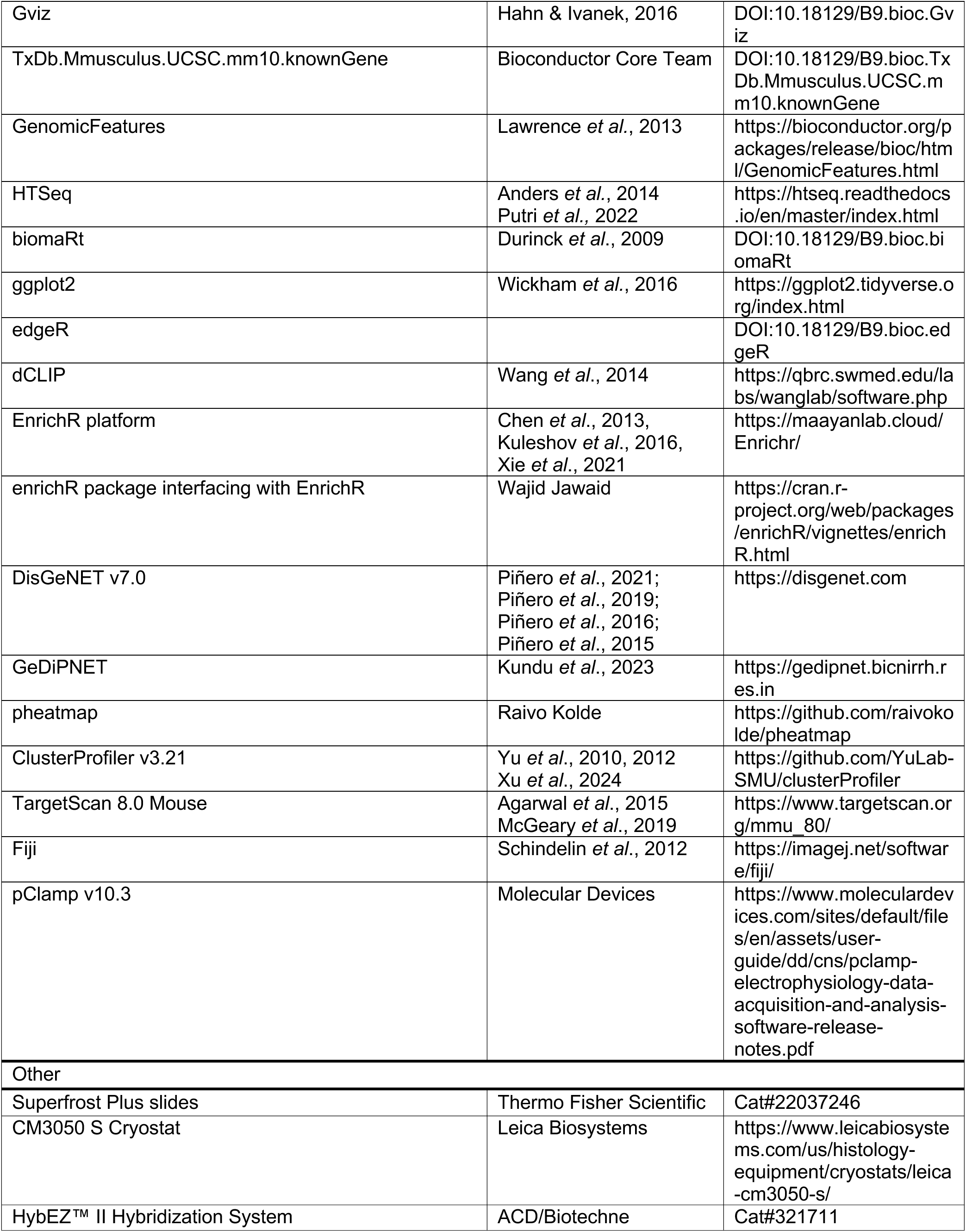
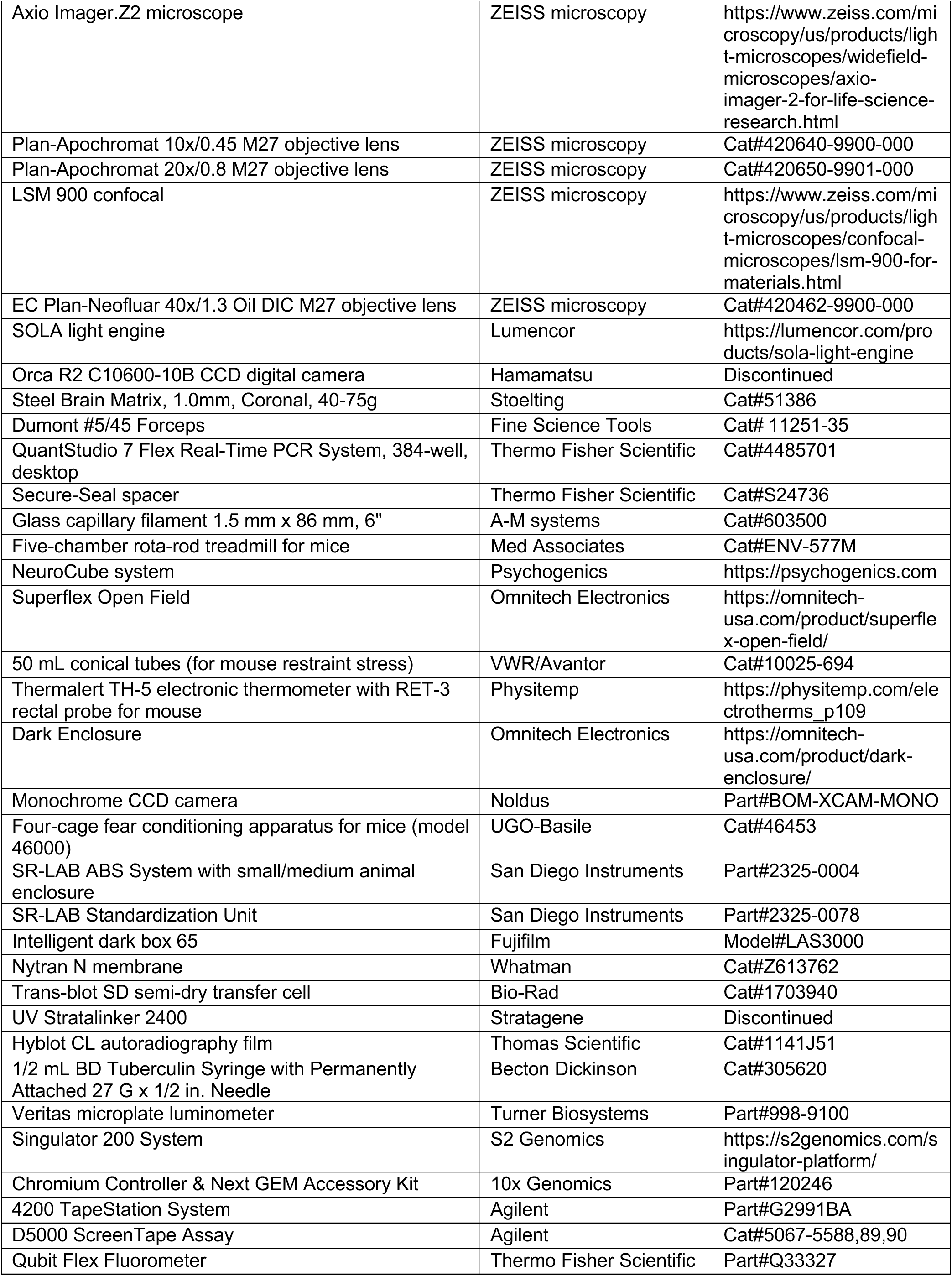
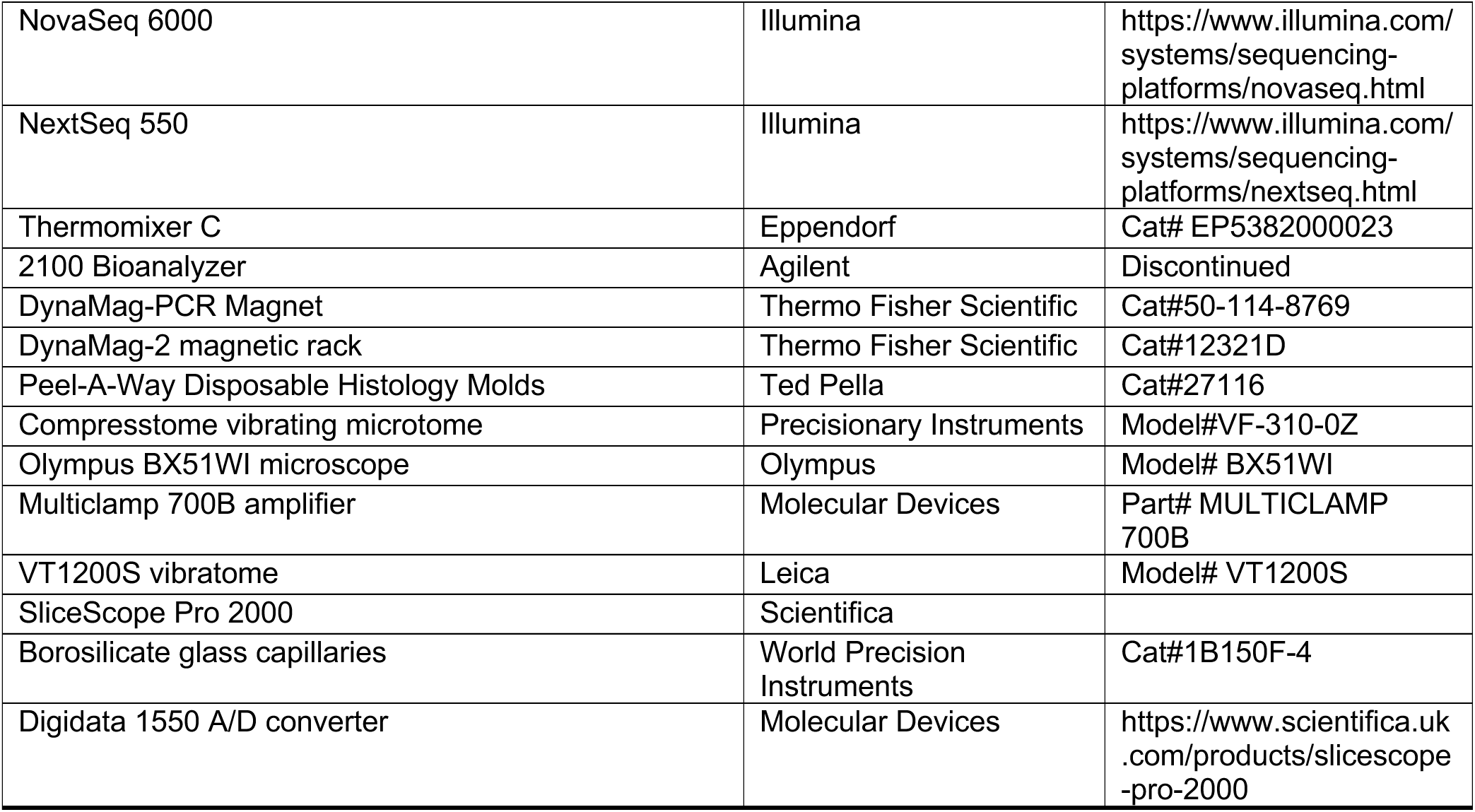

### EXPERIMENTAL MODEL AND STUDY PARTICIPANT DETAILS

#### Experimental animals

*miR-206*^flox/flox^ and *miR-206*^-/-^ mice were generated in this study and maintained on a C57BL/6J background. *Rosa26^FLPe^* (JAX #009086), *Meox2^Cre^* (JAX #003755), *Pcp2^Cre^/L7-Cre2* (JAX #004146), *Pvalb^IRES-Cre^* (JAX #017320), C57BL/6J (JAX #000664), and 129S1/SvImJ (JAX #002448) mice were imported from The Jackson Laboratory. *Rosa26^LSL-Flag^*^-*Ago2*^ mice were provided by A. Schaefer (Mount Sinai). CD-1 (Crl:CD1(ICR), #022)) mice were imported from Charles River. Mice were housed in an Association for Assessment and Accreditation of Laboratory Animal Care-approved vivarium on a 12:12 h light:dark cycle (lighted hours 07:00 to 19:00), with food and water ad libitum. All procedures were conducted in accordance with NIH guidelines and approved by the Institutional Animal Care and Use Committees at Duke University, The Scripps Research Institute Florida, and Icahn School of Medicine at Mount Sinai. All experiments were performed on adult male and female mice at least two months of age unless otherwise noted.

### METHOD DETAILS

#### Generation of miR-206 targeted mice

##### *miR-206* gene targeting strategy

Constitutive and conditional *miR-206* knockout mice were generated using previously described homologous recombination strategies^154,155^. A vector containing a 6-kb 5’ homology arm, a LoxP site, an *FRT*-flanked *pGK*-*Neo* cassette, the *miR-206* stem loop region, a second LoxP site, and a 1-kB 3’ homology arm was electroporated into R1 129/SvJ mouse embryonic stem (ES) cells^156^ (a gift of A. Nagy). Correctly targeted clones were identified by PCR screening and Southern blot of *HindIII*-digested genomic DNA. One confirmed clone was injected into C57BL/6J blastocysts, which were then implanted into pseudopregnant CD-1(ICR) females.

##### Neomycin resistance cassette removal and constitutive *miR-206* deletion

Male chimeras were bred to C57BL/6J females, and germline transmission was confirmed by PCR genotyping of tail DNA. *miR-206^Neo-flox^*offspring were crossed with *Rosa26^FLPe^* deleter mice^157^ to remove the neomycin resistance cassette, and *miR-206^flox/+^* offspring were confirmed by PCR. *miR-206^flox/+^* mice were then crossed with *Meox2^Cre^* deleter mice^158^ to generate the *miR-206* null allele, verified by PCR. *miR-206^flox/+^*and *miR-206^+/-^* mice were backcrossed at least 30 generations to C57BL/6J. Only FLP- and Cre-negative mice were used for breeding after the first backcross. All experimental cohorts were derived from *miR-206* heterozygote crosses.

##### Conditional *miR-206* deletion

Conditional deletion of miR-206 in Purkinje cells was achieved by crossing *miR-206^flox/flox^* mice with *Pcp2^Cre^/L7-Cre2* mice^159^. Conditional deletion of *miR-206* in parvalbumin-expressing neurons was achieved by crossing *miR-206^flox/flox^*mice with *Pvalb^IRES-Cre^* mice^160^. Effective deletion of miR-206 in the cerebellum was confirmed by quantitative RT-PCR of mature miR-206. Germline deletion of *miR-206* was detected in a small subset of *miR-206^flox/flox^::Pcp2^Cre^*offspring by PCR genotyping of tail DNA, consistent with prior reports of sporadic germline deletion for this Cre line^161^. Mice with detectable germline deletion of *miR-206* were excluded from experiments or breeding.

##### PCR genotyping

Genomic DNA was isolated from tissue using the HOTSHOT method^162^. Briefly, samples, were incubated in sodium hydroxide at 95°C for 1 h, followed by neutralization with Tris-HCl. PCR amplification was performed with BioTaq DNA polymerase (Meridian Bioscience) using 2 µl of genomic DNA per 25 µl PCR reaction. Thermocycling conditions were: 95°C for 5 min; 35 cycles of 95°C for 45 s, 60°C for 45 s, and 68°C for 45 s; 68°C for 10 min; and 4°C hold. Primer sequences are listed in the Key Resources Table.

#### RNAscope

A custom 20-ZZ-pair RNAscope^163^ probe was designed by Advanced Cell Diagnostics against a region of the mouse *7H4/miR-206/miR-133b* precursor RNA containing mature miR-206 that was previously validated by northern blotting^55^. Eight-week-old male and postnatal day 10 C57BL/6J mice were deeply anesthetized with isoflurane, decapitated, and their brains snap-frozen in isopentane in a metal beaker over dry ice. Brains were equilibrated in a cryostat (CM3050S, Leica Biosystems) for 1 h before sectioning, and mounted onto specimen holders using Tissue-Tek O.C.T. compound (Sakura Finetek USA). Fresh-frozen sagittal or coronal sections (14 μm) were collected onto Superfrost Plus slides (Fisher Scientific) and stored at -80°C in a sealed slide box containing anhydrous calcium sulfate dessicant (Hammond Drierite Company).

Slides were fixed in 4% paraformaldehyde in 1x nuclease-free PBS for 15 min at 4°C, then processed using the RNAscope Multiplex Fluorescent v2 Kit (Advanced Cell Diagnostics/Biotechne) according to the manufacturer’s fresh-frozen protocol, with modifications. After hydrogen peroxide treatment, slides were washed twice in 1x PBS and incubated with DNase I (Millipore-Sigma, 1:50 dilution in 10 mM Tris-Cl, 2.5 mM MgCl_2_, 0.5 mM CaCl_2_, pH7.5) to prevent hybridization of the *pre-miR-206* probe to genomic DNA. Slides were washed three times in 1x PBS, and probe hybridization was performed without protease digestion.

For 8-week-old samples, *pre-miR-206* (Mm-Mir206-O1, channel 1), *Vgat* (Mm-Slc32a1-C4), *Vglut1* (Mm-Slc17a6-C3), *Calb1* (Mm-Calb1-C3), and *Aldoc* (Mm-Aldoc-C4) probes (ACD/Biotechne) were developed with TSA-Cy3, -Cy5, and -fluorescein reagents (Akoya Biosciences) diluted 1:750 in TSA buffer (ACD/Biotechne). For P10 samples, *pre-miR-206* was developed with TSA-Cy3 followed by immunofluorescence staining of NeuN and Parvalbumin. After HRP blocker treatment (ACD/Biotechne), sections were washed once with RNAscope wash buffer (ACD/Biotechne) and twice with 1x PBS. They were then blocked for 1 h in 1x PBS, 5% normal goat serum, 2% bovine serum albumin, 0.1% Triton X-100 (all from Millipore Sigma), and incubated overnight at 4°C with rabbit anti-parvalbumin (Swant) and mouse anti-NeuN (Millipore Sigma) antibodies, both diluted 1:500 in blocking buffer. After three 10-min 1x PBS washes, sections were incubated for 3 h at room temperature with goat anti-mouse Alexa Fluor 647 and goat anti-rabbit Alexa Fluor 488 antibodies (Thermo Fisher Scientific), both at 1:500 in blocking buffer. Slides were washed (3 x 10 min, 1x PBS) and mounted with Fluoro-Gel with Tris buffer (Electron Microscopy Sciences).

Epifluorescence images were acquired using a widefield Axio Imager.Z2 microscope (ZEISS Microscopy) equipped with Plan-Apochromat 10x/0.45 and Plan-Apochromat 20x/0.8 objective lenses (ZEISS Microscopy), a SOLA light engine (Lumencor), and an Orca R2 C10600-10B CCD digital camera (Hamamatsu). Image acquisition, tiling, and stitching were performed with ZEN Blue software (ZEISS Microscopy). Confocal z-stacks of *pre-miR-20*6, *Vgat*, and *Calb1* signal in cerebellar lobule VI were acquired with an LSM 900 confocal microscope equipped with a 63x/1.4 N.A. Plan-Neofluar oil immersion objective and 405 nm diode, 488 nm argon, and 561 DPSS lasers (ZEISS microscopy). Z-stacks were acquired with a resolution of 1744 x 1744 pixels, a z-step of 0.23 µm, a pinhole of 1 Airy Unit, and optimized gain and offset settings. Images were acquired using ZEN Blue software (ZEISS microscopy).

#### RNA extraction, cDNA synthesis, and quantitative real-time PCR

Mice were deeply anesthetized with isoflurane and decapitated. Whole brain and cerebellum were dissected immediately after removal of the skull surface. Brain regions were isolated using Dumont #5/45 forceps (Fine Science Tools) from 1-mm coronal sections collected using a stainless-steel mouse brain matrix (Stoelting). Samples were snap-frozen in 1.5-mL tubes on dry ice, and total RNA was extracted using TRIzol reagent (Thermo Fisher Scientific) according to the manufacturer’s instructions.

For mRNA quantification, cDNA was synthesized from 1–2 µg of total RNA using the High-Capacity cDNA Reverse Transcription Kit (Thermo Fisher Scientific). For microRNA quantification, cDNA was generated by poly(A) tailing and reverse transcription of 500 ng–1 µg of total RNA using the qScript microRNA cDNA synthesis kit (QuantaBio). Individual mRNAs or microRNAs were quantified by real-time PCR using Power SYBR Green PCR master mix (Thermo Fisher Scientific) and custom intron-spanning mRNA primers or microRNA-specific primers paired with the PerfeCTa Universal PCR Primer (QuantaBio). Reactions were run on a QuantStudio 7 Flex Real-Time PCR System (Thermo Fisher Scientific) using the relative standard curve method. Expression levels of mRNAs and miRNAs were normalized to *Gapdh* and U6 small nuclear RNA, respectively.

#### miR-206 expression following acute or chronic restraint stress and forced swim stress

For restraint stress, male C57BL/6J mice were restrained in custom 50 mL conical tubes (Corning) with a 1/4-inch snout hole, 1/4-inch tail hole, and six 3/16-inch ventilation holes. Mice were restrained for 15 min for one session (acute restraint) or for two hours per day for 7 days (chronic restraint^164^). Mice were released and returned to their home cage after each restraint session. For forced swim stress, male C57BL/6J mice were placed in a 20-cm-diameter cylinder filled halfway with 24 ± 1°C water, and swimming was monitored for 10 min^165^. Co-housed mice underwent one swim stress per day for three days. Two hours after the onset of restraint (acute restraint) or 24 hours after the last stress session (chronic restraint and forced swim), mice were anesthetized and brain regions were dissected for quantitative RT-PCR of mature miR-206 as described above.

#### miR-206 expression following chronic social defeat stress (CSDS)

CSDS and social interaction/avoidance testing was performed as previously described^166,167^. Briefly, male C57BL/6J mice underwent 10 days of daily 10-min physical interaction with an unfamiliar CD-1 aggressor, followed by 24-h sensory contact across a perforated divider; unstressed controls were similarly rotated without aggressor exposure. The SI test was conducted 24 h after the final defeat, with locomotion and zone times recorded in the absence and presence of a novel CD-1 target. SI ratio (interaction time with target present/absent) was used to classify mice as susceptible (SS, ratio <1.0) or resilient (RES, ratio >1.0). 24 hours following behavioral testing, mice were anesthetized and brain regions dissected for quantitative RT-PCR of mature miR-206 as described above.

#### miR-206 expression following NMDA receptor blockade with dizocilpine

Male C57BL/6J mice were injected intraperitoneally with saline or 0.25 mg/kg dizocilpine (MK-801) (Sigma-Millipore) dissolved in saline once per day for 7 days. 24 hours following the last injection, mice were anesthetized and brain regions dissected for quantitative RT-PCR of mature miR-206 as described above.

#### Immunofluorescence

Mice were deeply anesthetized with isoflurane and intracardially perfused with 1x PBS, followed by 4% PFA in 1x PBS. Brains were dissected, post-fixed overnight in 4% PFA at 4°C, and cerebella sectioned at 50 μm thickness in the sagittal plane at ∼100 μm intervals using a vibratome (Vibratome 1000 PLUS, Vibratome). Floating sections were blocked in 1x PBS, 5% normal goat serum (Millipore Sigma), 2% bovine serum albumin (Millipore Sigma), 0.1% TX-100 (Millipore Sigma) for 1 h at room temperature and then incubated with mouse anti-CALB1 (Millipore Sigma; 1:1000 dilution) or rabbit anti-PV (Swant, 1:5000) in blocking buffer overnight at 4°C. Sections were washed three times for 20 min each in 1x PBS, incubated with goat anti-mouse Alexa Fluor 594 (Thermo Fisher Scientific; 1:500 dilution) in blocking buffer for 3 h at room temperature, then washed three times in 1x PBS. Sections were stained with DAPI (Millipore Sigma; 1:10,000 dilution) in 1x PBS for 10 min, washed briefly in 1x PBS, and mounted on Superfrost glass slides (Thermo Fisher Scientific) with Fluoro-Gel with Tris buffer (Electron Microscopy Sciences).

For CALB1 immunoreactive cell counts, six midline (cerebellar vermis) sections were stained per animal. For PV immunoreactive cell counts, 4 sections (mPFC), 16 sections (dorsal hippocampus), and 10 sections (GPe) were stained per animal. Tiled epifluorescence images were acquired using an Axio Imager.Z2 microscope (ZEISS microscopy) and 40x objective as described for RNAscope imaging. For CALB1and DAPI fluorescence intensity measurements of Purkinje cells, one medial and one lateral section were stained per animal, and six confocal images of lobule VI Purkinje cells were acquired per section using an LSM 900 confocal microscope equipped with a 40x/1.3 N.A. Plan-Neofluar oil immersion objective and 405 nm diode and 561 DPSS lasers (ZEISS microscopy). Z-stacks were acquired with a resolution of 1024 x 1024 pixels, a z-step of 1 µm, a pinhole of 1 Airy Unit, and optimized gain and offset settings. Images were acquired using ZEN Blue software (ZEISS microscopy).

#### Lucifer Yellow iontophoretic filling of Purkinje cells

Lucifer Yellow iontophoretic cell filling was performed as previously described^168^, with minor modifications. Mice were deeply anesthetized with isoflurane and transcardially perfused with 1% PFA in phosphate buffer (PB, pH 7.4) for 2 min, followed by 4% PFA and 0.125% glutaraldehyde in PB for 10 min. Brains were immediately dissected, post-fixed overnight at 4°C in 4% PFA and 0.125% glutaraldehyde in PB, and stored in phosphate-buffered saline (PBS) with 0.1% sodium azide at 4°C.

250-µm sagittal sections were collected throughout the cerebellar vermis using a vibratome (Vibratome 1000 PLUS, Vibratome). Sections were incubated in 250 ng/ml DAPI (Millipore Sigma) for 5 min to visualize the Purkinje cell layer, mounted on nitrocellulose membrane filters, and immersed in ice-cold PB. Individual Purkinje cells were iontophoretically injected with 5% Lucifer Yellow dye (Thermo Fisher Scientific) in distilled water under a direct current of 3–8 nA until the dye filled the distal dendritic arbors. Cells were spaced sufficiently to avoid dendritic overlap. Sections were mounted with Fluoromount-G (Southern Biotech) between Secure-Seal spacers (Thermo Fisher Scientific) on Superfrost plus slides (Thermo Fisher Scientific).

Lucifer Yellow-filled Purkinje cells were imaged using an LSM900 confocal microscope (ZEISS microscopy) equipped with a 40x/1.3 N.A. Plan-Neofluar oil immersion objective and an argon laser (488 nm excitation). Confocal z-stacks were acquired with a resolution of 1024 x 1024 pixels, a z-step of 0.31 µm, a pinhole of 1 Airy Unit, and optimized gain and offset settings. Images were acquired using ZEN Blue software (ZEISS microscopy).

#### Behavioral Assays

All behavioral testing was conducted during the active (dark) phase of the light cycle. Unless otherwise noted, mice were handled for 30 seconds per day for three consecutive days prior to testing and allowed to acclimate to the testing room for 1 h before each session. Behavioral equipment was cleaned using Micro-90 (Dubois Chemicals). Mice were tested between 3–6 months of age.

##### Prepulse inhibition of the acoustic startle response

Prepulse inhibition of the acoustic startle response (PPI) was tested in an SR-Lab startle response system (San Diego instruments) equipped with a cylindrical acrylic mouse enclosure mounted on a platform with piezoelectric detection of startle responses, all inside an isolation cabinet^169^. The SR-Lab Standardization Unit (San Diego Instruments) was used to calibrate signal from the piezo-electric detector to an average amplitude of 725 mV. White noise production was calibrated using a digital sound meter in the center of the chamber. On the day before testing, mice were acclimated to the enclosure for 5 min with continuous 70 dB background white noise.

On the test day, mice were acclimated for 5 minutes before presentation of a block of six 40-ms 120 dB broadband white noise pulse-alone (startle) trials; a block of 52 pseudorandomized trials consisting of 120 dB pulse-alone trials, 20-ms 74, 78, or 86 dB pre-pulse stimuli presented 100 ms prior to the 120 dB startle pulse, and no stimulus trials; and a final block of six 120 dB pulse-alone trials. White noise was used to circumvent age-related high frequency hearing loss in C57BL/6J^170,171^. 70 dB background white noise was played during inter-trial intervals that were pseudorandomized and ranged from 8-23 s. The whole-body flinch response to the startle pulse was recorded as 65 consecutive 1-ms voltage readings beginning from pulse stimulus onset. The session contained a total of 64 trials and lasted approximately 23 minutes.

##### Acoustic startle response threshold

The threshold of the acoustic startle response was determined as previously described^172^. Mice were habituated to and tested in the SR-Lab startle response system as above, but instead of the PPI protocol, they were presented with pulse-alone stimuli ranging from 65 to 120 dB. The first block consisted of six 40-ms broadband bursts of 120-dB broadband white noise. The second block consisted of 40-ms broadband bursts at 65, 69, 73, 77, 85, 90, 100, 110, or 120 dB, presented in a pseudorandom order for a total of six times for each dB level. The entire session contained 60 trials and lasted 20 min.

##### Rotarod

Motor coordination and learning were assessed using the accelerating rotarod task^173,174^. Mice underwent three trials in a single day on an ENV-577M five-compartment accelerating rotarod (Med Associates). Mice were sequentially placed on the rod rotating at a steady velocity of 4 rpm. Once all mice were loaded, the rod rotation accelerated from 4 to 40 rpm over 5 min. Latency to fall was defined as the duration the mouse remained on the road before falling or failing maintain position. The intertrial interval was at least 30 min to minimize fatigue. Up to five mice from the same home cage were tested simultaneously, with WT and KO animals counterbalanced across compartments.

##### Gait assessment

Mice were tested in a NeuroCube system (Psychogenics), which employs computer vision to detect changes in gait geometry and gait dynamics in rodents^175,176^. Mice were placed in the apparatus and allowed to walk freely for a 5-min session.

##### Open Field and home cage activity

Spontaneous locomotor activity was assessed in the open field as previously described^155,177–179^. Mice were placed in a 16 x 16-inch acrylic arena inside a Superflex automated infrared detection system enclosed in an environmental control chamber (Omnitech Electronics). Horizontal and vertical beam breaks were recorded using Fusion software (Omnitech electronics) in order to calculate distance traveled and vertical activity. Sessions lasted 60 min without illumination. For acute stressor testing, activity was recorded for 30 min, mice were briefly restrained by scruffing, and then returned to the chamber for an additional 90 min.

Home cage activity was monitored by placing individual mice in cages containing fresh corn cob bedding within the Superflex detection system and environmental control chamber. Mice had free access to food and water and were left undisturbed for 48 h under their normal light cycle. Fusion software was used to control the light cycle of the chambers.

##### Stress-induced hyperthermia

Stress-induced hyperthermia was measured as described previously^180^. Group-housed mice were weighed and separated into individual cages. On the following day, each mouse was gently restrained horizontally, and rectal temperature was measured using a Thermalert TH-5 electronic thermometer with a RET-3 rectal probe (Physitemp). The probe was coated with sterile silicone oil, inserted 2 cm into the rectum, and held until a stable temperature was recorded (∼ 20 seconds, *T_1_*). The mouse was returned to its cage, and the rectal probe was cleaned with 70% ethanol. After 10 minutes, a second measurement of rectal temperature was taken (*T_2_*) to determine the extent of stress-induced hyperthermia.

##### Tail Suspension

The tail suspension test^181^ was conducted in a black acrylic four-chambered apparatus with a white acrylic backing permeable to back-lit infrared light (custom-built by the Scripps Research Florida Behavior Core). A plastic tail guard was placed over each mouse’s tail to prevent tail climbing, and the tail tip was taped to a hook under the chamber roof. Mice were positioned on stands until all mice were in the apparatus, after which the stands were gently removed, leaving the mice suspended by their tails. Mice were recorded and tracked for 6 minutes, and immobility duration was calculated using Ethovision XT tracking software (Noldus Information Technology).

##### Dark light emergence

Anxiety-like behavior was assessed in light-dark boxes as previously described^94,182^. Each 16 x 16-inch acrylic open field chamber was divided into a dark enclosure covering one half and an illuminated area in the other, with a gate allowing free movement between the two sides (Omnitech Electronics). Chambers were paired with a Superflex automated infrared detection system within an environmental control chamber (Omnitech Electronics). Mice were initially placed in the dark chamber, and the gate barrier was lifted after 10 seconds to start the test. Mice were allowed to explore both compartments freely for 10 minutes. Horizontal and vertical beam breaks in the light and dark zones were recorded with Fusion software (Omnitech electronics) and used to quantify the total duration and number of entries in each compartment, as well as latency to enter the light side.

##### Hyponeophagia

The hesitation of mice to ingest a novel palatable food was used as another measure of anxiety-like behavior^183^. Mice were food-restricted for 18 hours prior to testing. Twenty sucrose pellets (20 mg, TestDiet) were placed in the center of a 40.5 cm x 20 cm x 22 cm acrylic chamber with white walls and floor illuminated with white light. Each mouse was allowed to explore the chamber freely for 30 minutes while being recorded with an overhead camera and tracked using Ethovision XT software (Noldus). The latency to eat a sucrose pellet continuously for at least 2 sec, the total number of food pellets consumed, and the number of fecal boli produced were manually scored.

##### Fear conditioning and extinction

Pavlovian fear conditioning and extinction tests were adapted from established methods^184,185^, using the Fear Conditioning System for mice (Ugo Basile). Mice were recorded and tracked with overhead cameras and Ethovision XT software (Noldus). Training was carried out under 70 lux white light illumination in a rectangular chamber with electrified metal grid floor above a removable tray containing clean corn cob bedding, all inside an isolation cubicle with a ventilation fan turned on. Mice were transported individually to the testing room in home cages with clean bedding. Overhead white light and background white noise were both on in the testing room, and mice were handled with latex gloves. The training protocol consisted of a 150-s period of acclimation to the chamber; 30 s of an 80 dB, 3 kHz tone, with 0.6 mA shock concurrent with the final 2 s of tone presentation; a 60 s interval; a second 30-s tone/shock pairing; a 90 sec interval; a third 30-s tone/shock pairing; and a 30 s interval before the end of the trial. Chambers were cleaned with 70% ethanol between sessions.

For contextual fear memory retrieval and extinction testing, 24 and 48 hours after training, mice were placed into same rectangular arena and same context in which training was conducted (transport in home cages, white light on in arena, white ventilation fan on, white light and white noise on in testing room, handling using latex gloves, arenas cleaned with 70% ethanol). The mice were recorded and tracked for 5 min without shocks or tone.

For assessment of cued fear memory retrieval and extinction, mice were tested at least three hours after each contextual test. Mice were placed in a novel context consisting of a custom-built PVC cylindrical arena with clear walls and white floor. Other altered contextual cues included transport in empty plastic containers, infra-red light on in arena, ventilation fan off, red light on and white noise off in testing room, handling using nitrile gloves, arenas cleaned with Micro-90, and an orange scent (orange essence diluted in water) placed in a plastic weight boat inside the enclosure. After a 3-min acclimation to the arena, mice were presented with 39 trials of a 20-s, 80 dB, 3 kHz tone without shock. The inter-trial interval was 20 s.

For each test, distance traveled and activity were monitored using Ethovision XT. Freezing was defined as inactivity lasting at least 1 s and was verified by visual inspection of the video recording. Percentage freezing duration was calculated for each 30-s, 1-min, and 20-s time bin for training, contextual, and cued tests, respectively.

##### Sociability

Mouse sociability was assayed using an automated three-chambered social approach task^186–188^. The apparatus was a three-chambered 60 x 40.5 x 22 cm rectangular arena with clear acrylic walls, a white matte bottom, and passages fitted with removable sliding doors between the chambers (Noldus). 10 x 20 cm cylindrical restraints (Noldus) were used as novel objects and for housing novel “stimulus” mice. Testing was conducted under red light during the active phase, with background white noise in the room. The 129S1/SvImJ strain (The Jackson Laboratory #002448) was used as the strain of stimulus mice due to a more passive nature and lowered tendency to climb the bars of the restraints. Stimulus mice were habituated to the restraints for 15 min per session, three sessions per day, for two days before the sociability assay.

On the day of testing, the experimental or “test” mouse (WT or miR-206 KO) was placed in the center chamber and the passage doors removed to allow the mouse to explore and acclimate to the entire empty apparatus. After 5 min, the doors were closed, and the mouse was returned to the center chamber. An age- and sex-matched stimulus mouse was placed inside a restraint in one chamber, and an empty restraint was placed in the other. The passage doors were removed, and the test mouse was allowed to explore and interact with the empty or novel mouse-containing restraint for 10 minutes. The test mouse was recorded and tracked using an overhead camera and Ethovision XT (Noldus). The duration of the test mouse in each chamber and the duration of the test mouse’s nose point detected within a circular “investigation” zone two cm adjacent to the restraint perimeter were calculated in Ethovision XT. Stimulus mice were not used on successive trials, and their placement in the restraints was counterbalanced across the left and right chambers.

#### miR-206 expression constructs

##### Dual promoter miR-206 expression plasmid cloning and transfection

GFP control and miR-206/GFP expression vectors were constructed from the pBUDCE4.1 dual promoter vector (ThermoFisher Scientific). GFP from pEGFP-N1 (Clontech) was cloned into *HindIII/XbaI* restriction sites downstream of the CMV promoter. The miR-206 stem-loop region was amplified by PCR of genomic DNA and cloned into *XhoI/BglII* restriction sites downstream of the EF1-α promoter. Correct clones were validated by colony PCR and Sanger Sequencing. COS-7 cells were grown in 10 cm plates coated with 0.1% gelatin (Millipore Sigma). When cells reached 60% confluence, they were transfected with 6 ug of plasmid DNA using 18 µL of Fugene 6 transfection reagent (FuGENE). After 48 hours, green fluorescence was visualized, cells were scraped in 1x PBS and centrifuged, and the pellet was frozen at -80°C. Small RNA-enriched RNA was prepared from frozen pellets using the miRVana miRNA isolation kit (Thermo Fisher Scientific), according to manufacturer’s instructions.

##### LNA-DIG miR-206 probe labeling

A miR-206 antisense DNA oligonucleotide containing locked nucleic acid (LNA) modifications (Integrated DNA Technologies) was 3’ end-labeled with digoxigenin (DIG) using the DIG oligonucleotide 3’ end-labeling kit (Millipore Sigma), unincorporated DIG was removed using Microspin G-25 columns (GE Healthcare), and labeling efficiency was determined according to the manufacturers’ instructions.

##### miR-206 northern blotting validation of miR-206 expression

miRNA northern blotting was carried out as previously described^129,189^. Briefly, 1 µg of small RNA-enriched RNA was denatured in 2x loading buffer (80% formamide, 10 mM EDTA pH 8, bromophenol blue) at 70°C for 10 min and chilled on ice. 10 μl of a 10 bp DNA ladder (Invitrogen) was similarly denatured in 2x loading buffer. RNA samples and ladder were separated on a denaturing 15% polyacrylamide gel (50% urea, 0.5x Tris-borate-EDTA (TBE) buffer, 0.1% ammonium persulfate and 0.05% TEMED) in 0.5x TBE buffer at 200 V. The gel was stained with 4 μg/ml ethidium bromide in 0.5x TBE for 10 mins. 78-nucleotide tRNA and 120-nucleotide 5S rRNA bands were imaged on an LAS3000 intelligent dark box 65 (Fujifilm) and used as loading controls. RNA was transferred in 0.5x TBE to a Nytran N nylon membrane (Whatman) using a semi-dry transfer apparatus (Bio-Rad) at 15 V for 75 min. The membrane was air dried for 10 min and RNAs crosslinked to the membrane upon exposure to 1200 mJ of UV light in a UV Stratalinker 2400 (Stratagene).

The membrane was incubated with 0.1 nM of miR-206 antisense DIG-LNA probe in PerfectHyb Plus (Millipore Sigma) overnight at 55°C with agitation. After probe hybridization, the membrane was washed with 2x SSC + 0.1% SDS for 30 min at room temperature followed by 0.5x SSC + 0.1% SDS for 30 min at 45°C. It was then washed in washing buffer (Tris-buffered saline + 0.3% Tween) and blocked in 1x blocking solution (Millipore Sigma, 10x stock diluted in washing buffer) for 30 min at room temperature, followed by incubation with anti-DIG antibody conjugated to alkaline phosphatase (Millipore Sigma) diluted 1:10,000 in 1x blocking solution for 1 h at room temperature. The membrane was washed three times with washing buffer, 15 min each, and finally washed twice with 0.1 M Tris-Cl pH 9.5 + 0.15 M NaCl, 15 min each. Alkaline phosphatase signal was developed with CSPD, ready-to-use (Millipore Sigma) and luminescence at 477 nm was detected using Hyblot CL autoradiography film (Thomas Scientific).

##### miR-206 luciferase sensor cloning

A modified version of pcDNA3.1(+) containing a firefly luciferase gene with a downstream multiple cloning site for cloning a 3’UTR region was a gift of Yun Lu^190^. Oligonucleotides containing two regions antisense to miR-206 separated by a *BamHI* site and flanked by *EcoRI* and *XhoI* sites were 5’ phosphorylated and annealed, then cloned into *EcoRI* and *XhoI* sites downstream of luciferase to create a miR-206 sensor.

##### Luciferase assay validation of miR-206 expression

COS-7 cells were grown in 24 well plates coated with 0.1% gelatin (Millipore Sigma). When cells reached 60% confluence, they were transfected in triplicate with 0.05 μg of pcDNA-luc constructs, 0.5 ng of pRL expressing *Renilla* luciferase, 0.5 μg of pBUD-GFP or pBUD-GFP-miR-206 and 0.9 μl of FuGENE 6 transfection reagent (FuGENE). 24 h after transfection, the relative firefly/*Renilla* luciferase levels were analyzed using the Dual-Luciferase Reporter Assay System (Promega), according to the manufacturer’s instructions. Cells in each well were lysed with 100 μl of lysis buffer for 15 min at room temperature. 5 μl of lysate was added to the middle of each well of a 96-well white Lumitrac 200 assay plate (USA Scientific), and firefly and Renilla luciferase activity were detected using a Veritas microplate luminometer and software (Turner Biosystems).

##### *Pcp2*-GFP and *Pcp2*-*pre-miR-206*-GFP AAV viral vector cloning and packaging

The pEMS2115 plasmid containing a Purkinje cell selective promoter (Ple155, *PCP2*) driving expression of EmGFP was a gift from Elizabeth Simpson^191^ (Addgene #49140). EmGFP was excised by restriction digestion with *NotI* enzyme and replaced with the following fragmentGENE DNA (Genewiz/Aventa): (1) *NotI-BamHI-XhoI-NheI*-Kozak-EmGFP-*SalI-NotI* to generate a new *Pcp2-*GFP control plasmid containing a multiple cloning site (MCS) and Kozak sequence upstream of GFP and (2) *NotI-BamHI-XhoI*-*pre-miR-206*-*NheI*-Kozak-EmGFP-*SalI-NotI* to generate a *Pcp2-pre-miR-206*-GFP plasmid expressing miR-206 and GFP. The miR-206 stem-loop sequence was identical to that of the pBUD-*pre-miR-206*-GFP plasmid validated *in vitro* as described above. Correct clones were identified by colony PCR using (1) Pcp2-F and MCS-R primers and (2) Pcp2-F and 206-R primers, followed by Sanger sequencing using Pcp2-F and WPRE-R primers. 100 μg of each plasmid was sent to the Duke Viral Vector Core for packaging into AAV viral particles with the PHP.eB serotype. Purified viral titers were 1.93x10^13^ vg/mL for *Pcp2*-GFP and 1.05x10^13^ vg/mL for *Pcp2-pre-miR-206-*GFP.

##### Intravenous AAV viral delivery

PHP.eB AAV-*Pcp2-*GFP (AAV-control) and AAV-*Pcp2-*miR-206-GFP (AAV-miR-206) were administered intravenously as previously described^192,193^. One-month-old mice were anesthetized with isoflurane (3 to 5% induction; 1 to 3% maintenance) and placed on a heat pad. 1x10^11^ vg of virus was diluted in 70 µL of sterile 1x DPBS per mouse and delivered by injection into the left retro-orbital sinus using a 27-gauge 0.5 ml tuberculin syringe (Becton Dickinson). Proparacaine hydrochloride analgesic solution (Bausch and Lomb) was applied to the eye and mice were monitored until fully recovered from anesthesia (1–2 min).

#### Single nuclei RNA-seq

Six-month-old male WT and miR-206 KO mice (n = 4 per genotype) were deeply anesthetized with isoflurane and decapitated. Intact cerebella were removed, flash frozen in liquid nitrogen, and stored at -80°C. Tissue was dissociated and nuclei from each sample isolated using the S2 Singulator Platform Low Input Protocol according to the manufacturer’s guidelines (S2 Genomics). Nuclei were processed with the Chromium Next GEM Single Cell 3’ v3.1 kit (10X Genomics). An estimated 10,000 nuclei were loaded onto each channel with a targeted cell recovery of 6000 nuclei/sample. Captured mRNA then underwent reverse transcription followed by cDNA amplification and fragmenting, end-repair, poly A-tailing, adapter ligation, and 10X-specific sample indexing according to the manufacturer’s protocol. Libraries were quantified using TapeStation (Agilent) and Qubit (Thermo Fisher Scientific) analysis. Libraries were sequenced in paired end mode on a NovaSeq instrument (Illumina) targeting a depth of 50,000-100,000 reads per cell.

#### Curio Seeker / Slide-seqV2

Two-month-old male WT and miR-206 KO mice (n = 4 per genotype) were deeply anesthetized with isoflurane and decapitated. The cerebellum and underlying hindbrain were rapidly dissected, embedded in O.C.T., and rapidly frozen in a dry ice–ethanol bath. From each animal, a single 10-μm thick sagittal section of the lateral cerebellar vermis was collected onto a 3 x 3 mm tile containing spatially indexed beads (Curio Seeker 3x3 Seeker Kit Bundle V1.1; Curio Bioscience). RNA hybridization to beads, reverse transcription, tissue clearing and bead resuspension, second strand synthesis, and cDNA amplification were performed according to the manufacturer’s protocol (Curio Seeker 3x3 mapping kit manual v2; Curio Bioscience), based on Slide-seq v2 methodology^194^. Amplified cDNA was purified using the SPRIselect reagent (Beckman Coulter), eluted in nuclease- free water, and quantified using a D5000 ScreenTape Assay on a 4200 TapeStation System (Agilent). All samples had DNA concentrations > 0.2 ng/µL and showed no primer-dimer contamination.

Illumina-compatible libraries were prepared from cDNA using the Nextera XT DNA Sample Preparation kit (Illumina) and purified using SPRIselect reagent. Final library concentrations exceeded 1 ng/µL, as verified with a Bioanalyzer (Agilent). Libraries were sequenced on an Illumina NovaSeq 6000 using the NovaSeq 6000 S1 Reagent Kit v1.5 (300 cycles) in a paired-end 50 bp configuration with dual 8 bp index reads. Each flow cell contained eight multiplexed samples, yielding approximately 200 million reads per sample with a 2% PhiX spike-in for quality control.

#### Metabolomics

Six-month-old male WT and KO mice (n = 12 WT, 10 KO) were deeply anesthetized with isoflurane and decapitated. The cerebellum was rapidly dissected, flash-frozen in dry ice, and stored at -80°C. Analysis of TCA cycle and amino acid intermediates was conducted using gas chromatography-mass spectrometry (GC-MS) on an Agilent 5977B GC/MSD, as previously described^195^. Briefly, ∼60 mg of frozen cerebellar tissue was extracted using 1 ml of a water/methanol/acetonitrile mixture (1:2:2). Post-extraction, solvents were transferred to a GC vial and dried using nitrogen gas. The extracts were reconstituted with 20 μL of MTBSTFA +1% TBDMCS and 40 μL of acetonitrile, then heated for 45 minutes at 75°C to produce TBDMS derivatives. These derivatives were detected using Electronic Ionization, and compound intensities were estimated using internal standard calibration curves with tricarballylic acid as a surrogate.

#### Argonaute2-FLAG HITS-CLIP

HITS-CLIP of FLAG-tagged Ago2 was performed as previously described^105,196,197^, with minor modifications. *Rosa26^LSL-Flag^*^-*Ago2*^ mice expressing Cre-inducible FLAG-HA2-tagged Ago2 from the *Rosa26* locus^196^ were crossed with *Pcp2^Cre^* mice^159^ or *Pvalb^Cre^* mice^160^ to drive tagged Ago2 expression in Purkinje cells or Purkinje and inhibitory interneurons, respectively. These “WT” mice were further crossed to *miR-206^fl/fl^* mice to generate conditional knockouts (“cKO”) lacking miR-206 in the same cells expressing Ago2, and both WT and cKO mice were used for comparative HITS-CLIP.

Two-month-old male mice (n = 5 per genotype) were deeply anesthetized with isoflurane, and cerebella were rapidly dissected, flash-frozen in liquid nitrogen, and stored at -80°C. Samples were then thawed on ice, triturated, and crosslinked twice on ice at 400 mJ/cm^2^ using a UV Stratalinker 2400 (Stratagene). Crosslinked tissue was homogenized in high-salt PXL buffer, sonicated, and treated sequentially with RQ1 DNase (Promega) followed by RNase A (Thermo Fisher Scientific). FLAG-Ago2-RNA complexes were immunoprecipitated using anti-FLAG magnetic beads (Millipore Sigma) pre-blocked with BSA (Jackson ImmunoResearch), tRNA (Thermo Fisher Scientific), and salmon sperm DNA (Thermo Fisher Scientific). After a series of high- and low-salt washes, beads were treated with alkaline phosphatase (Millipore Sigma) and polynucleotide kinase (PNK, New England Biolabs). RNA-protein complexes were digested with proteinase K (Millipore Sigma) and RNA was extracted using Trizol LS (Thermo Fisher Scientific), followed by chloroform extraction and precipitation of RNA with isopropanol and GlycoBlue Coprecipitant (Thermo Fisher Scientific). Purifed RNA was washed with 80% ethanol and resuspended in nuclease-free water.

RNA integrity and concentration were assessed using a Qubit Flex Fluorometer (Thermo Fisher Scientific) and RNA 6000 Pico kit with a Bioanalyzer (Agilent). Small RNA libraries were prepared using the NEBNext Multiplex Small RNA Library Prep kit (New England Biolabs) according to the manufacturer’s instructions. Fragments below 200 bp, corresponding to containing adapter-linked RISC-associated miRNA and mRNA target regions, were purified by gel extraction. Libraries were quantified, pooled, and sequenced on an Illumina NextSeq 550 platform (75 cycles, single-end, 75 bp reads) to a target depth of ∼10 million reads per sample.

#### TRAP-seq

Purkinje cell–specific translating ribosome affinity purification (TRAP) was performed as previously described^113,114,198^, with minor modifications. *Rosa26^CAG-LSL-EGFP-L10a^*mice expressing Cre-inducible EGFP-tagged large ribosomal protein subunit L10a from the *Rosa26* locus^199^ were crossed with *Pcp2^Cre^* mice to drive transgene expression in Purkinje cells. These “WT” mice were further crossed to *miR-206^fl/fl^* mice to generate conditional knockout mice (“cKO”) lacking miR-206 in Purkinje cells; WT and cKO mice were used for comparative TRAP-seq.

Two-month-old male mice (n = 4 per genotype) were deeply anesthetized with isoflurane, and cerebella were rapidly dissected and homogenized in ice-cold homogenization buffer containing DTT, RNase and protease inhibitors, and freshly added cycloheximide (100 µg/mL). Lysates were centrifuged at 2,000 × *g* for 10 min. NP-40 alternative (final 1%) and DHPC (final ∼30 mM) were added to the supernatant, followed by incubation on ice for 5 min and clarification at 20,000 × *g* for 15 min at 4°C.

Streptavidin MyOne T1 Dynabeads (Thermo Fisher Scientific, 200 µl per sample) were pre-coupled to biotinylated Protein L (80 µL; Pierce) and incubated with mouse anti-EGFP monoclonal antibodies^114^ (33.3 µg each of clones 19C8 and 19F7, Heintz Lab at Rockefeller University). Cleared lysate was incubated with antibody-coupled beads overnight at 4°C with gentle rotation. Beads were washed three times in 0.15 M KCl wash buffer and four times in 0.35 M KCl high-salt wash buffer, both containing freshly added cycloheximide. Ribosome-bound RNA (IP fraction) was eluted in RLT buffer (Qiagen) containing β-mercaptoethanol. Input and post-IP supernatant (Unbound) fractions were retained for quality control.

RNA was purified using the RNeasy Plus Micro kit (Qiagen) according to the manufacturer’s instructions and eluted in 14 µL RNase-free water. RNA concentration and integrity were measured using High Sensitivity RNA ScreenTape analysis (Agilent). Samples with RNA Integrity Number (RIN) ≥ 6.5 were considered acceptable for downstream applications. One KO sample failed to pass quality control and was excluded from the study. Enrichment of *Gfp* and Purkinje cell marker *Calb1* and depletion of granule cell marker *Pax6* in bound/IP versus input and unbound fractions was confirmed by qPCR. 15 ng of total RNA per sample was used for cDNA amplification and library preparation with the Illumina stranded mRNA kit (Illumina). Libraries were quantified, pooled, and sequenced on an Illumina NextSeq 550 platform (300 cycles, paired-end, 150 bp reads) yielding ∼40–60 million reads per sample.

#### *Ex vivo* Purkinje cell electrophysiology

##### Acute slice preparation for cell-attached patch-clamp recordings

Two-month-old male WT and miR-206 knockout KO mice were deeply anesthetized with isoflurane and decapitated. Sagittal cerebellar vermis slices (300 µm thick) were prepared using a Compresstome vibratome (Precisionary Instruments). Slices were collected in ice-cold sucrose-based cutting solution containing (in mM): 215 sucrose, 2.5 KCl, 1.6 Na_2_HPO_4_, 26 NaHCO_3_, 4 MgSO_4_, 1 CaCl_2_, and 20 glucose. Slices were incubated in continuously oxygenated (95% O_2_/5% CO_2_) artificial cerebrospinal fluid (ACSF) containing (in mM): 120 NaCl, 3.3 KCl, 1.2 NaHPO_4_, 26 NaHCO_3_, 1 MgSO_4_, 2 CaCl_2_, and 11 glucose (pH 7.2; 300 mOsm) at 32°C for 30 min, then maintained at room temperature for at least one hour prior to recording.

##### Cell-attached patch-clamp recordings of spontaneous action potentials

Slices were transferred to a submersion recording chamber and perfused with ACSF (1 mL/min) at room temperature. Neurons in the cerebellum were visualized using an Olympus BX51WI microscope equipped with differential interference contrast optics and infrared illumination. Recordings were obtained from Purkinje neurons in lobule VI using glass pipettes (2.5–3.5 MΩ) filled with ACSF. The pipettes were placed against the membrane to form a giga-ohm seal without rupturing the membrane. Spontaneous firing activity was recorded in current clamp mode (I = 0) in gap-free configuration for 3 min per neuron at a sampling rate of 500 kHz using a Multiclamp 700B amplifier (Molecular Devices).

##### Acute slice preparation for whole-cell patch clamp recordings

Two-month-old male WT and miR-206 knockout (KO) mice were anesthetized with isoflurane and transcardially perfused with ice-cold, oxygenated (95% O_2_/5% CO_2_) N-methyl-D-glucamine (NMDG)–HEPES solution containing (in mM): 92 NMDG, 2.5 KCl, 1.2 NaH_2_PO_4_, 30 NaHCO_3_, 20 HEPES, 25 glucose, 5 sodium ascorbate, 2 thiourea, 3 sodium pyruvate, 10 MgSO_4_·7H_2_O, and 0.5 CaCl_2_·2H_2_O; pH adjusted to 7.3–7.4, 300–310 mOsm. Brains were rapidly removed and immersed in ice-cold NMDG–HEPES solution for 1 min. Sagittal cerebellar slices (200 µm) containing lobule VI of the vermis were cut using a vibratome (Leica VT1200S) or Compresstome (Precisionary Instruments VF-310-0Z). Slices were transferred to a pre-warmed (32°C) recovery chamber and subjected to a stepwise Na^+^ spike-in protocol^200^, then maintained at room temperature (22–24°C) for at least 1 h in oxygenated (95% O_2_/5% CO_2_) HEPES-based holding solution containing (in mM): 92 NaCl, 2.5 KCl, 1.2 NaH_2_PO_4_, 30 NaHCO_3_, 20 HEPES, 25 glucose, 5 Na^+^ ascorbate, 2 thiourea, 3 Na^+^ pyruvate, 2 MgSO_4_·7H_2_O, and 2 CaCl_2_·2H_2_O.

##### Voltage- and current-clamp recordings

Patch electrodes (3–5 MΩ) were pulled from borosilicate glass capillaries (World Precision Instruments) using a micropipette puller (PC-10; Narishige). Slices were transferred to a recording chamber perfused with standard artificial cerebrospinal fluid (ACSF) containing (in mM): 124 NaCl, 2.5 KCl, 1.2 NaH_2_PO_4_, 24 NaHCO_3_, 5 HEPES, 12.5 glucose, 2 MgSO_4_·7H_2_O, and 2 CaCl_2_·2H_2_O; pH 7.3–7.4, 295–305 mOsm. Slices were continuously superfused with oxygenated ACSF (95% O_2_/5% CO_2_) at 32°C.

Whole-cell patch-clamp recordings were performed under a SliceScope Pro 2000 upright microscope (Scientifica) equipped with infrared differential interference contrast optics. Signals were amplified using a Multiclamp 700B amplifier (Molecular Devices), low-pass filtered at 3 kHz, amplified fivefold, and digitized at 10 kHz using a Digidata 1550 A/D converter (Molecular Devices). Input resistance was continuously monitored, and recordings were discarded if changed by >20%.

##### Whole-cell current-clamp recordings of spontaneous action potentials

For action potential measurements in Purkinje neurons, patch pipettes were filled with a potassium-based internal solution containing (in mM): 130 K^+^ gluconate, 4 KCl, 0.3 EGTA, 10 HEPES, 4 MgATP, 0.3 Na_2_GTP, and 10 phosphocreatine; pH 7.3 (KOH). The external solution was standard ACSF. After establishing whole-cell configuration, resting membrane potential was noted and action potential firing was recorded immediately in current-clamp mode, as Purkinje neurons displayed rundown during prolonged recordings.

##### Whole-cell voltage-clamp recordings of potassium and HCN channel-mediated currents

To measure potassium channel and HCN channel currents in Purkinje cells, the same potassium-based internal solution was used. The external ACSF contained picrotoxin (100 µM), kynurenic acid (1 mM), tetrodotoxin (1 µM), and CdCl_2_ (200 µM) to block inhibitory and excitatory synaptic inputs, and Na^+^ and Ca^2+^ channels. Membrane capacitance was measured following membrane rupture and neurons were held at –70 mV. To measure potassium channel currents, a voltage step function beginning with a 150 ms hyperpolarization at –110 mV to deactivate all opened or inactivated channels, followed by 4-sec depolarization steps from –70 to +80 mV in 10 mV increments, and resulting currents were recorded. To measure HCN channel currents, 700-ms hyperpolarizing voltage steps from –70 to –130 mV in 10 mV increments were applied and resulting currents recorded. Purkinje cells from a separate cohort of WT mice were recorded while applying the same hyperpolarizing voltage step function before and after wash-in of ACSF or 30 µM of HCN channel blocker ZD7288.

##### Extracellular electrophysiology recordings

Acute parasagittal cerebellar slices were prepared as described for whole-cell recordings. Extracellular recordings were obtained from Purkinje neurons in lobule VI, using 7-10 ΜΩ pipettes containing ACSF. Purkinje cell firing was recorded under current clamp mode (gap-free) at I=0 and a sampling rate of 2 µs for at least 5 minutes using a Multiclamp 700B amplifier (Axon Instruments).

### QUANTIFICATION AND STATISTICAL ANALYSIS

Data are presented as mean ± SEM unless otherwise indicated. Statistical analyses were performed in GraphPad Prism (v 10.2.0) and R (v4.3.0). Two-group comparisons were made using unpaired two-tailed Welch’s t-tests, and multiple comparisons by one-way or two-way ANOVA followed by Šidák’s or Tukey’s post hoc test. All Purkinje cell morphological and electrophysiological analyses were performed blind to genotype. All statistical tests, sample numbers, and p values or adjusted p values are found in Supplemental Table S12. p < 0.05 was considered significant.

#### Purkinje cell and PV neuron cell counts

For quantification of WT and miR-206 KO Purkinje cell and PV neuron numbers, 10x tiled epifluorescence images of cerebellar sagittal sections were used. CALB1- or PV-immunoreactive cells were manually counted using the counter tool in Photoshop (Adobe), and counts were averaged across six sections (PC counts, cerebellum) or summed across multiple sections (PV neuron counts from 4, 16, and 10 sections for mPFC, dorsal hippocampus, and GPe, respectively) per mouse. For quantification of Purkinje cells in WT and KO mice expressing viral *Pcp2*-GFP or *Pcp2-*miR-206-GFP, CALB1-immunoreactive cells were segmented and counted using CellPose 2.0^201,202^, a deep-learning-based segmentation algorithm, and GFP-expressing Purkinje cells were manually counted using Photoshop (Adobe). Cell counts were summed across six sections per mouse.

#### *Calb1* and *Aldoc* RNAscope fluorescence intensity

Three 10x tiled epifluorescence images of sagittal sections of cerebellar vermis were analyzed per mouse. Purkinje cells were segmented based on *Calb1* fluorescence using CellPose 2.0, and cell outlines were imported into ImageJ as regions of interest (ROIs) along with raw image data. Mean fluorescence intensity for *Calb1* and *Aldoc* channels was calculated per cell, exported, and analyzed in Graphpad Prism. Data from at least 2500 Purkinje cells were evaluated per mouse.

#### CALB1 and DAPI immunofluorescence intensity

For quantification of fluorescence intensity in Purkinje cell soma and the molecular layer containing their dendrites, 40x confocal z-stack images of lobule VI were used. For soma measurements, a single optical section corresponding to the middle of each soma along the z-axis was manually selected in Zen Blue software (v 4.0.3). Soma boundaries were traced manually using the spline contour tool, and soma area and mean fluorescence intensity were measured for each traced region. At least 36 Purkinje cells were analyzed from lobule VI per animal.

#### Purkinje cell morphology and branching

Purkinje cells with Lucifer Yellow-filled soma and primary dendrites were selected for analyses of soma area and number of primary dendrites using Zen Blue software. Each soma boundary was traced for a single optical section at the middle of the soma along the z-axis, and area measured. The number of primary dendrites for each Purkinje cell was manually counted from orthogonal projections of z-stack images. For Purkinje cells with soma and dendritic trees completely filled with Lucifer Yellow, a straight line was drawn from the midpoint of the soma to the distal dendritic tree endings within the z-stack orthogonal projection and distance measured using Zen Blue software.

Fully labeled Purkinje cells with clearly distinguishable dendritic trees were reconstructed in Neurolucida360 (MBF Bioscience). Neurolucida Explorer was used for Sholl analysis^65^ to quantify the number of intersections, dendritic length, and number of branch nodes at 10-µm increments from the soma, as well as to calculate totals for each dendritic measure. Convex hull surface area and volume of the dendritic field were also computed^203^.

Sholl data were analyzed using a linear mixed-effects model to account for hierarchical sampling of multiple neurons per animal. Each dependent measure (intersections, dendritic length, or branch node number) was modeled as a function of genotype, radius, and their interaction, with random intercepts for neurons nested within animals. To model spatial dependence across radii, an autoregressive correlation structure (AR(1)) was incorporated^204^. Models were fit in R using the *lmer* function in the *lme4* package^205^. Type III Wald F-tests assessed main and interaction effects, and p-values were obtained via ANOVA. Post hoc pairwise comparisons were performed using the *emmeans* package^206^ to generate estimated marginal means with Wald t-tests. Convex hull area, soma surface area, and total branching measures were compared using Welch’s t-tests in GraphPad Prism.

#### Prepulse inhibition of acoustic startle

The average voltage of the peak startle amplitude (V_max_) and average latency to peak startle amplitude (T_max_) were calculated for the first six startle trials. Percentage habituation of the acoustic startle response was calculated as 100 × [(V_max_ average of first six startle trials) – (V_max_ average of last six startle trials)] / (V_max_ average of first six startle trials). Percentage pre-pulse inhibition of the acoustic startle response was calculated after subtracting the average amplitude of the no-stimulus trials from the average peak amplitude of the middle block of pulse-alone trials and 74-, 78-, or 86-dB pre-pulse trials. %PPI = 100 × [(V_max_ average of pulse-alone trials) – (V_max_ average of pre-pulse trials)] / (V_max_ average of pulse-alone trials).

PPI data were analyzed using two-way repeated measures ANOVA with Volume as the within-subject factor and Genotype as the between-subject factor. For WT/KO mice treated with control or miR-206-expressing virus, PPI data were analyzed using three-way repeated-measures ANOVA with Volume as the within-subject factor and Genotype (WT, KO) and Virus (control, miR-206) as between-subject factors. Type III sums of squares were used to account for unequal sample sizes, and Greenhouse-Geisser corrections were applied where sphericity assumptions were violated. Post-hoc comparisons were performed using Tukey-adjusted pairwise contrasts for significant main effects and interactions. P-values were adjusted to control for multiple comparisons.

#### Gait analyses

Digital video recordings of mice tested in the NeuroCube were processed using computer segmentation algorithms to extract locomotor behaviors, including gait, paw placement, limb coordination, and body movement (Psychogenics). NeuroCube features included geometric measures such as stride length, step length and base width; dynamic measures such as stride, stand, and swing duration; and paw features such as paw area, intensity, perimeter, and minimal and maximal diameters. Body motion metrics captured overall movement and variability of body dimensions, while coordination features described the relative positioning of paws, including angles defined by three-paw configurations. The most informative features defining the phenotype were identified and ranked using proprietary bioinformatics algorithms (Psychogenics), generating an overall discrimination index for all features or selected subsets. In the decorrelated feature space, Gaussian distributions representing each group were used to quantify overlap, providing a measure of separability (“distinguishability”) between WT and KO mice. Graphical representations were generated for each group, and p-values were calculated to assess the statistical significance of the discrimination ratios.

#### Single nuclei RNA sequencing

snRNA-seq reads were demultiplexed and aligned to the mouse mm10 genome, and gene counts quantified as UMIs using the *Cell Ranger* Software Suite (10x Genomics). Unfiltered Cell Ranger feature-barcode matrices were processed using *CellBender v0.3.2* for background noise correction and nuclei calling^207^. *CellBender* input parameters --expected-cells and --total-droplets-included were chosen for each file according to the estimated cell count and empty droplet plateau determined from the *Cell Ranger* barcode rank plot, according to *CellBender* usage guidelines. Feature-barcode matrices with (*CellBender*-filtered) and without (*Cell Ranger*-filtered) background noise correction were loaded into R as separate assays in a merged Seurat object and the extent of correction quantified as the difference in feature counts using the *scCustomize v3* and *Seurat v5* packages^208,209^. *CellBender*-filtered nuclei with at least 200 genes, less than 2% mitochondrial genes, and less than 1% ribosomal genes; and genes expressed in at least 10 nuclei were deemed to be high-quality and retained for further processing.

*SCTransform v2* (SCT) was applied to the filtered gene expression matrix for each animal in order to normalize counts, stabilize variance, and find variable features^210,211^. Principal component analysis with 30 dimensions was run on SCT data, and the resulting datasets integrated using reciprocal PCA (RPCA), with further batch correction with *Harmony*^212^. The *FindNeighbors* function in *Seurat* was used to construct a shared nearest-neighbor (SNN) graph based on the first 30 principal components. Cells were then clustered using the *FindClusters* function in *Seurat* employing the *Leiden* algorithm^213^. Cluster annotation was performed by transferring labels from a cerebellar cortex snRNA-seq reference atlas^66^ using the *TransferData* function in *Seurat*. Marker genes with at least 60% differential expression were identified for each cluster using the *FindAllMarkers* function on SCTransform-normalized data. Feature plots and dot plots of top marker genes identified in this study and previous studies^66^ were generated in *Seurat*. Differential gene expression (DGE) analysis was conducted using the *run_de* function from the *Libra* package^214^, applying the pseudobulk edgeR-LRT workflow to the raw count data. DGE data for each cell type were filtered for genes expressed in at least 10% of WT or KO cells (Tables S1 and S2) and plotted using the EnhancedVolcano package^215^.

#### Curio Seeker / Slide-seq v2

FASTQ files were processed by Curio Bioscience using the Curio Seeker bioinformatics pipeline for demultiplexing and barcode-based read assignment. Resulting Seurat objects containing gene expression matrices and spatial metadata were returned for downstream analyses. Using *scCustomize v3* and *Seurat v5* in R, data were filtered as follows: genes expressed in >= 5 cells (corresponding to beads), and cells with > 100 features, % mitochondrial genes <=5, and % ribosomal genes <=3. SCTransform, PCA, integration, batch correction, and unsupervised clustering were performed with the same parameters as those described for snRNAseq above, and clusters visualized using *spatialDimPlot* function. A distinct cluster corresponding to beads capturing background RNA in regions lacking tissue was identified and excluded from downstream analyses. *Seurat* objects were then subsetted to cerebellar cortex (Cb Cx) and deep cerebellar nuclei (CN) using spatial coordinates generated by tracing spatial DimPlot images in Fiji^216^.

First- and second-predicted cell type annotations were generated by robust cell type decomposition (RCTD) in the *spacexr* package^70^ using reference snRNA-seq datasets for Cb Cx^66^ and CN^69^. For the CN reference, cell types were grouped based on the original study as follows: “ex_0–ex_7/Class-A” = “CN_Excitatory_A”; “ex_8–ex_15/Class-B” = “CN_Excitatory B”; “gly_0–gly_4” = “CN_Glycinergic”; and “inh_1” = “CN_Inhibitory”. Spatial arrangements of the predicted cell identities were visualized using the *SpatialDimPlot* function in *Seurat,* and relative proportions calculated in R and plotted using *ggplot2*^217^. Dot plots of select marker genes from the reference datasets were generated for each predicted cell type using the *DotPlot_scCustom* function in *scCustomize*. Differential gene expression between WT and KO was assessed separately for cerebellar cortex (Cb Cx) and CN using cell type-specific inference of differential expression (C-SIDE) in the *spacexr* package^71^ (Tables S3 and S4, respectively) and plotted for each cell type or for all CN neurons using *EnhancedVolcano*. Gene enrichment in molecular ontology pathways was performed using *ClusterProfiler*^218,219^ and visualized using heat plots and gene-concept network (*cnet*) plots. Enrichment of disease-gene associations was analyzed using EnrichR^220–222^, querying GeDipNET^223^ and DisGeNET^224^ disease-gene association libraries. Enrichment results were plotted as dot plots in R using the *ggplot2* package, with dot color and size representing significance and gene count, respectively.

#### Argonaute2-FLAG HITS-CLIP

Sequencing data were assessed for quality using *FASTQC*^225^, and adapters were trimmed using *Trimmomatic*^226^. Poly(A) sequences were also removed by using a modified adapter file in *Trimmomatic*. Trimmed reads were aligned to the *Mus musculus* (mm10) genome using *HISAT2*^227^. Deletion of miR-206 in cKO samples was confirmed by visual inspection of sequencing reads using *IGV* Integrative Genomics Viewer^228^. BigWig alignment files were generated from mapped BAM files using *deepTools*^229^ with counts-per-million (CPM) normalization. HITS-CLIP coverage over the miR-206 locus (chr1:20,679,045–20,679,087; mm10) was extracted from normalized bigWig files using *rtracklayer*^230^ and *GenomicRanges*^231^ in R and visualized with *Gviz*^232^. Coverage tracks were grouped by genotype (WT, cKO), and the mean ± SEM plotted for the miR-206 locus.

Read coverage across transcript features was quantified from bigWig alignment files in R. Genomic annotations for *Mus musculus* (mm10 assembly) were obtained from the *TxDb.Mmusculus.UCSC.mm10.knownGene* package. Genomic ranges corresponding to exons, introns, 5′ untranslated regions (5′UTRs), and 3′ untranslated regions (3′UTRs) were extracted using the *GenomicFeatures*^231^ and *rtracklayer*^230^ packages. For each sample, normalized signal intensity values were imported from BigWig files using the import() function (*rtracklayer*). Coverage overlapping each genomic feature type was calculated by summing signal scores within the corresponding feature intervals. Percentages of total coverage mapping to exons, introns, 5′UTRs, and 3′UTRs were then computed per sample.

A gene count matrix was generated from mapped HITS-CLIP sequences using *htseq-count* within *HTSeq*^233,234^, imported into R, and annotated with gene biotype information retrieved from Ensembl using *biomaRt*^235^. Read counts were summed per biotype for each sample and converted to percentages of total mapped reads. Overall biotype proportions were displayed as stacked bar graphs in GraphPad Prism. Differential gene expression (DE) between WT and cKO HITS-CLIP samples was assessed using the *edgeR*^236^ package’s likelihood ratio test (LRT), applying *filterByExpr* for minimal expression filtering prior to normalization and model fitting (Tables S5 and S6). DE results from *edgeR* were annotated with gene biotype and Ensembl identifiers using *biomaRt*. Genes were classified as miRNA, lncRNA, protein-coding, or other, and enrichment of significantly altered genes (*P* < 0.05) within each RNA class was assessed using Fisher’s exact test with Benjamini–Hochberg correction; results were visualized as a bubble plot using *ggplot2* in R.

Aligned HITS-CLIP SAM files were concatenated based on genotype and analyzed using *dCLIP*^111^ to identify differential binding between WT and cKO samples. miRNA targets were inferred from clusters showing higher binding in WT compared to cKO samples. Because miRNA binding is predominantly in the 3’UTR of targets, the analysis focused on clusters within 3’ UTRs. The top 500 transcripts with predicted differential binding, ranked by average binding strength, were retained for downstream analysis. Average binding strength and binding differential of the top transcripts for each experiment (Tables S7 and S8) were visualized as x-y scatterplots using the *ggplot2*. The overlap between the top *dCLIP*-predicted transcripts from the *Pvalb^Cre^*and *Pcp2^Cre^* experiments (“shared transcripts”, Table S9) was analyzed using Fisher’s Exact Test with Benjamini–Hochberg correction in GraphPad Prism. DE tables were cross referenced with transcripts predicted by TargetScan 8.0 to be targets of miR-1-3p/miR-206-3p^112,237^ (Table S10); TargetScan-predicted targets are in bold font in Tables S7–S9.

Shared *dCLIP*-predicted transcripts were analyzed for functional enrichment using the EnrichR platform^220–222^ and *enrichR* package in R, querying the following libraries: GO Molecular Function (GO_MF), GO Biological Process (GO_BP), GO Cellular Component (GO_CC), GeDiPNet, and DisGeNET. Enrichment results were plotted as dot plots in R using the *ggplot2* package, with dot color and size representing significance and gene count, respectively.

#### TRAP-seq

Sequencing data were assessed for quality using *FASTQC*^225^, and adapters were trimmed using *Trimmomatic*^226^. Trimmed reads were aligned to the *Mus musculus* (mm10) genome using *HISAT2*^227^. A gene count matrix was generated from the mapped TRAP-seq sequences using *htseq-count* within *HTSeq*^233^ and imported into R for subsequent analyses. Genes with consistently low counts across samples were removed using the *filterByExpr* function in *edgeR*. Remaining counts were normalized for library size and composition using the trimmed mean of M-values (TMM) method. Multidimensional scaling of TMM-normalized counts revealed no sample outliers. Normalized counts were converted to counts per million (CPM) and log2-transformed for downstream visualization. Mean normalized counts for Purkinje cell, astrocyte, granule cell, and ODC markers were plotted for WT and cKO groups in GraphPad Prism.

Differential expression between WT and cKO samples was assessed using the edgeR quasi-likelihood F-test (QLF), which models counts with a negative binomial generalized linear model and incorporates gene-specific dispersion estimates (Table S11). MA plots displaying log2 fold change vs average log2 CPM were generated using *ggplot2*. Mean normalized counts for the top DE transcripts by *P* value were plotted for WT and cKO groups in GraphPad Prism. DE transcripts (*P* value <0.05) were analyzed for functional enrichment using the EnrichR platform^220–222^ and *enrichR* package in R, querying the following libraries: GO Molecular Function (GO_MF), GO Biological Process (GO_BP), GO Cellular Component (GO_CC), GeDiPNet, and DisGeNET. Enrichment results were plotted as dot plots in R using the *ggplot2* package, with dot color and size representing significance and gene count, respectively. Log2 CPM expression data of significantly altered Kcn-family genes (*P* < 0.05) were z-score normalized and visualized as a heatmap using the *pheatmap* package in R.

#### Cell-attached electrophysiology of Purkinje cells

For each cell, spike number was quantified over a one-minute interval in the middle of the recording using template-based event detection in pClamp 10.3 (Molecular Devices). Templates were generated from the average of 20 events, and automated detections were verified manually. Mean firing rates were calculated per cell.

#### Voltage- and current-clamp electrophysiology of Purkinje cells

##### Patch clamp recordings of Purkinje cell spontaneous action potentials

Purkinje neurons exhibited either silent or tonic firing patterns during recordings. Data from the first minute after achieving whole cell configuration were excluded from analysis. Action potential frequency and amplitude were quantified during the first tonic firing phase using threshold-based event detection in pClamp 10.3.

##### Potassium and HCN channel-mediated currents

Currents were normalized to membrane capacitance and expressed as current density (pA/pF).

##### Extracellular electrophysiology of Purkinje cells

Spike number was quantified for a one-minute period in the middle of the recording when the signal was most stable. For both tonic and burst-firing cells, spike number was measured using threshold-based event detection performed using Clampfit 10.7 (Molecular Devices) and the mean firing rate per cell was calculated. For burst-firing cells, calculations were performed for the entire train of burst activity.

## SUPPLEMENTAL FIGURE LEGENDS

**Figure S1.**
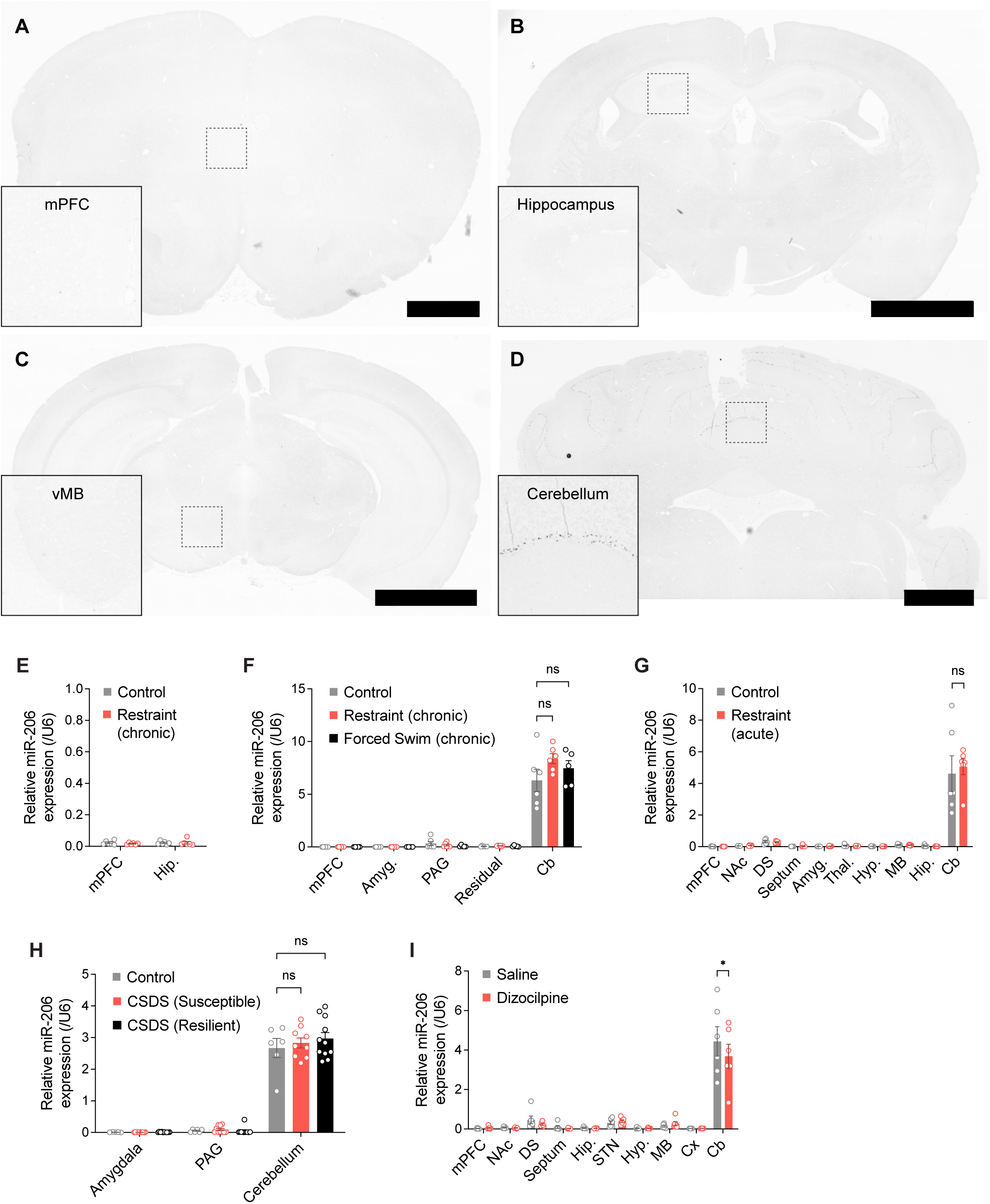
miR-206 expression in brain is undetectable outside of the cerebellum at baseline, after stress, or with NMDA receptor block. **(A–D)** *pre-miR-206* RNAscope signal is not found in coronal sections through medial prefrontal cortex (A), hippocampus (B), or ventral midbrain (C), but is only detected in the PC layer of cerebellum of adult male mice (D). Images were stitched from multiple tiled epifluorescence images. Scale bars: 1 mm. **(E–I)** Quantitative RT-PCR for mature miR-206 across brain regions in mice exposed to chronic restraint stress (E, F), forced swim stress (F), acute restraint stress (G), chronic social defeat stress (CSDS, susceptible and resilient groups), and subchronic administration of the non-competitive NMDA receptor antagonist dizocilpine (I). miR-206 was only detected in cerebellum across all treatment groups, and dizocilpine caused a slight reduction in expression in cerebellum. n = 6 control, 6 chronic restraint stressed male mice (E), 6 control, 6 chronic restraint stressed, 5 forced swim stressed male mice (F), 6 control, 6 acute restraint stressed mice (G), 6 control, 10 CSDS susceptible, and 11 CSDS resilient male mice (H), and 6 saline, 6 dizocilpine-treated male mice (I). *p = 0.0207, two-way ANOVA and Tukey’s multiple comparisons test. mPFC, medial prefrontal cortex; NAc, nucleus accumbens; DS, dorsal striatum; Septum; Amyg., amygdala; Thal., thalamus; Hyp., hypothalamus; MB, ventral midbrain; Hip, hippocampus; PAG, periaqueductal grey; Residual, remaining brain tissue after other regions dissected; STN, subthalamic nucleus; Cx, cortex; Cb, cerebellum. Error bars indicate SEM.

**Figure S2.**
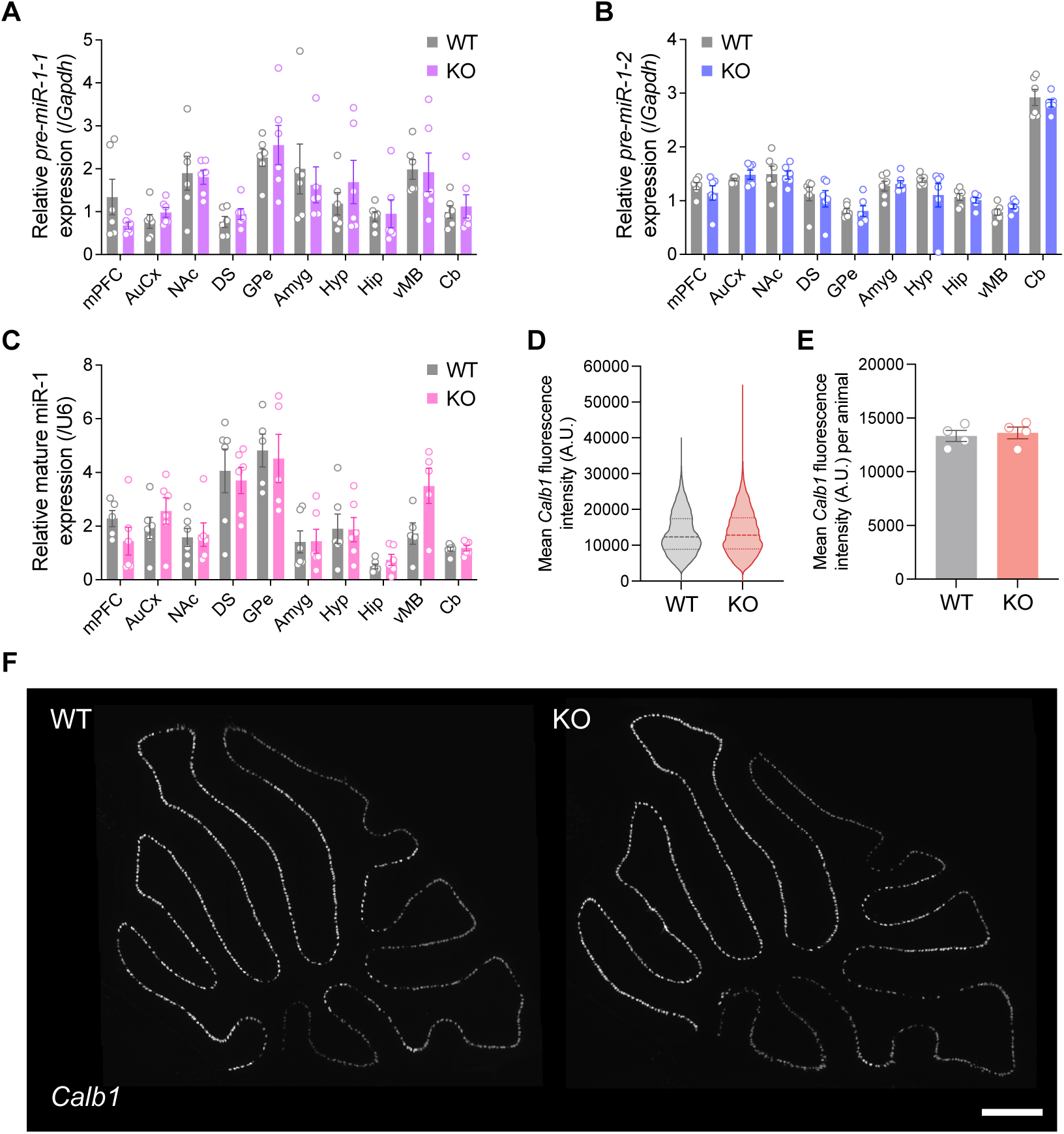
miR-1 and *Calb1* expression are unaffected in miR-206 KO mice. **(A–C)** Quantitative RT-PCR detection of (A) *pre-miR-1-1*, (B) *pre-miR-1-2*, and (C) mature miR-1 in brain regions of WT and miR-206 KO mice. mPFC, medial prefrontal cortex; AuCx, auditory cortex; NAc, nucleus accumbens; DS, dorsal striatum; GPe, globus pallidus external; Amyg, amygdala; Hyp, hypothalamus; Hip, hippocampus; vMB, ventral midbrain; Cb, cerebellum. n = 6 WT, 6 KO males. **(D–F)** *Calb1* RNAscope fluorescence intensity per PC (D) and averaged across all PCs per animal (E) are equivalent in WT and KO mice. n = greater than 2000 PCs per animal and 4 WT, 4 KO male mice. Representative tiled 10x epifluorescence images of sagittal cerebellar sections in (F). Scale bar: 500 µm.

**Figure S3.**
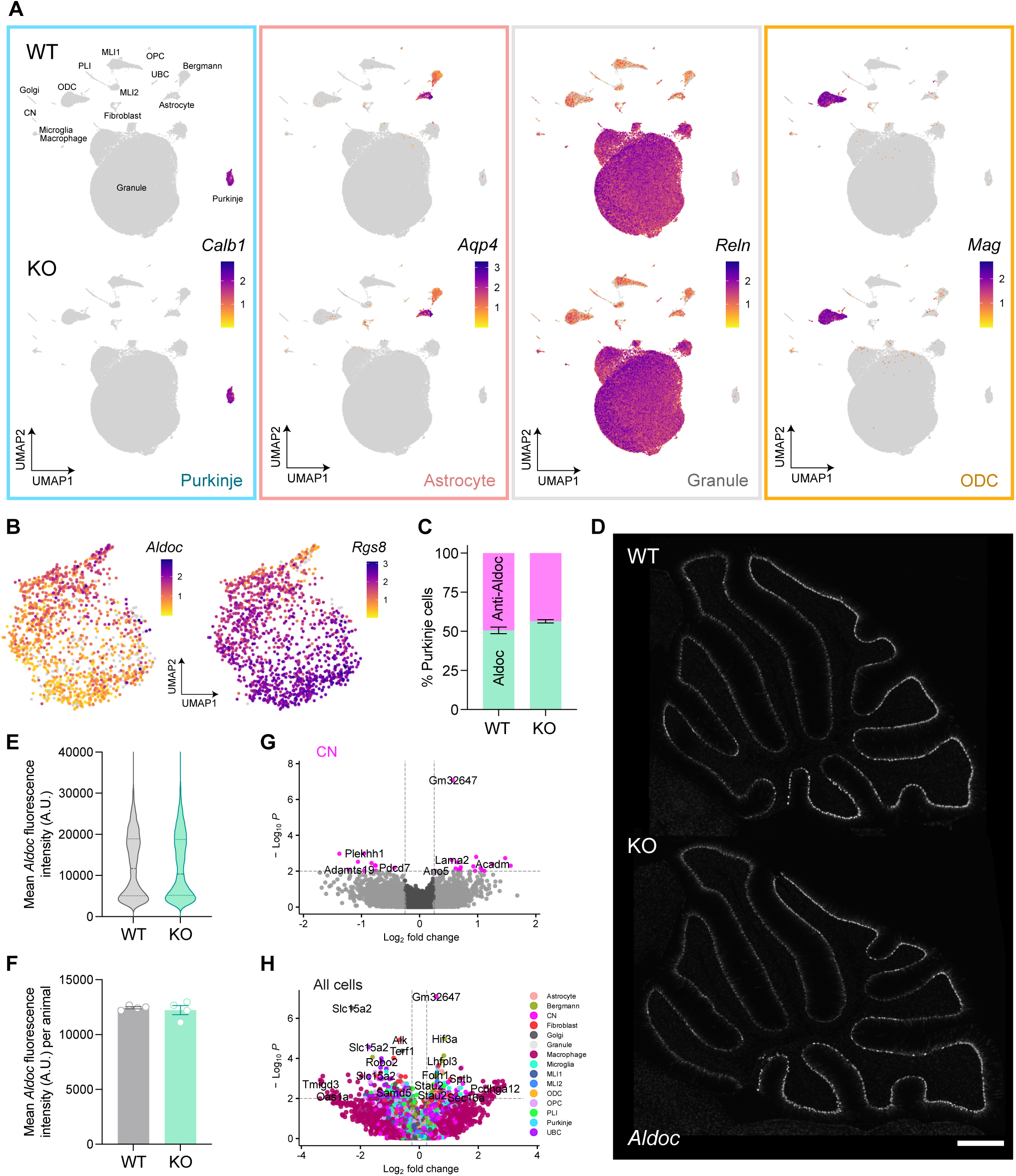
Extended snRNA-seq and RNAscope data, related to Figure 3. **(A)** Feature plots of scaled expression of known cerebellar marker genes confirm Purkinje, astrocyte, granule, and oligodendrocyte (ODC) cluster identities. **(B)** Feature plots of scaled expression of markers of Aldoc (*Aldoc*) and Anti-Aldoc (*Rgs8*) PC subtypes. **(C)** Percentage of PCs assigned Aldoc and Anti-Aldoc identies in WT and miR-206 KO mice. n = 4 WT, 4 KO male mice. **(D–F)** RNAscope detection of *Aldoc* expression in WT and KO cerebellum. Representative tiled 10x epifluorescence images of sagittal cerebellar sections in (D). Fluorescence intensity per PC (E) and averaged across all PCs per animal (F) are equivalent in WT and KO mice. n = greater than 2000 PCs per animal and 4 WT, 4 KO male mice. Scale bar: 500 µm. **(G–H)** Volcano plots of differential gene expression in KO cerebellar nuclei (CN) (G), and all cells (H). Error bars indicate SEM.

**Figure S4.**
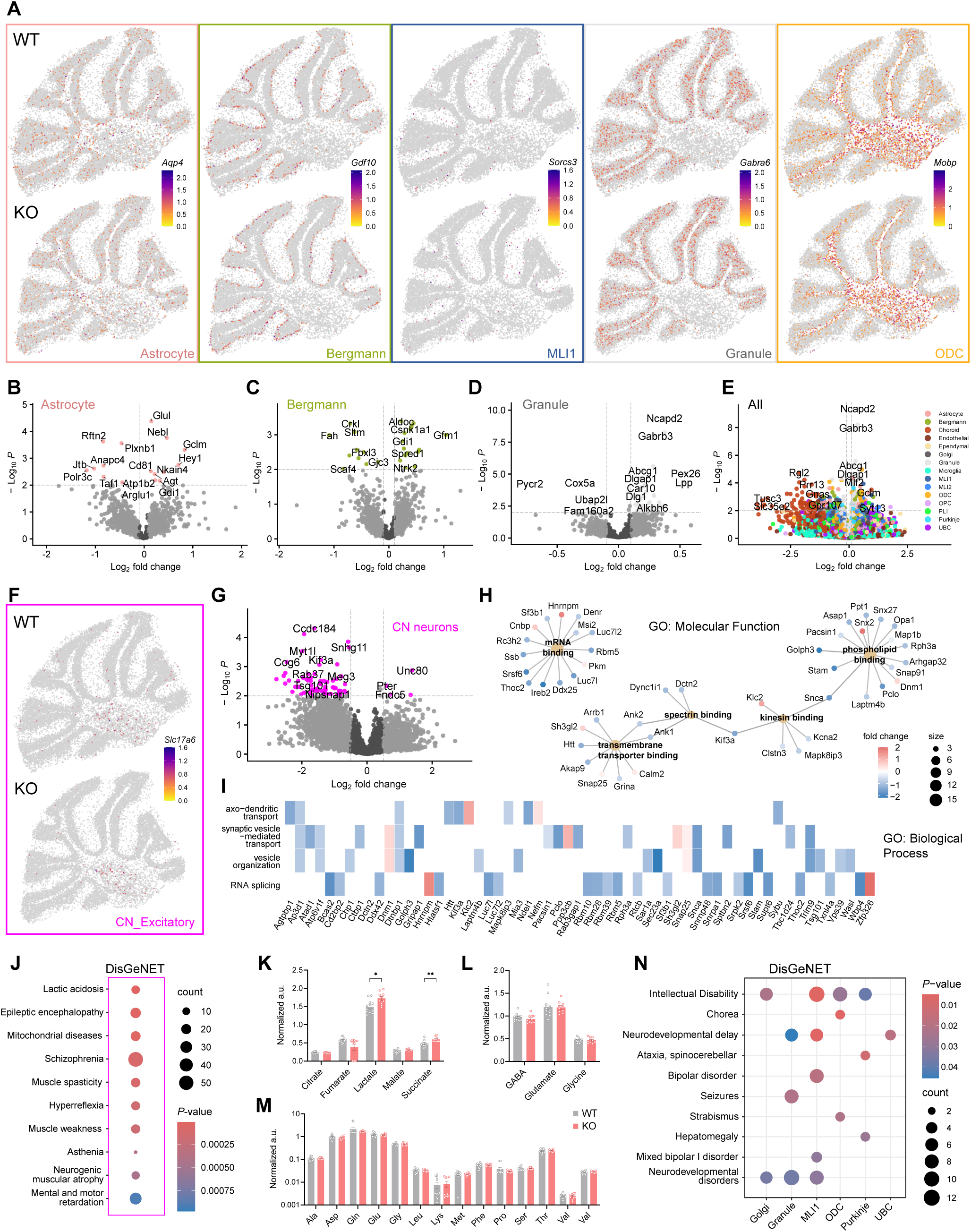
Extended spatial-seq data and metabolomics, related to Figure 3. **(A)** Spatial feature plots of scaled transcript expression of astrocyte, Bergmann glia, molecular layer interneuron 1 (MLI), granule and oligodendrocyte (ODC) markers in representative WT (top) and miR-206 KO (bottom) cerebellar sections. **(B–E)** Volcano plots of differential gene expression in KO astrocytes (B), Bergmann glia (C), granule cells (D), and all cell types (E). **(F)** Spatial feature plot of scaled expression of *Slc17a6*, an excitatory cerebellar nuclei (CN) neuron marker, in WT and KO samples. **(G)** Volcano plot of differential gene expression in KO CN neurons (all subtypes) using C-SIDE analysis. **(H)** Gene-concept network plot showing fold change of genes differentially expressed in KO CN neurons connected with their respective enriched molecular function gene ontology pathways. **(I)** Heat plot showing fold change of genes differentially expressed in KO CN neurons mapped to enriched biological process gene ontology pathways. **(J)** Dot plot of Disease Gene Network (DisGeNET) terms enriched in genes that are differentially expressed in KO CN neurons. **(K)** Mass spectrometry of tricarboxylic acid (TCA) cycle intermediates shows elevation of lactate and succinate in KO cerebellum. (L–N): n = 12 WT, 10 KO male mice. *p = 0.0312 (lactate), **p = 0.0079 (succinate), multiple t tests + Bonferroni correction. **(L–M)** Mass spectrometry shows no change in neurotransmitter (M) or proteinogenic (N) amino acid levels in KO cerebellum. **(N)** Dot plot of DisGeNET terms enriched in differentially expressed genes for multiple cerebellar cell types. Error bars indicate SEM.

**Figure S5.**
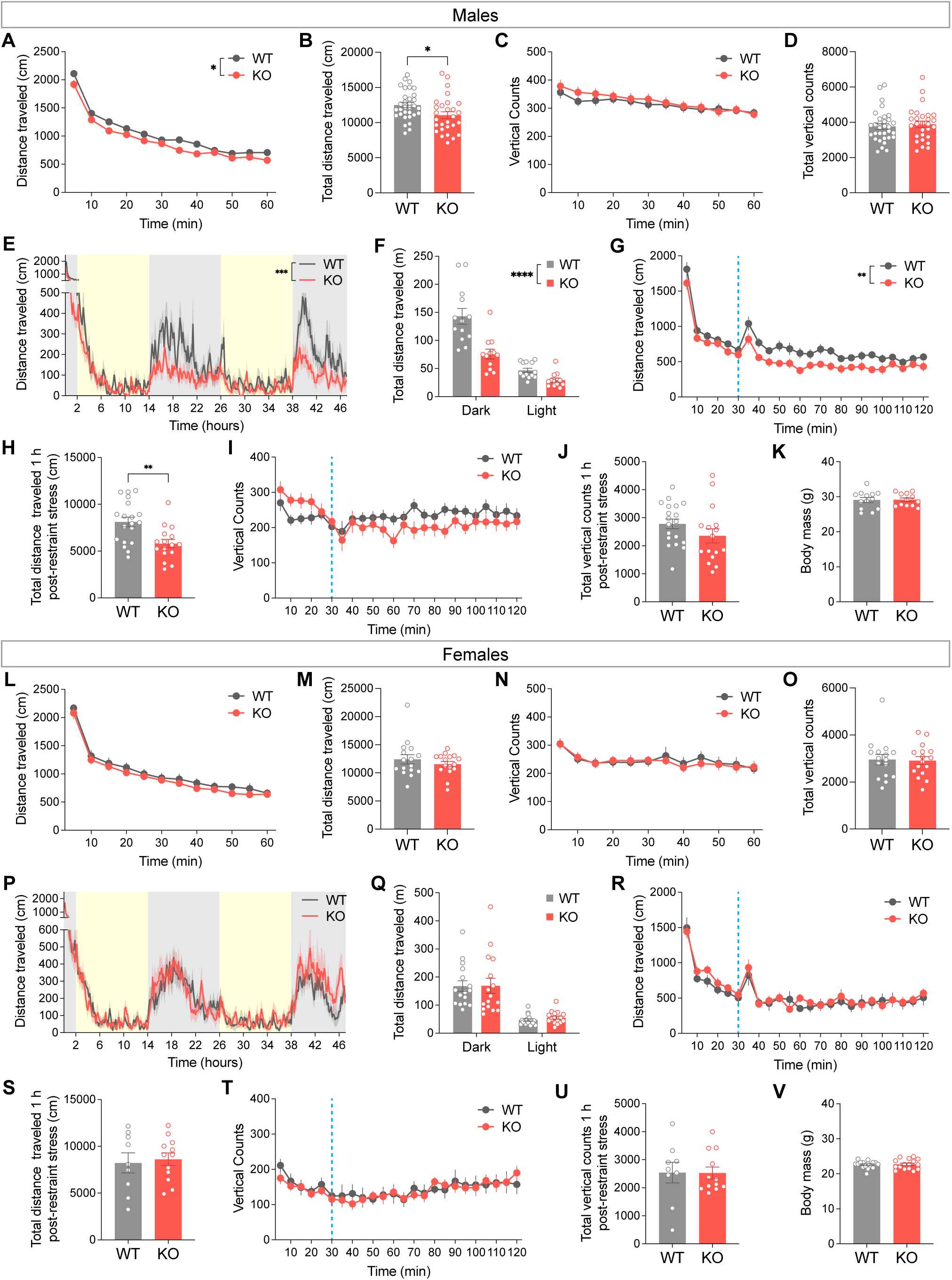
Male but not female miR-206 KO mice display stress-dependent hypolocomotion. **(A–D)** Male KO mice travel less distance in the open field chamber over 60 min (A–B) but vertical activity is unaffected (C–D). n = 32 WT, 28 KO mice. *p = 0.0192 (main effect of genotype), two-way RM ANOVA (A). *p = 0.0213, Welch’s t-test (B). **(E–F)** Male KO mice travel less distance in a home cage across two night-day cycles. Grey: dark/active phase; yellow: light/inactive phase. n = 13 WT, 12 KO mice. ***p = 0.0002 (main effect of genotype), two-way RM ANOVA (E). ****p < 0.0001, two-way ANOVA (F). **(G–J)** Male KO mice travel less distance as compared to WT in the open field chamber following two seconds of acute restraint stress (G–H), but vertical activity is similar for WT and KO (I–J). n = 19 WT, 16 KO mice. ** p = 0.0030, two-way RM ANOVA (G). ** p = 0.0018, Welch’s t-test (H). Data were analyzed for 60 min from time of restraint stress. **(K)** Body mass is equivalent in WT and KO males. n = 13 WT, 12 KO mice. **(L–O)** Female KO mice have no differences in baseline locomotion (L–M) or vertical activity (N–O) in the open field. n = 16 WT, 16 KO mice. **(P–Q)** Distance traveled in a home cage across two night-day cycles is similar for WT and KO females. n = 16 WT, 16 KO mice. **(R–U)** Distance traveled (R–S) and vertical activity (T–U) after acute restraint stress are equivalent in WT and KO females. n = 9 WT, 12 KO mice. **(V)** Body mass of females is not altered by miR-206 deletion. n = 16 WT, 16 KO mice. Error bars indicate SEM.

**Figure S6.**
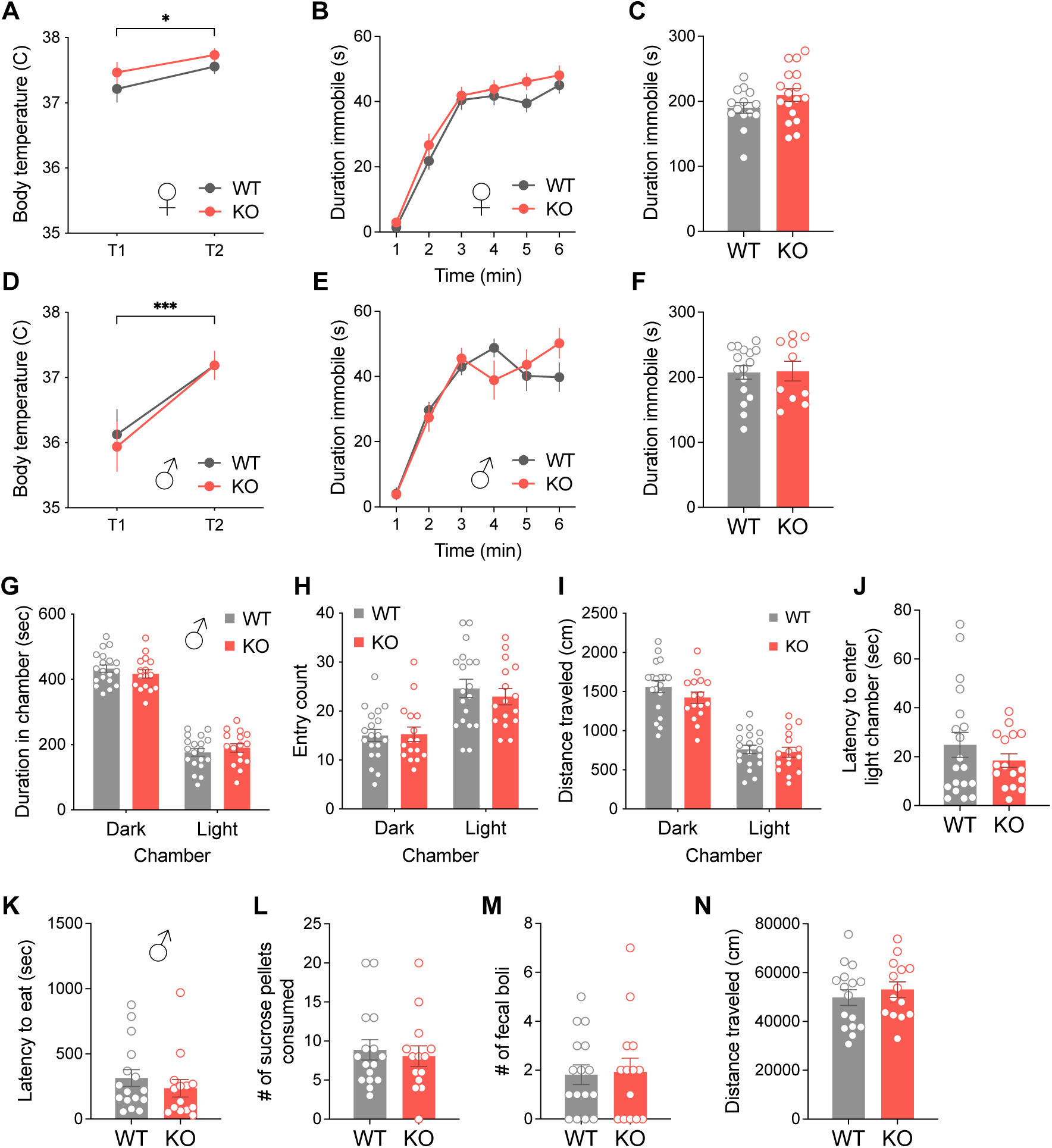
miR-206 deletion does not alter other stress- or anxiety-related behaviors. **(A,D)** Stress-induced hyperthermia is unchanged in miR-206 KO males (A) and females (D). n = 8 WT, 8 KO males; 7 WT, 9 KO females. ***p = 0.0002 (males), *p = 0.0111 (females) (main effect of time post- vs pre- stress; no main effect of genotype), two-way RM ANOVA. **(B–C and E–F)** miR-206 KO males (B, C) and females (E, F) are immobile in the tail suspension test for similar lengths of time as WT littermates. n = 16 WT, 10 KO males; 14 WT, 17 KO females. No effect of genotype by two-way RM ANOVA or Welch’s t-test. **(G–J)** No effect of miR-206 deletion on duration in chamber (G), entry count H), distance traveled (I), or latency to enter light chamber (J) in a 10-minute dark-light emergence task. All mice spent less time, made fewer entries into, and traveled less distance in the light chamber. n = 19 WT, 16 KO male mice. ****p < 0.0001 (duration, entry count, and distance, main effect of chamber), two-way ANOVA. Latency was analyzed by Welch’s t-test. **(K–N)** No effect of miR-206 deletion on latency to eat (K), number of sucrose pellets consumed (L), number of fecal boli (M), or distance traveled (N) in a 30-minute hyponeophagia task. n = 16 WT, 14 KO males. Data analyzed by Welch’s t-test. Error bars indicate SEM.

**Figure S7.**
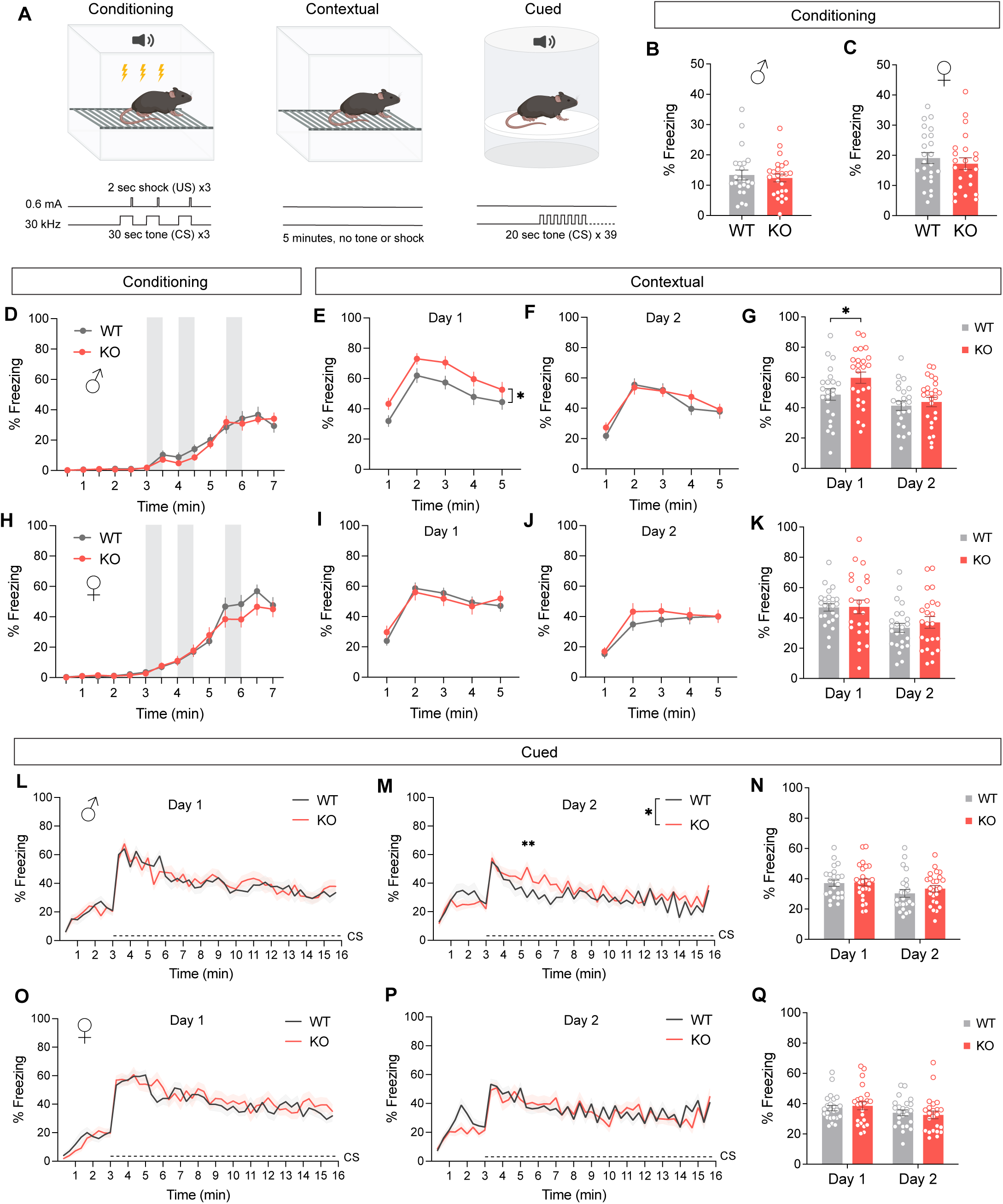
miR-206 KO male mice exhibit enhanced contextual fear memory retrieval. **(A)** Schematic of shock (US, unconditioned stimulus)- tone (CS, conditioned stimulus) pairings during fear conditioning, exposure to conditioning environment during contextual fear memory retrieval, and repeated 20-second tone presentations in a novel environment during cued fear memory retrieval. **(B, C)** Total percentage duration spent freezing during fear conditioning is unaltered in miR-206 KO males (B) and females (C). n = 23 WT, 25 KO males; 24 WT, 23 KO females (for all panels). **(D, H)** Freezing duration during fear conditioning rises similarly in miR-206 KO males (D) and females (H) as compared to WT following successive foot shocks. Gray bars indicate 30-second tone presentations with concurrent foot shock for the final 2 seconds. **(E, I)** Contextual fear memory retrieval is enhanced in miR-206 KO males (E) but not females (I). *p = 0.0422 (males, main effect of genotype). **(F, J)** Fear memory retrieval and extinction on a second day of exposure to contextual cues is not affected by miR-206 deletion in males or females. **(G, K)** Summary freezing duration across two days of contextual fear memory retrieval and extinction for males (G) and females (K). *p = 0.0495 (Day 1 WT vs KO males), Šidák’s multiple comparisons test. **(L, O)** Cued fear memory retrieval is unaltered in miR-206 KO males and females on the first day of tone presentation. **(M, P)** Extinction of cued fear memory on the second day of tone presentation is delayed in miR-206 males but not females. *p = 0.0302 (genotype x time interaction), **p = 0.0056 (WT vs KO, 320 second time point), Šidák’s multiple comparisons test. **(N, Q)** Summary freezing duration across two days of cued fear memory retrieval and extinction for males (N) and females (Q). All time-binned data were analyzed by two-way RM ANOVA. Summary data were analyzed by Welch’s t-test or two-way ANOVA. Error bars and bands indicate SEM.

**Figure S8.**
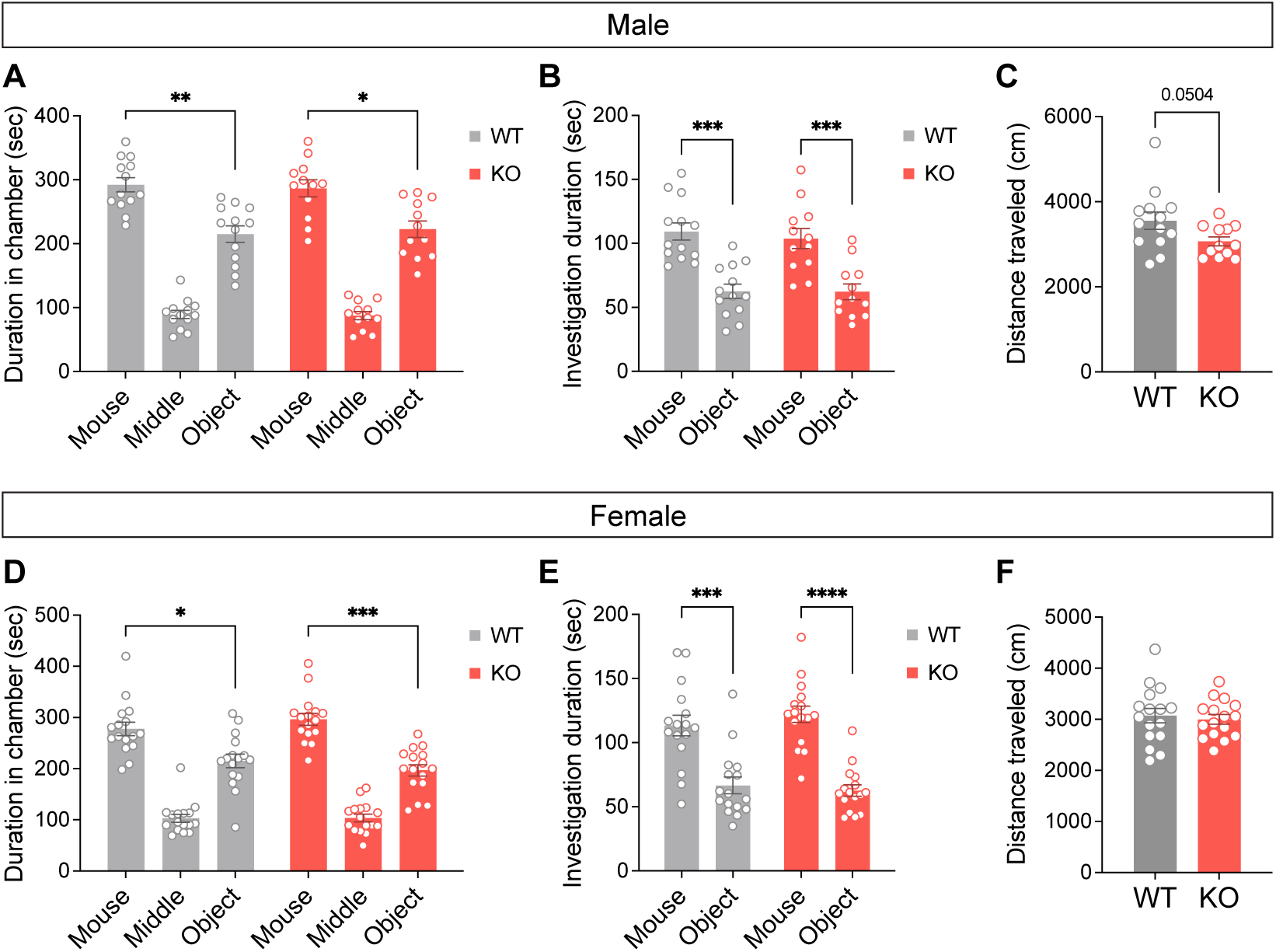
miR-206 deletion does not affect social preference in a three-chamber sociability test. **(A)** Male WT and miR-206 KO mice both spend a longer duration in a chamber containing an unfamiliar mouse than a chamber containing an empty basket. **p = 0.0058 (WT, mouse vs object), *p = 0.0283 (KO, mouse vs object). **(B)** Male WT and KO mice spend more time directly investigating the basket housing the novel mouse versus an empty basket. ***p = 0.0002 (WT, mouse vs basket), ***p = 0.0006 (KO, mouse vs basket). **(C)** Reduction in distance traveled by male KO mice did not reach significance (p = 0.0504), Welch’s t-test. **(D)** Female WT and KO mice both spend more time in the chamber paired with a novel mouse versus an empty basket. *p = 0.0273 (WT, mouse vs basket), ***p = 0.0003 (KO, mouse vs basket). **(E)** Female WT and KO mice both spend more time investigating the novel mouse-housing basket versus an empty basket. ***p = 0.0001 (WT, mouse vs basket), ****p < 0.0001 (KO, mouse vs basket). **(F)** Female WT and KO mice travel a similar distance during the test. n = 13 WT, 12 KO males; 16 WT, 16 KO females. All chamber and basket duration data were analyzed by two-way RM ANOVA followed by Holm-Šídák’s multiple comparisons test. Error bars indicate SEM.

**Figure S9.**
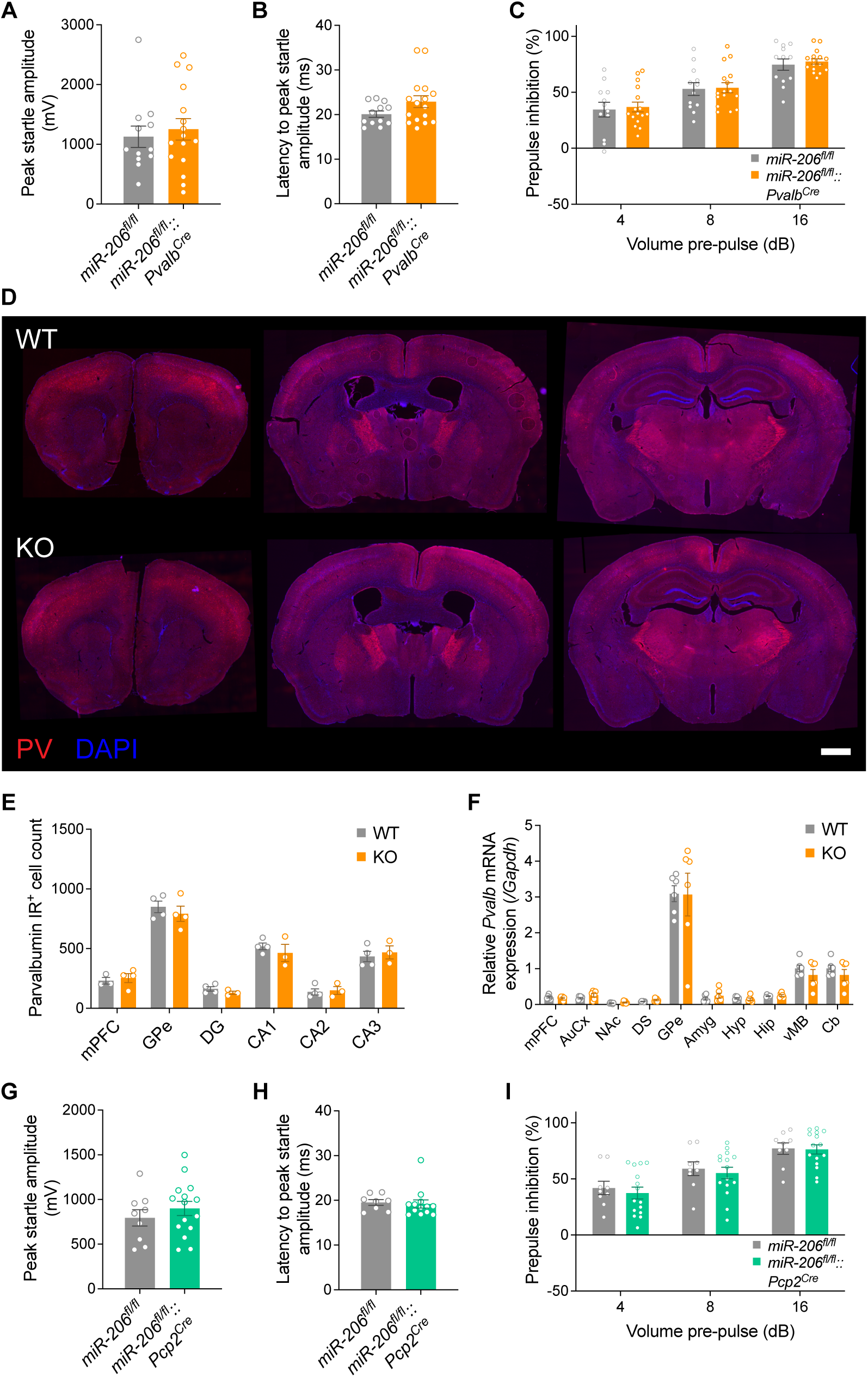
Extended sensorimotor gating and parvalbumin expression data, related to Figure 5. **(A–C)** Peak startle amplitude (A), latency to peak startle (B), and PPI (C) are unaltered in female mice with conditional deletion of miR-206 in parvalbumin-expressing neurons, including PCs and interneurons. n = 12 *miR-206^fl/fl^*, 16 *miR-206^fl/fl^*::*Pvalb^Cre^*mice. **(D)** Parvalbumin (PV) immunofluorescence and DAPI staining in representative coronal sections of WT and miR-206 KO mice. **(E)** PV immunoreactive cell counts in medial prefrontal cortex, globus pallidus external, and dentate gyrus, CA1, CA2, and CA3 hippocampal regions of WT and KO mice. n = 4 WT, 4 KO male mice. **(F)** Quantitative RT-PCR detection of *Pvalb* mRNA in brain regions of WT and miR-206 KO mice (mPFC, medial prefrontal cortex; NAc, nucleus accumbens; DS, dorsal striatum; Septum; Amyg., amygdala; Thal., thalamus; Hyp., hypothalamus; vMB, ventral midbrain; Hip, hippocampus; Cb, cerebellum). n = 6 WT, 6 KO males. **(G–I)** Peak startle amplitude (G), latency to peak startle (H), and PPI (I) are unchanged in female mice with conditional deletion of miR-206 in PCs and retinal bipolar neurons. n = 9 *miR-206^fl/fl^*, 15 *miR-206^fl/fl^*::*Pcp2^Cre^* female mice. Error bars indicate SEM.

**Figure S10.**
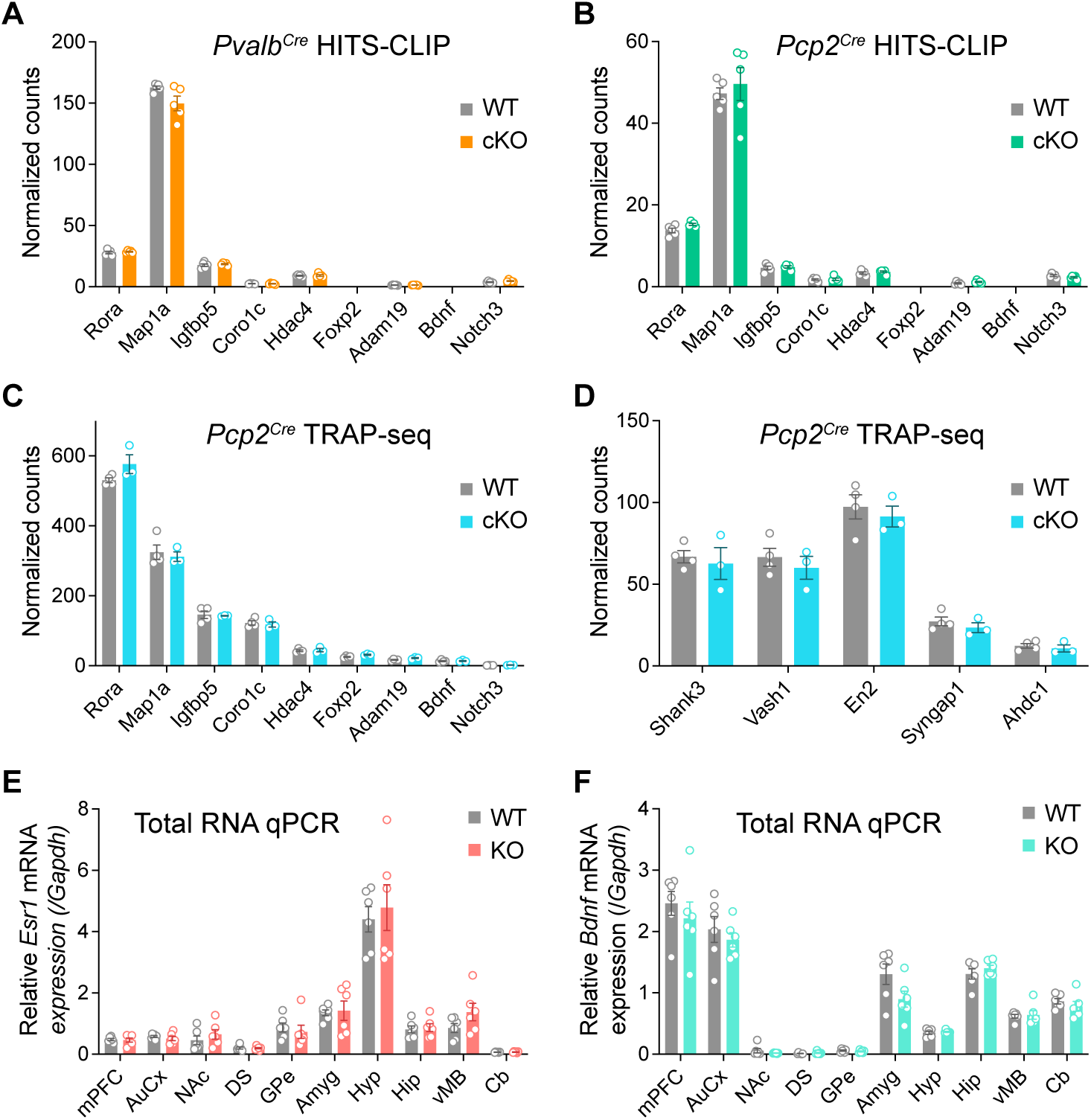
miR-206 deletion does not alter levels of previously identified miR-206 target mRNAs in the RISC or translating ribosomes of Purkinje cells, or in other profiled brain regions. **(A–C)** Normalized sequencing counts of previously identified miR-206 targets for *Pvalb^Cre^* (A) and *Pcp2^Cre^* (B) HITS-CLIP and *Pcp2^Cre^* TRAP-seq (C) experiments. n = 5 WT, 5 cKO mice (HITS-CLIP); 4 WT, 3 KO mice (TRAP). **(D)** Normalized TRAP-seq counts of targets previously identified by miR-206 sponge-mediated knockdown in PCs^28^ are unaffected by conditional miR-206 deletion in PCs. n = 4 WT, 3 cKO mice. **(E–F)** Quantitative RT-PCR detection of *Estrogen receptor 1* (E) and *Brain-derived neurotrophic factor* (F) mRNA in brain regions of WT and miR-206 KO mice. mPFC, medial prefrontal cortex; AuCx, auditory cortex; NAc, nucleus accumbens; DS, dorsal striatum; GPe, globus pallidus external; Amyg, amygdala; Hyp, hypothalamus; Hip, hippocampus; vMB, ventral midbrain; Cb, cerebellum. n = 6 WT, 6 KO males. Error bars indicate SEM.

**Supplemental Table 1.** Differential gene expression in miR-206 KO cerebellar cell types, by pseudobulk edgeR-LRT analysis of snRNA-seq data.

**Supplemental Table 2.** Differential gene expression in miR-206 KO Purkinje cell subtypes, by pseudobulk edgeR-LRT analysis of snRNA-seq data.

**Supplemental Table 3.** Differential gene expression in miR-206 KO cerebellar cortex cell types, by C-SIDE analysis of spatial transcriptomics data.

**Supplemental Table 4.** Differential gene expression in miR-206 KO cerebellar nuclei cell types, by C-SIDE analysis of spatial transcriptomics data.

**Supplemental Table 5.** Differential gene expression of AGO2-bound RNAs in *Pvalb*-expressing cerebellar neurons with miR-206 deletion, by edgeR-LRT. RNA biotype annotation from Ensembl via BioMart.

**Supplemental Table 6.** Differential gene expression of AGO2-bound RNAs in *Pcp2*-expressing cerebellar neurons with miR-206 deletion, by edgeR-LRT. RNA biotype annotation from Ensembl via BioMart.

**Supplemental Table 7.** dCLIP predictions of miR-206 binding sites in *Pvalb*-expressing cerebellar neurons.

**Supplemental Table 8.** dCLIP predictions of miR-206 binding sites in *Pcp2*-expressing cerebellar neurons.

**Supplemental Table 9.** dCLIP predictions of miR-206 binding sites shared between *Pvalb-* and *Pcp2-*expressing cerebellar neurons.

**Supplemental Table 10.** Predicted targets of miR-1/miR-206 in *Mus musculus* by TargetScan 8.0.

**Supplemental Table 11.** Differential gene expression of ribosome-bound RNAs in *Pcp2*-expressing cerebellar neurons with miR-206 deletion, by edgeR-QLF.

**Supplemental Table 12.** Summary statistics.

**Supplemental Table 13.** Oligonucleotide sequences.

